# Novel Multi-Epitope Vaccine candidate derived from critically important proteins of Monkeypox isolates

**DOI:** 10.1101/2022.09.15.507808

**Authors:** Sukrit Srivastava, Michael Kolbe

## Abstract

**Background:** On May 7, 2022, a case of monkeypox virus (MPXV) has been reported to the WHO. It causes a viral zoonotic disease with characteristics comparable to that of smallpox cases. Monkeypox could be a serious health concern as it is already spreading to multiple countries worldwide. To prevent the monkeypox infection and for its early detection, a potential vaccine and diagnostic candidate is urgently required.

**Method:** In present study we have used different in silico methods to screen the entire genome with its approximately 191 genes for potential epitopes present in proliferation and virulent proteins of monkeypox virus. All protein sequences were retrieved from different genomic and proteome databases listed in Uniprot or NCBI.

**Results:** In the present study we have screened potential epitopes from 11 different proteins of Monkeypox. All the included protein play an important role in pathogenesis and/or proliferation of Monkeypox virus. We have identified in total 984 CTL and 168 HTL epitopes with highest score in our epitope screening. The reported epitopes could be potential candidates for the design of an early detection diagnostic kit specific for the monkeypox virus. Out of these target peptides we have included a total of 39 CTL epitopes and 39 HTL epitopes in design of multi-epitope vaccine candidates. These shortlisted epitopes are highly conserved amongst different strains and origin of monkeypox viruses. The population coverage by joint administration of CTL and HTL MEVs is predicted to be high with the epitopes showing potential to bind upto 24 different CTL and HTL HLA allele molecules. The epitopes used in MEVs are examined to be highly immunogenic, non allergic but antigenic, and non toxic. All the CTL and HTL MEVs designed utilizing the epitopes have physiochemical properties favor its over expression in human cells. The optimized cDNA constructs of CTL and HTL MEVs also favor over expression of MEVs in human cells. Overall, the MEV construct proposed by us are fissile for expression in the lab and for further in vivo studies.

**Conclusion:** Control and fight against emerging diseases such as MPXV requires pathogen diagnostic and novel vaccine approaches. We screened for several epitopes and designed a MEV providing a potential solution for both purposes. Our method allows rapid screening and provides a rational strategy for the development of vaccine candidate effective in fighting MPXV and other unexpected upcoming diseases.

## INTRODUCTION

The monkeypox virus (MPXV) classified as species ‘Monkeypox virus Zaire-96-I-16’, belong to the orthopoxviruses, a genus of viruses in the family Poxviridae and subfamily Chordopoxvirinae^1,2^. The virus causes the monkeypox disease that is coupled with a rash and symptoms similar to a milder smallpox infection. The first documented monkeypox cases in the Western hemisphere was reported in the United States during 2003. The recent cases of monkeypox virus infection and its multi-country outbreak is detected outside the endemic areas of Africa in May 2022^3,4,5^. The MPXV encodes approximately 191 proteins as revealed from its genome (GCF_000857045.1_ViralProj15142_genomic)^6,7^. Out of these, at least 11 proteins seem to be important targets for vaccine design and development^8^ base upon criteria such as proliferation and cell surface binding leading to pathogenesis. The selected proteins include (sp|Q8V4S4|POXIN_MONPZ) Poxin-Schlafen OS, (sp|Q8V4Y0|CAHH_MONPZ) Cell surface-binding protein, (sp|Q8V571|P28_MONPZ) E3 ubiquitin-protein ligase p28-like, (sp|P04363|KITH_MONPZ) Thymidine kinase, (sp|Q8V4T3|SODL_MONPZ) Cu-Zn superoxide dismutase-like protein, (sp|Q8V4U9|A28_MONPZ) Envelope protein A28 homolog, (sp|Q8V4V3|RP132_MONPZ) DNA-directed RNA polymerase 132 kDa polypeptide, (sp|Q8V518|I1_MONPZ Telomere-binding protein, (sp|Q8V4T7|PROF_MONPZ) Profilin, (sp|Q8V4V4|VTF3L_MONPZ) Intermediate transcription factor 3 large subunit, (sp|Q8V566|VHR2_MONPZ) Probable host range protein 2. Here, we provide a detailed study on screening of potential CTL and HTL epitopes present in those 11 proteins. Further we also provide a multi-epitope vaccine utilizing the epitopes from all these 11 protein of Monkeypox virus.

## METHODOLOGY

### Screening of Potential Epitopes from Monkeypox virus proteins

#### T cell Epitope Prediction

##### Screening for Cytotoxic T lymphocyte (CTL) Epitope

Identification of Cytotoxic T lymphocyte epitopes was performed by the IEDB (Immune Epitope Database) tool “Proteasomal cleavage/TAP transport/MHC class I combined predictor” (http://tools.iedb.org/processing/) and “MHC-I Binding Predictions” (http://tools.iedb.org/mhci/)^9,10^. The tools provide a “Total Score” and percentile rank respectively. Further Immunogenicity of shortlisted CTL epitopes was obtained by using “MHC I Immunogenicity” tool of IEDB (http://tools.iedb.org/immunogenicity/)^11^. The physiochemical properties of constituting amino acid of the peptide sequence is utilized for immunogenicity prediction.

##### Screening for Helper T lymphocyte (HTL) Epitopes

IEDB tool “MHC-II Binding Predictions” (http://tools.iedb.org/mhcii/) was used to screen the Helper T lymphocyte (HTL) epitopes from MPXV proteins^12–15^. The tool utilizes three different methods viz. combinatorial library, SMM_align and Sturniolo for this screening^12–15^. The tool generates a percentile rank for each screened epitopes. The lower value of percentile rank indicates its higher immunogenic potential.

##### CTL and HTL epitope Toxicity prediction

The ToxinPred tool was used to characterize the toxicity of the CTL and HTL epitopes^16^. The tool use the “SVM (Swiss-Prot) based” (support vector machine) method to analyze the toxicity.

##### Conservation analysis of antigenic patches

The conservation study of the amino acid sequence of the shortlisted CTL and HTL epitopes was analyzed with the “Epitope Conservancy Analysis” tool of IEDB^17,18^.

### Multi-Epitope Vaccines against Monkeypox Virus

#### Design of Multi-Epitope Vaccines

The identified and shortlisted CTL and HTL epitopes from the monkeypox virus proteins were further used as immunogenic composition to design one CTL and one HTL Multi-Epitope Vaccine constructs as explained in the result section.

#### Physicochemical property analysis of the designed MPVs

The CTL and HTL MEVs were analyzed with the ProtParam tool to analyze their empirical properties^19^. The tool provides the empirical details of the various physicochemical parameters viz. amino acid length, molecular weight, theoretical pI, expected half-life (in *E. coli*, Yeast & Mammalian cell), Aliphatic index, Grand average of hydropathicity (GRAVY) and the instability index score. The aliphatic index and the grand average of hydropathicity (GRAVY) indicate globular and hydrophilic nature of the protein. The instability index score indicates propensity of the stability of the protein (MEVs).

#### Interferon-gamma inducing epitope prediction from the MPVs

The CTL and HTL MEVs were analyzed for potential interferon-gamma (IFN-γ) inducing epitopes (from CD4+ T cells) utilizing the “IFN epitope” tool implementing the “Motif and SVM hybrid” method i.e. MERCI: Motif-EmeRging and with Classes-Identification, and the SVM: support vector machine hybrid. This prediction is based on the IEDB database of 3705 IFN-gamma inducing and 6728 non-inducing epitopes^20,21^.

#### MPVs allergenicity and antigenicity prediction

The CTL and HTL MPVs were analyzed for allergenicity and antigenicity with the tool AllergenFP and VaxiJen respectively^22,23^. The AllergenFP prediction is based on the protein’s Hydrophobicity, size, helix-forming propensity, relative abundance of amino acids and ß-strand forming propensity. The VaxiJen tool utilizes an approach with alignment-free, based on physicochemical properties of given candidate protein sequence.

#### Analysis of MPVs cDNA for expression in human host cell line

The Codon-optimized complementary DNA (cDNA) of the CTL and HTL MEVs were generated and analyzed for the favored expression in Mammalian cell line (Human) by Java Codon Adaptation Tool. The cDNA of MEVs were further analyzed by GenScript Rare Codon Analysis Tool for large scale expression potential. The tool analyses GC content, Codon Adaptation Index (CAI) and the Tandem rare codon frequency^24,25^. The CAI score indicates the propensity of expression in the chosen human cell line expression system. The tandem rare codon frequency indicates the presence of low-frequency codons in the given cDNA.

## RESULTS & DISCUSSION

### Screening of Potential Epitopes

#### T cell Epitope Prediction

##### Screening of Cytotoxic T lymphocyte (CTL) Epitope

We found in total 984 Cytotoxic T lymphocyte (CTL) epitopes in our screening of eleven highly relevant virulente and proliferation proteins from the 191 proteins of Monkeypox virus. From those we shortlisted 39 top scoring/ranking epitope. The selected peptides were utilized for the design of MEV candidates against monkeypox virus. The immunogenicity of the shortlisted CTL epitopes was also determined (Supplementary Table S1, S2).

##### Screening of Helper T lymphocyte (HTL) epitopes

A total of 168 helper T lymphocyte (HTL) epitopes were screened from the 11 proteins of Monkeypox virus proteins. Out of these 39 screened epitope with high “Percentile rank” were shortlisted and utilized to design MEV candidate against monkeypox vaccine. The smaller the value of percentile rank indicates higher affinity of the peptide to the HLA allele binders (Supplementary Table S3).

##### CTL and HTL epitope Toxicity prediction

All the screened and shortlisted epitopes provided in Supplementary Table 1,2 & 3 are analyzed to be non-toxic.

##### Conservation analysis of the epitopes

The conservation analysis of the screened epitopes revealed that most of the epitopes are 100 % conserved amongst different strains and origin of monkeypox viruses^65^. Some of them are 60 to 90% conserved. For this analysis different protein sequences from different starins and origin of monkeypox were retrieved from NCBI protein database (Supplementary Table 1, 2 & 3).

### Multi-Epitope Vaccines against Monkeypox Virus

#### Design of Multi-Epitope Vaccines

The top scoring CTL and HTL epitopes were utilized to design one CTL and one HTL Multi-Epitope vaccines against Monkeypox virus. Short peptides GGGGS and EAAAK were used as flexible and rigid linkers respectively. The GGGGS linker provides conformational flexibility to the vaccine tertiary structure and hence facilitates conformation folding of the vaccine. The EAAAK linker facilitates in domain formation with its rigid nature and hence facilitates the vaccine to gain its final stable folded structure^26–29^.The human beta-defensin 2 (hBD-2) (PDB ID: 1FD3, Sequence: GIGDPVTCLKSGAICHPVFCPRRYKQIGTCGLPGTKCCKKP) and the human beta-defensin 3 (hBD-3) (PDB ID: 1KJ6, Sequence: GIINTLQKYYCRVRGGRCAVLSCLPKEEQIGKCSTRGRKCCRRKK) were used at N and C terminals respectively, as adjuvants to design the MEVs^26–35^. The Human Beta-defensins (hBD) have an important role in the chemotactic activity facilitating the memory T cells, immature dendritic cells and monocytes involved in the degranulation of mast cells. Due to such important role of hBDs in immune response enhancement, the hBDs have been chosen here and utilized as adjuvants for the MEV designs. The epitopes used to design CTL and HTL MEVs cover upto 24 different CTL and HTL HLA alleles. This large number of HLA alleles coverage indicates large human population coverage by the designed MEVs (Supplementary Table 4 & 5).

#### Physicochemical property analysis of designed MEVs

ProtParam calculations were performed to analyze the empirical physiochemical properties of both the CTL and HTL MEVs. The CTL MEV is composed of 661 amino acids and has molecular weight of 67.58 kDa with theoretical pI of 9.16. The expected half-life of the CTL MEV in *E.coli*, yeast and mammalian reticulocytes were predicted to be 10hrs, 20min and 30hrs respectively; the aliphatic index of the CTL MEV was 54.02 and the grand average of hydropathicity (GRAVY) was −0.227. Both the aliphatic index and the GRAVY scores indicate globular and hydrophilic nature for the CTL MEV. The instability index score of the CTL MEV was 52.65 indicating a stable protein folding under native conditions for CTL MEV. Further, the ProtParam analysis of the HTL MEV revealed it to be composed of 877 amino acids, have a molecular weight of 92.18 kDa and theoretical pI 9.70. The expected half-life of the HTL MEV in *E.coli*, yeast and mammalian reticulocytes was predicted to be 10hrs, 20min and 30hrs respectively. The aliphatic index of HTL MEV was calculated to be 88.93, and the grand average of hydropathicity (GRAVY) of the HTL MEV was 0.008, both indicate the HTL MEV to have a globular and hydrophilic nature. The instability index of the HTL MEV was 46.95 indicating its stable protein folding (Supplementary Table S6).

Overall, the physiochemical properties of both the CTL and the HTL MEVs indicate that both the vaccine candidates could be natively expressed as stable proteins in human cells and purified with standardized purifications methods and protocols.

#### Interferon-gamma inducing epitope prediction from the MPVs

The IFN-γ inducing 15 mer peptide epitopes were screened from both the CTL and HTL MEVs. A total of 74 CTL MEV and 95 HTL MEV INF-γ inducing epitopes with a score of 1 or more than 1 were shortlisted (Supplementary Table S7 & S8).

#### MEVs allergenicity and antigenicity prediction

The *allergenicity and antigenicity analysis of both the CTL and HTL MEVs revealed that both the MPVs were non allergic but antigenic in nature*. Both of these properties are essential for a vaccine candidate (Supplementary Table S9).

#### Analysis of cDNA of both the MEVs for cloning and expression in mammalian cell line

The complementary DNA codon optimized for CTL and HTL expression in mammalian host cell lines (Human) was generated. The GC content of the optimized CTL-MEV cDNA was 69.79% and a CAI (Codon Adaptation Index) score was 1.00 with 0% tandem rare codons. Likewise, the GC content of the optimized HTL-MEV cDNA was 70.69%, CAI score was 1.00 with 0% tandem rare codons. The optimum GC content of a cDNA for expression in human cell line should be within the range of 30% to 70%, the CAI score should be between 0.8-1.0, and the tandem rare codon frequency that indicates the presence of low-frequency codons, should be <30%, therefore the cDNA of both the MEVs satisfy all the mentioned parameters and are predicted to have high expression in the mammalian host cell line (Human) (Supplementary Table S10).

## DISCUSSION

Monkeypox is the causing agent of the recent outbreak in May 2022. The outbreak started in May 2022 and has quickly spread to multiple countries. A vaccine candidate against monkey pox is urgently required. We are reporting one of the first Multi-Epitope Vaccine candidates with epitopes derived from 11 different proteins of monkeypox. We have reported a total of 984 CTL and 168 HTL epitopes which have scored high in our epitope screening. Out of these we have included a total of 39 CTL epitopes and 39 HTL epitopes in the designed MEVs candidates. These epitopes are highly conserved amongst different strains and origin of monkeypox viruses. The population coverage by joint administration of CTL and HTL MEVs is expected to be high since a total of 24 different CTL and HTL HLA alleles are covered by the chosen epitopes foe MEVs. The epitopes used in MEVs are examined to be highly immunogenic, Non allergic but antigenic, and non toxic. The CTL and HTL MEVs designed utilizing the epitopes have all its physiochemical properties in favor of its over expression in vitro in human expression cells. The optimized cDNA constructs of CTL and HTL MEVs also favors over expression of MEVs in human expression cells. Overall the construct of MEVs we propose here are fissile for expression in lab for further in vivo trials. Moreover, the high scoring epitopes we have screen are also potential candidate for diagnostic kit design for early detection of the monkeypox viral infection.

## CONCLUSION

In the present study, we have identified a total of 1152 highly immunogenic novel CTL and HTL epitopes from 11 different proteins of monkeypox virus. These epitopes are examined for their conservancy, immunogenicity, toxicity, non allergic and antigenic nature. Out of these we have utilize 78 epitopes to design one CTL and one HTL MEVs. The MEVs have also shown to bind to 24 difference HLA alleles indicating large human population coverage. Both the CTL and HTL MEVs, show to favor over expression in human cells. Here we conclude that the CTL and HTL MEVs we propose here would be a potential vaccine candidate for vaccine against monkeypox virus infection. Moreover, we also conclude that the high scoreing epitopes we have screened and reported here could be a potential composition for early detection diagnostic kit design against Monkeypox virus infection.

## AUTHOR’S CONTRIBUTION

Idea conceived, Methodology designed and performed: SS; Scientific reporting and revising the manuscript: S.S. and M.K.

## ACKNOWLEDGEMENTS

We acknowledge the Indian Foundation for Fundamental Research Trust (IFFR Trust) and Helmholtz Center for Infection Research (HZI) for providing resources and funding.

## ADDITIONAL INFORMATION

Authors declare to have no competing interests.

## SUPPLEMENTARY TABLES

**Supplementary Table S1:** High Percentile Ranking CTL epitopes-HLA allele pairs screened from eleven proteins of Monkeypox virus by the “MHC-I Processing Predictions” tool of IEDB. Out of these epitopes, the top scoring epitopes were used to design Multi-epitope vaccine.

**Supplementary Table S2:** High Scoring CTL epitopes-HLA allele pairs screened from eleven proteins of Monkeypox virus by the “MHC-I Binding Predictions” tool of IEDB. Out of these epitopes, the top scoring epitopes were used to design Multi-epitope vaccine.

**Supplementary Table S3:** High Percentile Ranking HTL epitopes-HLA allele pairs screened from eleven proteins of Monkeypox virus by the “MHC-II Binding Predictions” tool of IEDB. Out of these epitopes, the top scoring epitopes were used to design Multi-epitope vaccine.

**Supplementary Table S4:** Construct of CTL-MEV and HTL-MEV.

**Supplementary Table S5:** HLA alleles covered by the CTL and HTL epitopes.

**Supplementary Table S6:** Empirical physicochemical properties of Multi-Patch Vaccine candidates.

**Supplementary Table S7:** INF-γ inducing POSITIVE epitopes, screened from the CTL MEV.

**Supplementary Table S8:** INF-γ inducing POSITIVE epitopes, screened from the HTL MPVs.

**Supplementary Table S9:** AllergenFP and Vaxijen analysis of MPVs. For the Vaxijen the default threshold is 0.4, and here all the MPVs have scored above 0.4, indicating potential antigenic nature.

**Supplementary Table S10:** Analysis of codon-optimized cDNA of all the MEVs.

## REFERENCES

1. NCBI Monkypox virus taxonomy tree: Taxonomy Browser - NCBI - NLM (nih.gov] (https://www.ncbi.nlm.nih.gov/data-hub/taxonomy/tree/?taxon=619591]

2. UNIPROT taxonomy(Taxonomy-Monkeypox virus (strain Zaire-96-I-16)): Monkeypox virus (strain Zaire-96-I-16] (MPX) | Taxonomy | UniProt (https://www.uniprot.org/taxonomy/619591]

3. Shantier, S., Mustafa, M., Abdelmoneim, A., Fadl, H., Elbager, S. and Makhawi, A., 2022. Novel Multi Epitope-based Vaccine against Monkeypox Virus: Vaccinomic approach.

4. Nalca, A., et al., Reemergence of monkeypox: prevalence, diagnostics, and countermeasures. Clin Infect Dis, 2005. 41(12): p. 1765–71.

5. Cunha, B.E., Monkeypox in the United States: an occupational health look at the first cases. Aaohn j, 2004. 52(4): p. 164–8.

6. NCBI Genome/Proteome (Genome assembly ViralProj15142): https://www.ncbi.nlm.nih.gov/data-hub/genome/GCF_000857045.1/

7. NCBI Taxonomy (Reference genome ViralProj15142; Dec 12, 2001 Strain: Zaire-96-I-16.): https://www.ncbi.nlm.nih.gov/data-hub/taxonomy/10244/

8. UNIPROT Monkeypox proteins: https://www.uniprot.org/uniprotkb?facets=reviewed%3Atrue&query=monkeypox

9. Peters B, Bulik S, Tampe R, Van Endert PM, Holzhutter HG. 2003. Identifying MHC class I epitopes by predicting the TAP transport efficiency of epitope precursors. J Immunol171:1741–1749.

10. Hoof, I., Peters, B., Sidney, J., Pedersen, L.E., Sette, A., Lund, O., Buus, S. and Nielsen, M., 2009. NetMHCpan, a method for MHC class I binding prediction beyond humans. Immunogenetics, 61(1), p.1.

11. Calis JJA, Maybeno M, Greenbaum JA, Weiskopf D, De Silva AD, Sette A, Kesmir C, Peters B. 2013. Properties of MHC class I presented peptides that enhance immunogenicity. PloS Comp. Biol. 8(1):361.

12. Wang, P., Sidney, J., Kim, Y., Sette, A., Lund, O., Nielsen, M. and Peters, B., 2010. Peptide binding predictions for HLA DR, DP and DQ molecules. BMC bioinformatics, 11(1), p.568.

13. Sidney, J., Assarsson, E., Moore, C., Ngo, S., Pinilla, C., Sette, A. and Peters, B., 2008. Quantitative peptide binding motifs for 19 human and mouse MHC class I molecules derived using positional scanning combinatorial peptide libraries. Immunome research, 4(1), p.2.

14. Nielsen, M., Lundegaard, C. and Lund, O., 2007. Prediction of MHC class II binding affinity using SMM-align, a novel stabilization matrix alignment method. BMC bioinformatics, 8(1), p.238.

15. Sturniolo, T., Bono, E., Ding, J., Raddrizzani, L., Tuereci, O., Sahin, U., Braxenthaler, M., Gallazzi, F., Protti, M.P., Sinigaglia, F. and Hammer, J., 1999. Generation of tissue-specific and promiscuous HLA ligand databases using DNA microarrays and virtual HLA class II matrices. Nature biotechnology, 17(6), p.555.

16. Gupta, S., Kapoor, P., Chaudhary, K., Gautam, A., Kumar, R., Raghava, G.P. and Open Source Drug Discovery Consortium, 2013. In silico approach for predicting toxicity of peptides and proteins. PLoS One, 8(9), p.e73957.

17. Sievers, F., Wilm, A., Dineen, D., Gibson, T.J., Karplus, K., Li, W., Lopez, R., McWilliam, H., Remmert, M., Söding, J. and Thompson, J.D., and Higgins D.G., 2011. Fast, scalable generation of high-quality protein multiple sequence alignments using Clustal Omega. Molecular systems biology, 7(1), p.539.

18. Bui HH, Sidney J, Li W, Fusseder N, Sette A. 2007. Development of an epitope conservancy analysis tool to facilitate the design of epitope-based diagnostics and vaccines. BMC Bioinformatics 8:361.

19. Gasteiger, E., Hoogland, C., Gattiker, A., Duvaud, S.E., Wilkins, M.R., Appel, R.D. and Bairoch, A., 2005. Protein identification and analysis tools on the ExPASy server (pp. 571–607). Humana Press.

20. Nagpal, G., Gupta, S., Chaudhary, K., Dhanda, S.K., Prakash, S. and Raghava, G.P., 2015. VaccineDA: Prediction, design and genome-wide screening of oligodeoxynucleotide-based vaccine adjuvants. Scientific reports, 5, p.12478.

21. Dhanda, S. K., Vir, P. & Raghava, G. P. Designing of interferon-gamma inducing MHC class-II binders. Biol. Direct. 8, 30 (2013).

22. Dimitrov I, Naneva L, Doytchinova I, Bangov I. AllergenFP: allergenicity prediction by descriptor fingerprints. Bioinformatics. 2014 Mar 15;30(6):846–51. doi: 10.1093/bioinformatics/btt619. Epub 2013 Oct 27. PMID: 24167156. https://ddg-pharmfac.net/AllergenFP/

23. Irini A Doytchinova and Darren R Flower. VaxiJen: a server for prediction of protective antigens, tumour antigens and subunit vaccines. BMC Bioinformatics. 2007 8:4. http://www.ddg-pharmfac.net/vaxijen/VaxiJen/VaxiJen.html

24. Morla, S., Makhija, A. & Kumar, S. Synonymous codon usage pattern in glycoprotein gene of rabies virus. Gene. 584, 1–6 (2016).

25. Wu, X., Wu, S., Li, D., Zhang, J., Hou, L., Ma, J., Liu, W., Ren, D., Zhu, Y. and He, F., 2010. Computational identification of rare codons of Escherichia coli based on codon pairs preference. Bmc Bioinformatics, 11(1), p.61.

26. Hu W, Li F, Yang X, Li Z, Xia H, Li G, Wang Y, Zhang Z. A flexible peptide linker enhances the immunoreactivity of two copies HBsAg preS1 (21-47) fusion protein. J Biotechnol. 2004;107:83–90.

27. Chen, X., Zaro, J.L. and Shen, W.C., 2013. Fusion protein linkers: property, design and functionality. Advanced drug delivery reviews, 65(10), pp.1357–1369.

28. Hoover, D.M., Rajashankar, K.R., Blumenthal, R., Puri, A., Oppenheim, J.J., Chertov, O. and Lubkowski, J., 2000. The structure of human ß-defensin-2 shows evidence of higher order oligomerization. Journal of Biological Chemistry, 275(42), pp.32911–32918. PDB ID: 1FD3.

29. Srivastava, S., Chatziefthymiou, S.D. and Kolbe, M., 2022. Vaccines Targeting Numerous Coronavirus Antigens, Ensuring Broader Global Population Coverage: Multi-epitope and Multi-patch Vaccines. In Vaccine Design (pp. 149–175). Humana, New York, NY.

30. Wilson, S.S., Wiens, M.E. and Smith, J.G., 2013. Antiviral mechanisms of human defensins. Journal of molecular biology, 425(24), pp.4965–4980.

31. Duits, L.A., Nibbering, P.H., Strijen, E., Vos, J.B., Mannesse-Lazeroms, S.P., Sterkenburg, M.A. and Hiemstra, P.S., 2003. Rhinovirus increases human β-defensin-2 and −3 mRNA expression in cultured bronchial epithelial cells. Pathogens and Disease, 38(1), pp.59–64.

32. Yang, D., Biragyn, A., Kwak, L.W. and Oppenheim, J.J., 2002. Mammalian defensins in immunity: more than just microbicidal. Trends in immunology, 23(6), pp.291–296.

33. Biragyn, A., Surenhu, M., Yang, D., Ruffini, P.A., Haines, B.A., Klyushnenkova, E., Oppenheim, J.J. and Kwak, L.W., 2001. Mediators of innate immunity that target immature, but not mature, dendritic cells induce antitumor immunity when genetically fused with nonimmunogenic tumor antigens. The Journal of Immunology, 167(11), pp.6644–6653.

34. Duits, L.A., Nibbering, P.H., van Strijen, E., Vos, J.B., Mannesse-Lazeroms, S.P., van Sterkenburg, M.A. and Hiemstra, P.S., 2003. Rhinovirus increases human ß-defensin-2 and-3 mRNA expression in cultured bronchial epithelial cells. FEMS Immunology & Medical Microbiology, 38(1), pp.59–64.

35. Kohlgraf, K.G., Pingel, L.C., Dietrich, D.E. and Brogden, K.A., 2010. Defensins as anti-inflammatory compounds and mucosal adjuvants. Future microbiology, 5(1), pp.99–113.

