## Supplementary for "Novel Multi-Epitope Vaccine candidate derived from critically important proteins of Monkeypox isolates"

**Supplementary Table S1:** High Percentile Ranking CTL epitopes-HLA allele pairs screened from eleven proteins of Monkeypox virus by the “MHC-I Processing Predictions” tool of IEDB. Out of these epitopes, the top scoring epitopes were used to design Multi-epitope vaccine.

| # | Allele | Start | End | Peptide Length | Peptide | Proteasome Score | TAP Score | MHC Score | Processing Score | Total Score | MHC IC50[nM] | Immunogenicity | Toxicity | Conservancy | Protein |
|---|---|---|---|---|---|---|---|---|---|---|---|---|---|---|---|
| 1 | B*58:01 | 472 | 481 | 10 | AVFADWPESW | 1.59 | 0.55 | -1.39 | 2.14 | 0.75 | 24.3 | 0.29207 | Non-Toxin | 77.78% (7/9) | Poxin-Schlafen |
| 2 | A*30:02 | 127 | 136 | 10 | AVYGKVQHEY | 1.43 | 1.54 | -1.63 | 2.96 | 1.33 | 43.1 | -0.16014 | Non-Toxin | 66.67% (6/9) | Poxin-Schlafen |
| 3 | A*26:01 | 484 | 492 | 9 | DTSGSMKKY | 1.5 | 1.14 | -1.58 | 2.64 | 1.06 | 37.8 | -0.654 | Non-Toxin | 77.78% (7/9) | Poxin-Schlafen |
| 4 | A*33:01 | 177 | 185 | 9 | EIMRMRFKR | 1.03 | 0.68 | -0.84 | 1.71 | 0.87 | 6.9 | -0.1544 | Non-Toxin | 66.67% (6/9) | Poxin-Schlafen |
| 5 | A*68:01 | 177 | 185 | 9 | EIMRMRFKR | 1.03 | 0.68 | -0.92 | 1.71 | 0.79 | 8.3 | -0.1544 | Non-Toxin | 66.67% (6/9) | Poxin-Schlafen |
| 6 | A*01:01 | 474 | 482 | 9 | FADWPESWY | 1.41 | 1.2 | -1.68 | 2.6 | 0.92 | 48.2 | 0.30334 | Non-Toxin | 77.78% (7/9) | Poxin-Schlafen |
| 7 | B*35:01 | 474 | 482 | 9 | FADWPESWY | 1.41 | 1.2 | -1.73 | 2.6 | 0.87 | 54 | 0.30334 | Non-Toxin | 77.78% (7/9) | Poxin-Schlafen |
| 8 | B*15:01 | 120 | 129 | 10 | HVLETGNAVY | 1.36 | 1.29 | -1.89 | 2.65 | 0.76 | 78 | 0.219 | Non-Toxin | 66.67% (6/9) | Poxin-Schlafen |
| 9 | B*15:01 | 425 | 433 | 9 | IQKLPVVHF | 1.37 | 1.12 | -1.59 | 2.49 | 0.89 | 39.3 | 0.00064 | Non-Toxin | 66.67% (6/9) | Poxin-Schlafen |
| 10 | A*30:02 | 89 | 98 | 10 | IVTYRHKNYY | 1.39 | 1.35 | -1.85 | 2.74 | 0.89 | 71.2 | -0.11984 | Non-Toxin | 77.78% (7/9) | Poxin-Schlafen |
| 11 | A*23:01 | 57 | 67 | 11 | KRYIGALLPMF | 1.37 | 1.38 | -1.65 | 2.75 | 1.11 | 44.2 | 0.07062 | Non-Toxin | 77.78% (7/9) | Poxin-Schlafen |
| 12 | B*15:01 | 465 | 474 | 10 | KVERCCCAVF | 1.68 | 1.15 | -1.97 | 2.83 | 0.85 | 94.2 | -0.01403 | Non-Toxin | 77.78% (7/9) | Poxin-Schlafen |
| 13 | A*30:02 | 441 | 450 | 10 | KYACRFIKVY | 1.64 | 1.44 | -2.22 | 3.08 | 0.87 | 164.9 | 0.09025 | Non-Toxin | 77.78% (7/9) | Poxin-Schlafen |
| 14 | B*35:01 | 63 | 72 | 10 | LLPMFECNEY | 1.18 | 1.34 | -1.41 | 2.52 | 1.11 | 25.6 | 0.03349 | Non-Toxin | 77.78% (7/9) | Poxin-Schlafen |
| 15 | B*35:01 | 64 | 72 | 9 | LPMFECNEY | 1.18 | 1.19 | -0.59 | 2.36 | 1.77 | 3.9 | 0.16059 | Non-Toxin | 77.78% (7/9) | Poxin-Schlafen |
| 16 | B*53:01 | 64 | 72 | 9 | LPMFECNEY | 1.18 | 1.19 | -1.59 | 2.36 | 0.77 | 39.2 | 0.16059 | Non-Toxin | 77.78% (7/9) | Poxin-Schlafen |
| 17 | B*15:01 | 346 | 354 | 9 | LQAGESVKF | 1.46 | 1.07 | -1.75 | 2.53 | 0.78 | 55.7 | -0.10259 | Non-Toxin | 88.89% (8/9) | Poxin-Schlafen |
| 18 | A*02:06 | 292 | 300 | 9 | LQMDSMEAL | 1.6 | 0.48 | -0.69 | 2.08 | 1.39 | 4.9 | -0.25372 | Non-Toxin | 66.67% (6/9) | Poxin-Schlafen |
| 19 | A*30:02 | 1 | 10 | 10 | MFYAHAFGGY | 1.2 | 1.38 | -1.61 | 2.58 | 0.97 | 40.7 | 0.26637 | Non-Toxin | 77.78% (7/9) | Poxin-Schlafen |
| 20 | A*01:01 | 293 | 302 | 10 | QMDSMEALEY | 1.17 | 1.27 | -1.62 | 2.45 | 0.83 | 41.3 | -0.1468 | Non-Toxin | 88.89% (8/9) | Poxin-Schlafen |
| 21 | B*15:01 | 376 | 385 | 10 | QQLPSILSSF | 1.33 | 1.2 | -1.3 | 2.53 | 1.23 | 19.8 | -0.29298 | Non-Toxin | 100.00% (9/9) | Poxin-Schlafen |
| 22 | A*30:02 | 29 | 37 | 9 | RKYSVVSVY | 1.28 | 1.5 | -2.03 | 2.78 | 0.75 | 106.9 | -0.20411 | Non-Toxin | 77.78% (7/9) | Poxin-Schlafen |
| 23 | A*23:01 | 58 | 67 | 10 | RYIGALLPMF | 1.37 | 1.39 | -0.73 | 2.77 | 2.04 | 5.4 | -0.0178 | Non-Toxin | 77.78% (7/9) | Poxin-Schlafen |
| 24 | A*24:02 | 58 | 67 | 10 | RYIGALLPMF | 1.37 | 1.39 | -0.94 | 2.77 | 1.83 | 8.7 | -0.0178 | Non-Toxin | 77.78% (7/9) | Poxin-Schlafen |
| 25 | B*15:01 | 242 | 251 | 10 | SIHHLWSVVY | 1.71 | 1.29 | -1.83 | 3 | 1.17 | 67.8 | 0.15118 | Non-Toxin | 66.67% (6/9) | Poxin-Schlafen |
| 26 | A*30:02 | 242 | 251 | 10 | SIHHLWSVVY | 1.71 | 1.29 | -2.05 | 3 | 0.95 | 112.3 | 0.15118 | Non-Toxin | 66.67% (6/9) | Poxin-Schlafen |
| 27 | A*30:02 | 383 | 392 | 10 | SSFANTKGGY | 1.07 | 1.38 | -1.39 | 2.45 | 1.06 | 24.4 | -0.04573 | Non-Toxin | 88.89% (8/9) | Poxin-Schlafen |
| 28 | A*30:02 | 90 | 98 | 9 | VTYRHKNYY | 1.39 | 1.37 | -1.32 | 2.76 | 1.45 | 20.7 | -0.12824 | Non-Toxin | 77.78% (7/9) | Poxin-Schlafen |
| 29 | A*02:03 | 249 | 257 | 9 | VVYDHLNVV | 1.36 | 0.33 | -0.78 | 1.69 | 0.91 | 6 | 0.06084 | Non-Toxin | 66.67% (6/9) | Poxin-Schlafen |
| 30 | B*35:01 | 442 | 450 | 9 | YACRFIKVY | 1.64 | 1.33 | -2.11 | 2.97 | 0.87 | 127.8 | 0.11598 | Non-Toxin | 77.78% (7/9) | Poxin-Schlafen |
| 31 | B*15:01 | 424 | 433 | 10 | YIQKLPVVHF | 1.37 | 1.18 | -1.66 | 2.54 | 0.88 | 46.1 | -0.1836 | Non-Toxin | 66.67% (6/9) | Poxin-Schlafen |
| 32 | A*68:01 | 460 | 468 | 9 | YVCAIKVER | 1.07 | 0.7 | -0.98 | 1.78 | 0.8 | 9.5 | 0.04181 | Non-Toxin | 77.78% (7/9) | Poxin-Schlafen |
| 33 | A*02:03 | 234 | 243 | 10 | YVLMKRLESI | 1.24 | 0.23 | -0.73 | 1.47 | 0.74 | 5.4 | -0.3625 | Non-Toxin | 77.78% (7/9) | Poxin-Schlafen |
| **#** | **Allele** | **Start** | **End** | **Peptide Length** | **Peptide** | **Proteasome Score** | **TAP Score** | **MHC Score** | **Processing Score** | **Total Score** | **MHC IC50[nM]** | **Immunogenicity** | **Toxicity** | Conservancy | **Protein** |
| 34 | B*15:01 | 224 | 232 | 9 | KLNDDTQVY | 1.62 | 1.26 | -1.78 | 2.88 | 1.1 | 59.6 | 0.00472 | Non-Toxin | 100.00% (7/7) | Cell surface-binding protein |
| 35 | A*32:01 | 274 | 283 | 10 | KTFAIIAIVF | 1.52 | 1.25 | -1.23 | 2.76 | 1.53 | 17 | 0.50984 | Non-Toxin | 100.00% (7/7) | Cell surface-binding protein |
| 36 | B*58:01 | 274 | 283 | 10 | KTFAIIAIVF | 1.52 | 1.25 | -1.4 | 2.76 | 1.36 | 25 | 0.50984 | Non-Toxin | 100.00% (7/7) | Cell surface-binding protein |
| 37 | B*57:01 | 274 | 283 | 10 | KTFAIIAIVF | 1.52 | 1.25 | -1.75 | 2.76 | 1.01 | 56 | 0.50984 | Non-Toxin | 100.00% (7/7) | Cell surface-binding protein |
| 38 | A*23:01 | 268 | 276 | 9 | KYIEGNKTF | 1.77 | 1.27 | -1.46 | 3.04 | 1.58 | 29 | 0.01154 | Non-Toxin | 100.00% (7/7) | Cell surface-binding protein |
| 39 | A*24:02 | 268 | 276 | 9 | KYIEGNKTF | 1.77 | 1.27 | -1.56 | 3.04 | 1.48 | 36 | 0.01154 | Non-Toxin | 100.00% (7/7) | Cell surface-binding protein |
| 40 | B*35:01 | 160 | 169 | 10 | LPSTLDYFTY | 1.32 | 1.16 | -1.03 | 2.47 | 1.45 | 10.6 | 0.11416 | Non-Toxin | 100.00% (7/7) | Cell surface-binding protein |
| 41 | B*15:01 | 121 | 130 | 10 | LQVSDHKNVY | 1.5 | 1.29 | -1.43 | 2.79 | 1.36 | 27.2 | -0.28431 | Non-Toxin | 100.00% (7/7) | Cell surface-binding protein |
| 42 | B*58:01 | 146 | 154 | 9 | MSAPFDSVF | 1.46 | 1.21 | -0.93 | 2.66 | 1.74 | 8.5 | 0.02092 | Non-Toxin | 100.00% (7/7) | Cell surface-binding protein |
| 43 | B*35:01 | 146 | 154 | 9 | MSAPFDSVF | 1.46 | 1.21 | -1.07 | 2.66 | 1.59 | 11.8 | 0.02092 | Non-Toxin | 100.00% (7/7) | Cell surface-binding protein |
| 44 | B*15:01 | 146 | 154 | 9 | MSAPFDSVF | 1.46 | 1.21 | -1.2 | 2.66 | 1.47 | 15.8 | 0.02092 | Non-Toxin | 100.00% (7/7) | Cell surface-binding protein |
| 45 | A*30:02 | 146 | 155 | 10 | MSAPFDSVFY | 1.56 | 1.38 | -1.33 | 2.93 | 1.6 | 21.6 | 0.08465 | Non-Toxin | 100.00% (7/7) | Cell surface-binding protein |
| 46 | B*58:01 | 146 | 155 | 10 | MSAPFDSVFY | 1.56 | 1.38 | -1.34 | 2.93 | 1.59 | 22 | 0.08465 | Non-Toxin | 100.00% (7/7) | Cell surface-binding protein |
| 47 | A*01:01 | 146 | 155 | 10 | MSAPFDSVFY | 1.56 | 1.38 | -1.39 | 2.93 | 1.55 | 24.3 | 0.08465 | Non-Toxin | 100.00% (7/7) | Cell surface-binding protein |

| | | | | | | | | | | | | | | | |
|---|---|---|---|---|---|---|---|---|---|---|---|---|---|---|---|
| 48 | B*15:01 | 146 | 155 | 10 | MSAPFDSVFY | 1.56 | 1.38 | -1.44 | 2.93 | 1.5 | 27.3 | 0.08465 | Non-Toxin | 100.00% (7/7) | Cell surface-binding protein |
| 49 | A*68:01 | 146 | 155 | 10 | MSAPFDSVFY | 1.56 | 1.38 | -1.49 | 2.93 | 1.44 | 31 | 0.08465 | Non-Toxin | 100.00% (7/7) | Cell surface-binding protein |
| 50 | A*11:01 | 146 | 155 | 10 | MSAPFDSVFY | 1.56 | 1.38 | -1.7 | 2.93 | 1.24 | 49.7 | 0.08465 | Non-Toxin | 100.00% (7/7) | Cell surface-binding protein |
| 51 | B*35:01 | 146 | 155 | 10 | MSAPFDSVFY | 1.56 | 1.38 | -1.77 | 2.93 | 1.16 | 59.1 | 0.08465 | Non-Toxin | 100.00% (7/7) | Cell surface-binding protein |
| 52 | B*15:01 | 145 | 154 | 10 | NMSAPFDSVF | 1.46 | 1.23 | -1.32 | 2.69 | 1.37 | 20.8 | -0.00575 | Non-Toxin | 100.00% (7/7) | Cell surface-binding protein |
| 53 | B*15:01 | 20 | 28 | 9 | RLKTLDIHY | 1.49 | 1.39 | -1.53 | 2.88 | 1.35 | 34.2 | 0.11036 | Non-Toxin | 100.00% (7/7) | Cell surface-binding protein |
| 54 | A*30:02 | 20 | 28 | 9 | RLKTLDIHY | 1.49 | 1.39 | -1.88 | 2.88 | 1 | 75.6 | 0.11036 | Non-Toxin | 100.00% (7/7) | Cell surface-binding protein |
| 55 | A*02:06 | 162 | 170 | 9 | STLDYFTYL | 1.53 | 0.37 | -0.69 | 1.91 | 1.22 | 4.9 | 0.15592 | Non-Toxin | 100.00% (7/7) | Cell surface-binding protein |
| 56 | B*58:01 | 87 | 96 | 10 | YSGEINLVHW | 1.51 | 0.37 | -0.87 | 1.87 | 1 | 7.4 | 0.27835 | Non-Toxin | 100.00% (7/7) | Cell surface-binding protein |
| 57 | B*35:01 | 61 | 69 | 9 | YVLSTIHIY | 1.28 | 1.26 | -1.13 | 2.54 | 1.41 | 13.4 | 0.09807 | Non-Toxin | 100.00% (7/7) | Cell surface-binding protein |
| **#** | **Allele** | **Start** | **End** | **Peptide Length** | **Peptide** | **Proteasome Score** | **TAP Score** | **MHC Score** | **Processing Score** | **Total Score** | **MHC IC50[nM]** | **Immunogenicity** | **Toxicity** | Conservancy | **Protein** |
| 58 | A*31:01 | 205 | 213 | 9 | CINIWHRTR | 1.2 | 0.65 | -1.34 | 1.85 | 0.51 | 22 | 0.44433 | Non-Toxin | 100.00% (19/19) | E3 ubiquitin-protein ligase |
| 59 | A*26:01 | 153 | 161 | 9 | DIIKILDKY | 1.36 | 1.24 | -1.5 | 2.6 | 1.11 | 31.4 | -0.16192 | Non-Toxin | 100.00% (19/19) | E3 ubiquitin-protein ligase |
| 60 | B*15:01 | 34 | 43 | 10 | IIRNINNITY | 1.77 | 1.35 | -1.98 | 3.12 | 1.14 | 95.6 | 0.2625 | Non-Toxin | 52.63% (10/19) | E3 ubiquitin-protein ligase |
| 61 | A*30:02 | 34 | 43 | 10 | IIRNINNITY | 1.77 | 1.35 | -2.18 | 3.12 | 0.94 | 152.8 | 0.2625 | Non-Toxin | 52.63% (10/19) | E3 ubiquitin-protein ligase |
| 62 | A*01:01 | 157 | 165 | 9 | ILDKYEDMY | 1.32 | 1.26 | -1.69 | 2.58 | 0.89 | 48.6 | -0.20303 | Non-Toxin | 100.00% (19/19) | E3 ubiquitin-protein ligase |
| 63 | B*15:01 | 124 | 132 | 9 | ILRGLVNWY | 1.28 | 1.33 | -2.21 | 2.61 | 0.4 | 161.9 | 0.20292 | Non-Toxin | 100.00% (19/19) | E3 ubiquitin-protein ligase |
| 64 | A*23:01 | 98 | 107 | 10 | IYGLYIHYNL | 1.67 | 0.53 | -1.35 | 2.2 | 0.86 | 22.3 | 0.14939 | Non-Toxin | 100.00% (19/19) | E3 ubiquitin-protein ligase |
| 65 | A*24:02 | 98 | 107 | 10 | IYGLYIHYNL | 1.67 | 0.53 | -1.63 | 2.2 | 0.57 | 42.6 | 0.14939 | Non-Toxin | 100.00% (19/19) | E3 ubiquitin-protein ligase |
| 66 | B*44:03 | 169 | 177 | 9 | KEKECGICY | 1.41 | 1.28 | -2.28 | 2.69 | 0.41 | 189.6 | 0.09097 | Non-Toxin | 100.00% (19/19) | E3 ubiquitin-protein ligase |
| 67 | B*44:02 | 169 | 177 | 9 | KEKECGICY | 1.41 | 1.28 | -2.28 | 2.69 | 0.41 | 191.1 | 0.09097 | Non-Toxin | 100.00% (19/19) | E3 ubiquitin-protein ligase |
| 68 | A*31:01 | 49 | 57 | 9 | KINPHLANR | 0.92 | 0.71 | -0.82 | 1.63 | 0.81 | 6.6 | 0.03704 | Non-Toxin | 100.00% (19/19) | E3 ubiquitin-protein ligase |
| 69 | A*31:01 | 49 | 58 | 10 | KINPHLANRF | 1.51 | 1.17 | -2.09 | 2.68 | 0.59 | 121.8 | 0.06905 | Non-Toxin | 100.00% (19/19) | E3 ubiquitin-protein ligase |
| 70 | B*15:01 | 49 | 58 | 10 | KINPHLANRF | 1.51 | 1.17 | -2.17 | 2.68 | 0.51 | 146.3 | 0.06905 | Non-Toxin | 100.00% (19/19) | E3 ubiquitin-protein ligase |
| 71 | A*31:01 | 49 | 59 | 11 | KINPHLANRFR | 1.18 | 0.71 | -1.32 | 1.89 | 0.56 | 21.1 | 0.15153 | Non-Toxin | 100.00% (19/19) | E3 ubiquitin-protein ligase |
| 72 | A*03:01 | 96 | 105 | 10 | KNIYGLYIHY | 1.59 | 1.34 | -1.84 | 2.94 | 1.1 | 69.2 | 0.18942 | Non-Toxin | 100.00% (19/19) | E3 ubiquitin-protein ligase |
| 73 | A*30:02 | 96 | 105 | 10 | KNIYGLYIHY | 1.59 | 1.34 | -1.86 | 2.94 | 1.07 | 73.1 | 0.18942 | Non-Toxin | 100.00% (19/19) | E3 ubiquitin-protein ligase |
| 74 | B*15:01 | 96 | 105 | 10 | KNIYGLYIHY | 1.59 | 1.34 | -2.38 | 2.94 | 0.56 | 237.6 | 0.18942 | Non-Toxin | 100.00% (19/19) | E3 ubiquitin-protein ligase |
| 75 | A*11:01 | 96 | 105 | 10 | KNIYGLYIHY | 1.59 | 1.34 | -2.41 | 2.94 | 0.53 | 255.2 | 0.18942 | Non-Toxin | 100.00% (19/19) | E3 ubiquitin-protein ligase |
| 76 | A*30:02 | 94 | 102 | 9 | KNKNIYGLY | 1.16 | 1.23 | -1.39 | 2.39 | 0.99 | 24.8 | 0.07173 | Non-Toxin | 100.00% (19/19) | E3 ubiquitin-protein ligase |
| 77 | A*68:02 | 37 | 45 | 9 | NINNITYYI | 1.27 | 0.32 | -0.83 | 1.59 | 0.76 | 6.7 | 0.15225 | Non-Toxin | 52.63% (10/19) | E3 ubiquitin-protein ligase |
| 78 | A*68:01 | 97 | 105 | 9 | NIYGLYIHY | 1.59 | 1.47 | -2.28 | 3.06 | 0.78 | 191.2 | 0.14984 | Non-Toxin | 100.00% (19/19) | E3 ubiquitin-protein ligase |
| 79 | B*35:01 | 97 | 105 | 9 | NIYGLYIHY | 1.59 | 1.47 | -2.58 | 3.06 | 0.48 | 379.6 | 0.14984 | Non-Toxin | 100.00% (19/19) | E3 ubiquitin-protein ligase |
| 80 | A*30:02 | 97 | 105 | 9 | NIYGLYIHY | 1.59 | 1.47 | -2.59 | 3.06 | 0.47 | 393 | 0.14984 | Non-Toxin | 100.00% (19/19) | E3 ubiquitin-protein ligase |
| 81 | B*15:01 | 97 | 105 | 9 | NIYGLYIHY | 1.59 | 1.47 | -2.61 | 3.06 | 0.45 | 404.7 | 0.14984 | Non-Toxin | 100.00% (19/19) | E3 ubiquitin-protein ligase |
| 82 | B*58:01 | 122 | 131 | 10 | QSILRGLVNW | 1.74 | 0.46 | -1.16 | 2.19 | 1.03 | 14.6 | 0.13606 | Non-Toxin | 100.00% (19/19) | E3 ubiquitin-protein ligase |
| 83 | B*57:01 | 122 | 131 | 10 | QSILRGLVNW | 1.74 | 0.46 | -1.49 | 2.19 | 0.71 | 30.6 | 0.13606 | Non-Toxin | 100.00% (19/19) | E3 ubiquitin-protein ligase |
| 84 | A*31:01 | 59 | 68 | 10 | RAWKKRIAGR | 0.79 | 0.8 | -1.15 | 1.59 | 0.44 | 14.2 | -0.1266 | Non-Toxin | 100.00% (19/19) | E3 ubiquitin-protein ligase |
| 85 | A*30:01 | 228 | 236 | 9 | RFRKITMSK | 0.72 | 0.41 | -0.53 | 1.12 | 0.59 | 3.4 | -0.27892 | Non-Toxin | 73.68% (14/19) | E3 ubiquitin-protein ligase |
| 86 | A*30:01 | 228 | 237 | 10 | RFRKITMSKF | 1.24 | 1.3 | -1.56 | 2.54 | 0.98 | 36 | -0.46372 | Non-Toxin | 73.68% (14/19) | E3 ubiquitin-protein ligase |
| 87 | A*23:01 | 228 | 237 | 10 | RFRKITMSKF | 1.24 | 1.3 | -1.94 | 2.54 | 0.6 | 87.3 | -0.46372 | Non-Toxin | 73.68% (14/19) | E3 ubiquitin-protein ligase |
| 88 | A*24:02 | 228 | 237 | 10 | RFRKITMSKF | 1.24 | 1.3 | -2.12 | 2.54 | 0.42 | 132.9 | -0.46372 | Non-Toxin | 73.68% (14/19) | E3 ubiquitin-protein ligase |
| 89 | A*30:02 | 228 | 238 | 11 | RFRKITMSKFY | 1.56 | 1.47 | -2.16 | 3.04 | 0.87 | 145 | -0.47313 | Non-Toxin | 73.68% (14/19) | E3 ubiquitin-protein ligase |
| 90 | A*30:01 | 228 | 238 | 11 | RFRKITMSKFY | 1.56 | 1.47 | -2.31 | 3.04 | 0.72 | 204.8 | -0.47313 | Non-Toxin | 73.68% (14/19) | E3 ubiquitin-protein ligase |
| 91 | A*30:02 | 36 | 44 | 9 | RNINNITYY | 1.14 | 1.34 | -1.61 | 2.48 | 0.88 | 40.4 | 0.18627 | Non-Toxin | 52.63% (10/19) | E3 ubiquitin-protein ligase |
| 92 | A*30:01 | 226 | 234 | 9 | RTRFRKITM | 1.09 | 0.25 | -0.76 | 1.33 | 0.58 | 5.7 | 0.117 | Non-Toxin | 73.68% (14/19) | E3 ubiquitin-protein ligase |
| 93 | B*07:02 | 211 | 219 | 9 | RTRRETGAL | 1.3 | 0.52 | -1.25 | 1.83 | 0.58 | 17.7 | 0.25438 | Non-Toxin | 47.37% (9/19) | E3 ubiquitin-protein ligase |
| 94 | A*30:01 | 211 | 219 | 9 | RTRRETGAL | 1.3 | 0.52 | -1.31 | 1.83 | 0.52 | 20.5 | 0.25438 | Non-Toxin | 47.37% (9/19) | E3 ubiquitin-protein ligase |
| 95 | B*40:01 | 150 | 158 | 9 | SELDIIKIL | 1.53 | 0.39 | -1.44 | 1.92 | 0.49 | 27.3 | 0.16936 | Non-Toxin | 100.00% (19/19) | E3 ubiquitin-protein ligase |
| 96 | A*01:01 | 12 | 21 | 10 | SIDHVTILQY | 1.45 | 1.3 | -1.73 | 2.75 | 1.02 | 53.2 | 0.16599 | Non-Toxin | 100.00% (19/19) | E3 ubiquitin-protein ligase |
| 97 | A*30:02 | 129 | 138 | 10 | VNWYIANNTY | 1.51 | 1.33 | -2.26 | 2.83 | 0.58 | 181.2 | 0.24701 | Non-Toxin | 100.00% (19/19) | E3 ubiquitin-protein ligase |
| **#** | **Allele** | **Start** | **End** | **Peptide Length** | **Peptide** | **Proteasome Score** | **TAP Score** | **MHC Score** | **Processing Score** | **Total Score** | **MHC IC50[nM]** | **Immunogenicity** | **Toxicity** | Conservancy | **Protein** |

| | | | | | | | | | | | | | | | |
|---|---|---|---|---|---|---|---|---|---|---|---|---|---|---|---|
| 98 | B*15:01 | 31 | 40 | 10 | AQYKCVTIKY | 1.47 | 1.45 | -1.38 | 2.92 | 1.55 | 23.8 | -0.20764 | Non-Toxin | 100.00% (607/607) | Thymidine kinase |
| 99 | A*30:02 | 31 | 40 | 10 | AQYKCVTIKY | 1.47 | 1.45 | -1.96 | 2.92 | 0.96 | 91.2 | -0.20764 | Non-Toxin | 100.00% (607/607) | Thymidine kinase |
| 100 | B*15:01 | 138 | 147 | 10 | CMKCFKEASF | 1.13 | 1.15 | -1.85 | 2.28 | 0.44 | 70.2 | -0.18964 | Non-Toxin | 100.00% (607/607) | Thymidine kinase |
| 101 | A*26:01 | 158 | 166 | 9 | EIIGGNDMY | 1.24 | 1.27 | -1.37 | 2.52 | 1.14 | 23.7 | 0.02033 | Non-Toxin | 100.00% (607/607) | Thymidine kinase |
| 102 | B*35:01 | 158 | 166 | 9 | EIIGGNDMY | 1.24 | 1.27 | -2.15 | 2.52 | 0.36 | 141.7 | 0.02033 | Non-Toxin | 100.00% (607/607) | Thymidine kinase |
| 103 | A*26:01 | 20 | 28 | 9 | ELIRRVRRY | 1.48 | 1.22 | -1.76 | 2.7 | 0.94 | 58.2 | 0.25846 | Non-Toxin | 99.84% (606/607) | Thymidine kinase |
| 104 | B*15:01 | 84 | 93 | 10 | GQFFPDIVEF | 1.41 | 1.03 | -1.62 | 2.44 | 0.82 | 41.9 | 0.38522 | Non-Toxin | 99.67% (605/607) | Thymidine kinase |
| 105 | A*02:03 | 125 | 134 | 10 | IPLSEMVVKL | 1.57 | 0.32 | -1.61 | 1.89 | 0.29 | 40.4 | -0.29587 | Non-Toxin | 100.00% (607/607) | Thymidine kinase |
| 106 | B*15:01 | 105 | 113 | 9 | IVAALDGTF | 1.12 | 1.2 | -1.54 | 2.32 | 0.78 | 34.6 | 0.11343 | Non-Toxin | 100.00% (607/607) | Thymidine kinase |
| 107 | B*35:01 | 105 | 113 | 9 | IVAALDGTF | 1.12 | 1.2 | -2.06 | 2.32 | 0.26 | 115.1 | 0.11343 | Non-Toxin | 100.00% (607/607) | Thymidine kinase |
| 108 | A*02:03 | 65 | 73 | 9 | KLCDVLEAI | 1.36 | 0.29 | -1.14 | 1.66 | 0.51 | 13.9 | 0.14194 | Non-Toxin | 99.84% (606/607) | Thymidine kinase |
| 109 | A*02:01 | 65 | 73 | 9 | KLCDVLEAI | 1.36 | 0.29 | -1.21 | 1.66 | 0.45 | 16.2 | 0.14194 | Non-Toxin | 99.84% (606/607) | Thymidine kinase |
| 110 | A*02:06 | 65 | 73 | 9 | KLCDVLEAI | 1.36 | 0.29 | -1.22 | 1.66 | 0.44 | 16.5 | 0.14194 | Non-Toxin | 99.84% (606/607) | Thymidine kinase |
| 111 | A*02:03 | 13 | 21 | 9 | MLSGKSTEL | 1.21 | 0.38 | -1.06 | 1.59 | 0.53 | 11.6 | -0.29407 | Non-Toxin | 0.00% (0/607) | Thymidine kinase |
| 112 | B*15:01 | 165 | 174 | 10 | MYQSVCRKCY | 1.52 | 1.33 | -2.42 | 2.85 | 0.44 | 260.3 | -0.38115 | Non-Toxin | 100.00% (607/607) | Thymidine kinase |
| 113 | A*30:02 | 165 | 174 | 10 | MYQSVCRKCY | 1.52 | 1.33 | -2.53 | 2.85 | 0.32 | 338 | -0.38115 | Non-Toxin | 100.00% (607/607) | Thymidine kinase |
| 114 | A*23:01 | 85 | 93 | 9 | QFFPDIVEF | 1.41 | 1.21 | -1.64 | 2.63 | 0.98 | 43.8 | 0.26706 | Non-Toxin | 99.67% (605/607) | Thymidine kinase |
| 115 | A*24:02 | 85 | 93 | 9 | QFFPDIVEF | 1.41 | 1.21 | -2.26 | 2.63 | 0.36 | 182.3 | 0.26706 | Non-Toxin | 99.67% (605/607) | Thymidine kinase |
| 116 | B*07:02 | 116 | 124 | 9 | RPFNNILNL | 1.44 | 0.5 | -1.48 | 1.93 | 0.46 | 29.9 | 0.13733 | Non-Toxin | 99.67% (605/607) | Thymidine kinase |
| 117 | A*30:02 | 24 | 33 | 10 | RVRRYQIAQY | 1.3 | 1.51 | -1.81 | 2.81 | 1.01 | 64 | 0.0431 | Non-Toxin | 100.00% (607/607) | Thymidine kinase |
| 118 | A*30:01 | 24 | 33 | 10 | RVRRYQIAQY | 1.3 | 1.51 | -1.93 | 2.81 | 0.89 | 84.2 | 0.0431 | Non-Toxin | 100.00% (607/607) | Thymidine kinase |
| 119 | B*15:01 | 24 | 33 | 10 | RVRRYQIAQY | 1.3 | 1.51 | -2.23 | 2.81 | 0.58 | 171.1 | 0.0431 | Non-Toxin | 100.00% (607/607) | Thymidine kinase |
| 120 | A*31:01 | 18 | 27 | 10 | STELIRRVRR | 1.24 | 0.64 | -1.58 | 1.87 | 0.3 | 37.7 | 0.31514 | Non-Toxin | 99.84% (606/607) | Thymidine kinase |
| 121 | A*68:01 | 37 | 45 | 9 | TIKYSNDNR | 1.02 | 0.71 | -1.38 | 1.73 | 0.35 | 23.9 | -0.22597 | Non-Toxin | 100.00% (607/607) | Thymidine kinase |
| 122 | B*15:01 | 104 | 113 | 10 | VIVAALDGTF | 1.12 | 1.22 | -1.72 | 2.34 | 0.62 | 52.2 | 0.15223 | Non-Toxin | 100.00% (607/607) | Thymidine kinase |
| 123 | B*15:01 | 166 | 174 | 9 | YQSVCRKCY | 1.52 | 1.31 | -2.03 | 2.83 | 0.8 | 106.5 | -0.22944 | Non-Toxin | 100.00% (607/607) | Thymidine kinase |
| 124 | A*30:02 | 166 | 174 | 9 | YQSVCRKCY | 1.52 | 1.31 | -2.55 | 2.83 | 0.28 | 357.7 | -0.22944 | Non-Toxin | 100.00% (607/607) | Thymidine kinase |

| # | Allele | Start | End | Peptide Length | Peptide | Proteasome Score | TAP Score | MHC Score | Processing Score | Total Score | MHC IC50[nM] | Immunogenicity | Toxicity | Conservancy | Protein |
|---|---|---|---|---|---|---|---|---|---|---|---|---|---|---|---|
| 125 | B*35:01 | 66 | 74 | 9 | FVNRYGVAY | 1.49 | 1.25 | -0.9 | 2.74 | 1.84 | 8 | 0.13598 | Non-Toxin | 100.00% (86/86) | Cu-Zn superoxide dismutase-like protein |
| 126 | B*15:01 | 66 | 74 | 9 | FVNRYGVAY | 1.49 | 1.25 | -1.66 | 2.74 | 1.09 | 45.5 | 0.13598 | Non-Toxin | 100.00% (86/86) | Cu-Zn superoxide dismutase-like protein |
| 127 | A*31:01 | 36 | 44 | 9 | GTYSLIIHR | 1.18 | 0.66 | -1.12 | 1.84 | 0.72 | 13.2 | 0.07803 | Non-Toxin | 100.00% (86/86) | Cu-Zn superoxide dismutase-like protein |
| 128 | A*11:01 | 36 | 44 | 9 | GTYSLIIHR | 1.18 | 0.66 | -1.21 | 1.84 | 0.63 | 16.4 | 0.07803 | Non-Toxin | 100.00% (86/86) | Cu-Zn superoxide dismutase-like protein |
| 129 | A*68:01 | 36 | 44 | 9 | GTYSLIIHR | 1.18 | 0.66 | -1.24 | 1.84 | 0.61 | 17.3 | 0.07803 | Non-Toxin | 100.00% (86/86) | Cu-Zn superoxide dismutase-like protein |
| 130 | A*30:02 | 36 | 45 | 10 | GTYSLIIHRY | 1.59 | 1.29 | -1.82 | 2.88 | 1.07 | 65.4 | 0.13395 | Non-Toxin | 100.00% (86/86) | Cu-Zn superoxide dismutase-like protein |
| 131 | A*11:01 | 36 | 45 | 10 | GTYSLIIHRY | 1.59 | 1.29 | -2.02 | 2.88 | 0.86 | 105.1 | 0.13395 | Non-Toxin | 100.00% (86/86) | Cu-Zn superoxide dismutase-like protein |
| 132 | A*30:02 | 65 | 74 | 10 | IFVNRYGVAY | 1.49 | 1.45 | -2.25 | 2.94 | 0.69 | 178.8 | 0.14329 | Non-Toxin | 100.00% (86/86) | Cu-Zn superoxide dismutase-like protein |
| 133 | B*15:01 | 65 | 74 | 10 | IFVNRYGVAY | 1.49 | 1.45 | -2.25 | 2.94 | 0.69 | 178.9 | 0.14329 | Non-Toxin | 100.00% (86/86) | Cu-Zn superoxide dismutase-like protein |
| 134 | A*02:06 | 25 | 33 | 9 | KVLGSVIGL | 1.54 | 0.47 | -1.39 | 2.01 | 0.62 | 24.7 | 0.04038 | Non-Toxin | 100.00% (86/86) | Cu-Zn superoxide dismutase-like protein |
| 135 | A*02:03 | 99 | 107 | 9 | RLACGVIGI | 1.23 | 0.33 | -0.82 | 1.57 | 0.75 | 6.6 | 0.16243 | Non-Toxin | 100.00% (86/86) | Cu-Zn superoxide dismutase-like protein |
| 136 | A*30:02 | 69 | 76 | 8 | RYGVAYVY | 1.58 | 1.48 | -2.48 | 3.06 | 0.58 | 304 | 0.122 | Non-Toxin | 100.00% (86/86) | Cu-Zn superoxide dismutase-like protein |
| 137 | A*23:01 | 69 | 77 | 9 | RYGVAYVYL | 1.51 | 0.63 | -1.56 | 2.14 | 0.58 | 36.2 | 0.11984 | Non-Toxin | 100.00% (86/86) | Cu-Zn superoxide dismutase-like protein |
| 138 | A*24:02 | 69 | 77 | 9 | RYGVAYVYL | 1.51 | 0.63 | -1.56 | 2.14 | 0.58 | 36.3 | 0.11984 | Non-Toxin | 100.00% (86/86) | Cu-Zn superoxide dismutase-like protein |
| 139 | A*68:01 | 35 | 44 | 10 | SGTYSLIIHR | 1.18 | 0.58 | -1.12 | 1.76 | 0.64 | 13.2 | 0.09348 | Non-Toxin | 100.00% (86/86) | Cu-Zn superoxide dismutase-like protein |
| 140 | B*15:01 | 29 | 38 | 10 | SVIGLKSGTY | 1.49 | 1.37 | -1.75 | 2.86 | 1.12 | 55.6 | -0.24795 | Non-Toxin | 100.00% (86/86) | Cu-Zn superoxide dismutase-like protein |
| 141 | A*30:02 | 29 | 38 | 10 | SVIGLKSGTY | 1.49 | 1.37 | -1.84 | 2.86 | 1.02 | 69.1 | -0.24795 | Non-Toxin | 100.00% (86/86) | Cu-Zn superoxide dismutase-like protein |
| 142 | A*26:01 | 29 | 38 | 10 | SVIGLKSGTY | 1.49 | 1.37 | -1.97 | 2.86 | 0.89 | 92.4 | -0.24795 | Non-Toxin | 100.00% (86/86) | Cu-Zn superoxide dismutase-like protein |
| 143 | A*23:01 | 108 | 117 | 10 | SYINEKIIHF | 1.36 | 1.27 | -1.19 | 2.63 | 1.44 | 15.6 | 0.18069 | Non-Toxin | 100.00% (86/86) | Cu-Zn superoxide dismutase-like protein |
| 144 | A*24:02 | 108 | 117 | 10 | SYINEKIIHF | 1.36 | 1.27 | -1.41 | 2.63 | 1.22 | 25.8 | 0.18069 | Non-Toxin | 100.00% (86/86) | Cu-Zn superoxide dismutase-like protein |
| 145 | A*30:02 | 30 | 38 | 9 | VIGLKSGTY | 1.49 | 1.31 | -2.16 | 2.8 | 0.64 | 144.5 | -0.31461 | Non-Toxin | 100.00% (86/86) | Cu-Zn superoxide dismutase-like protein |
| 146 | A*30:02 | 67 | 76 | 10 | VNRYGVAYVY | 1.58 | 1.21 | -2.07 | 2.78 | 0.71 | 118.7 | 0.14411 | Non-Toxin | 100.00% (86/86) | Cu-Zn superoxide dismutase-like protein |
| 147 | B*15:01 | 109 | 117 | 9 | YINEKIIHF | 1.36 | 1.13 | -1.72 | 2.49 | 0.77 | 52.7 | 0.14515 | Non-Toxin | 100.00% (86/86) | Cu-Zn superoxide dismutase-like protein |
| 148 | A*02:03 | 109 | 118 | 10 | YINEKIIHFL | 1.63 | 0.45 | -1.14 | 2.08 | 0.94 | 13.7 | 0.23923 | Non-Toxin | 100.00% (86/86) | Cu-Zn superoxide dismutase-like protein |

| | | | | | | | | | | | | | | | |
|---|---|---|---|---|---|---|---|---|---|---|---|---|---|---|---|
| 149 | A*02:01 | 109 | 118 | 10 | YINEKIIHFL | 1.63 | 0.45 | -1.29 | 2.08 | 0.78 | 19.7 | 0.23923 | Non-Toxin | 100.00% (86/86) | Cu-Zn superoxide dismutase-like protein |
| 150 | A*02:06 | 109 | 118 | 10 | YINEKIIHFL | 1.63 | 0.45 | -1.36 | 2.08 | 0.71 | 23 | 0.23923 | Non-Toxin | 100.00% (86/86) | Cu-Zn superoxide dismutase-like protein |
| **#** | **Allele** | **Start** | **End** | **Peptide Length** | **Peptide** | **Proteasome Score** | **TAP Score** | **MHC Score** | **Processing Score** | **Total Score** | **MHC IC50[nM]** | **Immunogenicity** | **Toxicity** | Conservancy | **Protein** |
| 151 | B*58:01 | 12 | 20 | 9 | ATAAVCLLF | 1.1 | 1.18 | -1.6 | 2.28 | 0.68 | 39.7 | 0.02568 | Non-Toxin | 100.00% (574/574) | Envelope protein A28 |
| 152 | A*11:01 | 38 | 47 | 10 | ATHAAFEYSK | 1.14 | 0.33 | -0.93 | 1.47 | 0.53 | 8.6 | 0.19644 | Non-Toxin | 99.83% (573/574) | Envelope protein A28 |
| 153 | B*35:01 | 36 | 45 | 10 | FNATHAAFEY | 1.25 | 1.1 | -1.42 | 2.35 | 0.93 | 26.5 | 0.31549 | Non-Toxin | 100.00% (574/574) | Envelope protein A28 |
| 154 | A*01:01 | 117 | 126 | 10 | FSDVINIDIY | 1.39 | 1.17 | -0.89 | 2.56 | 1.68 | 7.7 | 0.3938 | Non-Toxin | 99.83% (573/574) | Envelope protein A28 |
| 155 | B*35:01 | 83 | 91 | 9 | FVSASIFGF | 1.21 | 1.1 | -1.61 | 2.31 | 0.7 | 41 | 0.06845 | Non-Toxin | 100.00% (574/574) | Envelope protein A28 |
| 156 | B*15:01 | 83 | 91 | 9 | FVSASIFGF | 1.21 | 1.1 | -1.75 | 2.31 | 0.56 | 56.3 | 0.06845 | Non-Toxin | 100.00% (574/574) | Envelope protein A28 |
| 157 | A*30:02 | 21 | 30 | 10 | IQSYSIYENY | 1.38 | 1.26 | -1.78 | 2.65 | 0.87 | 59.9 | -0.01168 | Non-Toxin | 100.00% (574/574) | Envelope protein A28 |
| 158 | B*15:01 | 21 | 30 | 10 | IQSYSIYENY | 1.38 | 1.26 | -1.79 | 2.65 | 0.86 | 61.8 | -0.01168 | Non-Toxin | 100.00% (574/574) | Envelope protein A28 |
| 159 | A*23:01 | 26 | 36 | 11 | IYENYGNIKEF | 1.52 | 1.19 | -2.05 | 2.71 | 0.66 | 111.3 | 0.05087 | Non-Toxin | 100.00% (574/574) | Envelope protein A28 |
| 160 | B*44:03 | 34 | 43 | 10 | KEFNATHAAF | 1.17 | 1.14 | -1.85 | 2.31 | 0.46 | 71.6 | 0.19372 | Non-Toxin | 100.00% (574/574) | Envelope protein A28 |
| 161 | B*15:01 | 70 | 78 | 9 | KQKWRCVVY | 1.61 | 1.37 | -1.62 | 2.98 | 1.36 | 41.6 | 0.2115 | Non-Toxin | 100.00% (574/574) | Envelope protein A28 |
| 162 | A*30:02 | 70 | 78 | 9 | KQKWRCVVY | 1.61 | 1.37 | -1.71 | 2.98 | 1.26 | 51.5 | 0.2115 | Non-Toxin | 100.00% (574/574) | Envelope protein A28 |
| 163 | A*30:01 | 70 | 78 | 9 | KQKWRCVVY | 1.61 | 1.37 | -2.47 | 2.98 | 0.51 | 292.7 | 0.2115 | Non-Toxin | 100.00% (574/574) | Envelope protein A28 |
| 164 | B*15:01 | 18 | 27 | 10 | LLFIQSYSIY | 1.24 | 1.33 | -1.22 | 2.57 | 1.35 | 16.7 | -0.16732 | Non-Toxin | 100.00% (574/574) | Envelope protein A28 |
| 165 | A*30:02 | 18 | 27 | 10 | LLFIQSYSIY | 1.24 | 1.33 | -2.08 | 2.57 | 0.49 | 121.5 | -0.16732 | Non-Toxin | 100.00% (574/574) | Envelope protein A28 |
| 166 | B*35:01 | 37 | 45 | 9 | NATHAAFEY | 1.25 | 1.25 | -0.91 | 2.5 | 1.59 | 8.2 | 0.27738 | Non-Toxin | 100.00% (574/574) | Envelope protein A28 |
| 167 | A*30:02 | 22 | 30 | 9 | QSYSIYENY | 1.38 | 1.38 | -1.84 | 2.76 | 0.92 | 69.5 | 0.03917 | Non-Toxin | 100.00% (574/574) | Envelope protein A28 |
| 168 | B*58:01 | 22 | 30 | 9 | QSYSIYENY | 1.38 | 1.38 | -2.28 | 2.76 | 0.48 | 191.2 | 0.03917 | Non-Toxin | 100.00% (574/574) | Envelope protein A28 |
| 169 | A*31:01 | 101 | 110 | 10 | RSIRKFNTMR | 0.93 | 0.8 | -0.96 | 1.73 | 0.77 | 9.2 | -0.07665 | Non-Toxin | 100.00% (574/574) | Envelope protein A28 |
| 170 | A*02:03 | 87 | 95 | 9 | SIFGFQAEV | 1.06 | 0.33 | -0.82 | 1.39 | 0.57 | 6.6 | 0.16858 | Non-Toxin | 100.00% (574/574) | Envelope protein A28 |
| 171 | A*31:01 | 102 | 110 | 9 | SIRKFNTMR | 0.93 | 0.8 | -0.96 | 1.73 | 0.77 | 9.1 | -0.16213 | Non-Toxin | 100.00% (574/574) | Envelope protein A28 |
| 172 | A*01:01 | 116 | 126 | 11 | TFSDVINIDIY | 1.39 | 1.29 | -1.86 | 2.67 | 0.82 | 71.8 | 0.35388 | Non-Toxin | 99.83% (573/574) | Envelope protein A28 |
| 173 | B*15:01 | 108 | 117 | 10 | TMRQCIDFTF | 1.59 | 1.23 | -1.46 | 2.81 | 1.35 | 29 | 0.1197 | Non-Toxin | 100.00% (574/574) | Envelope protein A28 |
| 174 | A*23:01 | 108 | 117 | 10 | TMRQCIDFTF | 1.59 | 1.23 | -1.97 | 2.81 | 0.85 | 92.4 | 0.1197 | Non-Toxin | 100.00% (574/574) | Envelope protein A28 |
| 175 | A*32:01 | 108 | 117 | 10 | TMRQCIDFTF | 1.59 | 1.23 | -2.13 | 2.81 | 0.69 | 133.6 | 0.1197 | Non-Toxin | 100.00% (574/574) | Envelope protein A28 |
| 176 | B*58:01 | 11 | 20 | 10 | VATAAVCLLF | 1.1 | 1.15 | -1.56 | 2.26 | 0.7 | 36.3 | 0.06368 | Non-Toxin | 100.00% (574/574) | Envelope protein A28 |
| 177 | B*15:01 | 69 | 78 | 10 | VKQKWRCVVY | 1.61 | 1.24 | -2 | 2.85 | 0.85 | 100 | 0.01971 | Non-Toxin | 100.00% (574/574) | Envelope protein A28 |
| **#** | **Allele** | **Start** | **End** | **Peptide Length** | **Peptide** | **Proteasome Score** | **TAP Score** | **MHC Score** | **Processing Score** | **Total Score** | **MHC IC50[nM]** | **Immunogenicity** | **Toxicity** | Conservancy | **Protein** |
| 178 | B*35:01 | 832 | 840 | 9 | FARDNQISF | 1.35 | 1.04 | -0.83 | 2.39 | 1.56 | 6.8 | -0.06056 | Non-Toxin | 100.00% (10/10) | DNA-directed RNA polymerase |
| 179 | B*35:01 | 663 | 671 | 9 | FPAEFRDGY | 1.16 | 1.04 | -0.54 | 2.21 | 1.66 | 3.5 | 0.31469 | Non-Toxin | 100.00% (10/10) | DNA-directed RNA polymerase |
| 180 | B*35:01 | 528 | 536 | 9 | FPITIENAL | 1.46 | 0.26 | -0.7 | 1.72 | 1.02 | 5 | 0.32351 | Non-Toxin | 100.00% (10/10) | DNA-directed RNA polymerase |
| 181 | B*15:01 | 33 | 41 | 9 | FQYVSYSNF | 1.15 | 1.16 | -0.97 | 2.31 | 1.34 | 9.3 | -0.26764 | Non-Toxin | 100.00% (10/10) | DNA-directed RNA polymerase |
| 182 | A*23:01 | 329 | 338 | 10 | FYMISLLRKF | 1.17 | 1.19 | -1.07 | 2.36 | 1.29 | 11.8 | -0.18774 | Non-Toxin | 100.00% (10/10) | DNA-directed RNA polymerase |
| 183 | A*24:02 | 329 | 338 | 10 | FYMISLLRKF | 1.17 | 1.19 | -1.35 | 2.36 | 1.01 | 22.3 | -0.18774 | Non-Toxin | 100.00% (10/10) | DNA-directed RNA polymerase |
| 184 | A*30:02 | 424 | 432 | 9 | GSYRTHPHY | 1.4 | 1.23 | -1.56 | 2.63 | 1.07 | 36.7 | 0.12867 | Non-Toxin | 100.00% (10/10) | DNA-directed RNA polymerase |
| 185 | B*35:01 | 896 | 904 | 9 | IADETELPY | 0.93 | 1.2 | -0.9 | 2.13 | 1.23 | 8 | 0.22416 | Non-Toxin | 100.00% (10/10) | DNA-directed RNA polymerase |
| 186 | B*58:01 | 838 | 846 | 9 | ISFDVSEKY | 1.57 | 1.32 | -1.8 | 2.89 | 1.09 | 63.1 | -0.09671 | Non-Toxin | 100.00% (10/10) | DNA-directed RNA polymerase |
| 187 | A*30:02 | 438 | 447 | 10 | ISIPRSVGFY | 1.36 | 1.39 | -1.32 | 2.75 | 1.43 | 20.9 | 0.0572 | Non-Toxin | 100.00% (10/10) | DNA-directed RNA polymerase |
| 188 | A*01:01 | 278 | 286 | 9 | ITDFDIDTY | 1.26 | 1.22 | -1.21 | 2.48 | 1.27 | 16.1 | 0.31328 | Non-Toxin | 60.00% (6/10) | DNA-directed RNA polymerase |
| 189 | A*01:01 | 494 | 502 | 9 | ITNILTSEY | 1.33 | 1.25 | -1.31 | 2.59 | 1.27 | 20.6 | 0.07644 | Non-Toxin | 100.00% (10/10) | DNA-directed RNA polymerase |
| 190 | A*30:02 | 494 | 502 | 9 | ITNILTSEY | 1.33 | 1.25 | -1.33 | 2.59 | 1.25 | 21.5 | 0.07644 | Non-Toxin | 100.00% (10/10) | DNA-directed RNA polymerase |
| 191 | A*30:02 | 508 | 517 | 10 | KICEYIRSYY | 1.32 | 1.42 | -1.71 | 2.74 | 1.02 | 51.6 | 0.11619 | Non-Toxin | 100.00% (10/10) | DNA-directed RNA polymerase |
| 192 | A*30:02 | 708 | 717 | 10 | KIDNGIHLMY | 1.28 | 1.36 | -1.65 | 2.64 | 0.99 | 44.7 | 0.08178 | Non-Toxin | 100.00% (10/10) | DNA-directed RNA polymerase |
| 193 | B*35:01 | 184 | 192 | 9 | KPNSFTFSF | 1.26 | 0.99 | -1.18 | 2.25 | 1.07 | 15.1 | -0.01589 | Non-Toxin | 100.00% (10/10) | DNA-directed RNA polymerase |
| 194 | A*32:01 | 90 | 98 | 9 | KSYDALATF | 1.5 | 1.23 | -0.9 | 2.73 | 1.83 | 8 | 0.10448 | Non-Toxin | 100.00% (10/10) | DNA-directed RNA polymerase |
| 195 | B*58:01 | 90 | 98 | 9 | KSYDALATF | 1.5 | 1.23 | -1.1 | 2.73 | 1.63 | 12.5 | 0.10448 | Non-Toxin | 100.00% (10/10) | DNA-directed RNA polymerase |
| 196 | B*57:01 | 90 | 98 | 9 | KSYDALATF | 1.5 | 1.23 | -1.38 | 2.73 | 1.35 | 24.1 | 0.10448 | Non-Toxin | 100.00% (10/10) | DNA-directed RNA polymerase |
| 197 | B*15:01 | 90 | 98 | 9 | KSYDALATF | 1.5 | 1.23 | -1.4 | 2.73 | 1.33 | 25.1 | 0.10448 | Non-Toxin | 100.00% (10/10) | DNA-directed RNA polymerase |
| 198 | A*30:02 | 1007 | 1015 | 9 | KVFFGPIYY | 1.13 | 1.45 | -1.12 | 2.57 | 1.45 | 13.2 | 0.28852 | Non-Toxin | 100.00% (10/10) | DNA-directed RNA polymerase |

| | | | | | | | | | | | | | | | |
|---|---|---|---|---|---|---|---|---|---|---|---|---|---|---|---|
| 199 | A*11:01 | 1007 | 1015 | 9 | KVFFGPIYY | 1.13 | 1.45 | -1.2 | 2.57 | 1.38 | 15.8 | 0.28852 | Non-Toxin | 100.00% (10/10) | DNA-directed RNA polymerase |
| 200 | A*03:01 | 1007 | 1015 | 9 | KVFFGPIYY | 1.13 | 1.45 | -1.23 | 2.57 | 1.35 | 16.8 | 0.28852 | Non-Toxin | 100.00% (10/10) | DNA-directed RNA polymerase |
| 201 | A*32:01 | 1007 | 1015 | 9 | KVFFGPIYY | 1.13 | 1.45 | -1.59 | 2.57 | 0.98 | 39 | 0.28852 | Non-Toxin | 100.00% (10/10) | DNA-directed RNA polymerase |
| 202 | A*30:02 | 461 | 469 | 9 | KYHPSQYLY | 1.26 | 1.5 | -1.37 | 2.76 | 1.39 | 23.4 | -0.2804 | Non-Toxin | 100.00% (10/10) | DNA-directed RNA polymerase |
| 203 | A*23:01 | 461 | 470 | 10 | KYHPSQYLYF | 1.15 | 1.33 | -0.96 | 2.49 | 1.53 | 9.1 | -0.28956 | Non-Toxin | 100.00% (10/10) | DNA-directed RNA polymerase |
| 204 | A*24:02 | 461 | 470 | 10 | KYHPSQYLYF | 1.15 | 1.33 | -0.96 | 2.49 | 1.52 | 9.2 | -0.28956 | Non-Toxin | 100.00% (10/10) | DNA-directed RNA polymerase |
| 205 | B*15:01 | 934 | 943 | 10 | LTAAYSTKPY | 1.13 | 1.27 | -1.4 | 2.4 | 1 | 24.9 | -0.26457 | Non-Toxin | 90.00% (9/10) | DNA-directed RNA polymerase |
| 206 | A*68:01 | 544 | 553 | 10 | MICDFVTDFR | 0.92 | 0.79 | -0.7 | 1.7 | 1.01 | 5 | 0.28268 | Non-Toxin | 100.00% (10/10) | DNA-directed RNA polymerase |
| 207 | B*15:01 | 928 | 938 | 11 | MLIEVILTAAY | 1.38 | 1.35 | -1.5 | 2.73 | 1.23 | 31.7 | 0.39537 | Non-Toxin | 90.00% (9/10) | DNA-directed RNA polymerase |
| 208 | B*15:01 | 966 | 975 | 10 | MQFAKQCYEY | 1.36 | 1.35 | -1.34 | 2.71 | 1.36 | 22.1 | -0.2408 | Non-Toxin | 100.00% (10/10) | DNA-directed RNA polymerase |
| 209 | A*30:02 | 966 | 975 | 10 | MQFAKQCYEY | 1.36 | 1.35 | -1.44 | 2.71 | 1.27 | 27.4 | -0.2408 | Non-Toxin | 100.00% (10/10) | DNA-directed RNA polymerase |
| 210 | B*35:01 | 629 | 637 | 9 | MVDIDQFTF | 1.48 | 1.03 | -1.41 | 2.51 | 1.09 | 26 | 0.17516 | Non-Toxin | 0.00% (0/10) | DNA-directed RNA polymerase |
| 211 | A*23:01 | 238 | 247 | 10 | QYLSRVSLEF | 1.08 | 1.25 | -1.08 | 2.33 | 1.26 | 11.9 | -0.18606 | Non-Toxin | 100.00% (10/10) | DNA-directed RNA polymerase |
| 212 | A*24:02 | 238 | 247 | 10 | QYLSRVSLEF | 1.08 | 1.25 | -1.28 | 2.33 | 1.06 | 18.9 | -0.18606 | Non-Toxin | 100.00% (10/10) | DNA-directed RNA polymerase |
| 213 | A*30:02 | 460 | 469 | 10 | RKYHPSQYLY | 1.26 | 1.5 | -1.44 | 2.76 | 1.32 | 27.6 | -0.25919 | Non-Toxin | 100.00% (10/10) | DNA-directed RNA polymerase |
| 214 | B*07:02 | 720 | 728 | 9 | RPIVISKAL | 1.57 | 0.35 | -0.72 | 1.91 | 1.2 | 5.2 | -0.10053 | Non-Toxin | 100.00% (10/10) | DNA-directed RNA polymerase |
| 215 | A*30:02 | 514 | 523 | 10 | RSYYKDDISY | 1.3 | 1.48 | -1.62 | 2.77 | 1.16 | 41.4 | -0.15678 | Non-Toxin | 100.00% (10/10) | DNA-directed RNA polymerase |
| 216 | A*32:01 | 181 | 190 | 10 | RVVKPNSFTF | 1.44 | 1.25 | -1.47 | 2.7 | 1.23 | 29.6 | -0.25495 | Non-Toxin | 100.00% (10/10) | DNA-directed RNA polymerase |
| 217 | B*15:01 | 181 | 190 | 10 | RVVKPNSFTF | 1.44 | 1.25 | -1.71 | 2.7 | 0.98 | 51.7 | -0.25495 | Non-Toxin | 100.00% (10/10) | DNA-directed RNA polymerase |
| 218 | B*44:03 | 7 | 15 | 9 | SEMDQRLGY | 1.48 | 1.28 | -1.23 | 2.76 | 1.53 | 17 | -0.08832 | Non-Toxin | 100.00% (10/10) | DNA-directed RNA polymerase |
| 219 | B*44:02 | 7 | 15 | 9 | SEMDQRLGY | 1.48 | 1.28 | -1.31 | 2.76 | 1.45 | 20.5 | -0.08832 | Non-Toxin | 100.00% (10/10) | DNA-directed RNA polymerase |
| 220 | B*15:01 | 353 | 362 | 10 | SMVCHRILTY | 1.46 | 1.32 | -1.27 | 2.78 | 1.51 | 18.7 | 0.17965 | Non-Toxin | 100.00% (10/10) | DNA-directed RNA polymerase |
| 221 | A*23:01 | 515 | 524 | 10 | SYYKDDISYF | 1.34 | 1.38 | -1.27 | 2.72 | 1.45 | 18.7 | -0.1915 | Non-Toxin | 100.00% (10/10) | DNA-directed RNA polymerase |
| 222 | A*24:02 | 515 | 524 | 10 | SYYKDDISYF | 1.34 | 1.38 | -1.58 | 2.72 | 1.14 | 38.1 | -0.1915 | Non-Toxin | 100.00% (10/10) | DNA-directed RNA polymerase |
| 223 | B*35:01 | 935 | 943 | 9 | TAAYSTKPY | 1.13 | 1.31 | -1.03 | 2.44 | 1.41 | 10.8 | -0.30406 | Non-Toxin | 100.00% (10/10) | DNA-directed RNA polymerase |
| 224 | A*24:02 | 361 | 369 | 9 | TYGKYFETL | 1.77 | 0.47 | -1.23 | 2.24 | 1.01 | 17.1 | 0.00778 | Non-Toxin | 100.00% (10/10) | DNA-directed RNA polymerase |
| 225 | A*23:01 | 361 | 369 | 9 | TYGKYFETL | 1.77 | 0.47 | -1.26 | 2.24 | 0.98 | 18.4 | 0.00778 | Non-Toxin | 100.00% (10/10) | DNA-directed RNA polymerase |
| 226 | B*15:01 | 913 | 921 | 9 | VIINSTSIF | 1.16 | 1.15 | -1.12 | 2.31 | 1.19 | 13.2 | -0.14973 | Non-Toxin | 100.00% (10/10) | DNA-directed RNA polymerase |
| 227 | A*01:01 | 842 | 850 | 9 | VSEKYTDMY | 1.21 | 1.19 | -1.4 | 2.4 | 1 | 25 | -0.23544 | Non-Toxin | 100.00% (10/10) | DNA-directed RNA polymerase |
| 228 | B*15:01 | 182 | 190 | 9 | VVKPNSFTF | 1.44 | 1.24 | -1.73 | 2.69 | 0.95 | 53.9 | -0.12171 | Non-Toxin | 100.00% (10/10) | DNA-directed RNA polymerase |
| 229 | B*35:01 | 895 | 904 | 10 | YIADETELPY | 0.93 | 1.39 | -1.18 | 2.32 | 1.13 | 15.3 | 0.24873 | Non-Toxin | 100.00% (10/10) | DNA-directed RNA polymerase |
| 230 | A*01:01 | 895 | 904 | 10 | YIADETELPY | 0.93 | 1.39 | -1.26 | 2.32 | 1.06 | 18.2 | 0.24873 | Non-Toxin | 100.00% (10/10) | DNA-directed RNA polymerase |
| 231 | B*15:01 | 330 | 338 | 9 | YMISLLRKF | 1.17 | 1.13 | -1.12 | 2.3 | 1.18 | 13.1 | -0.22683 | Non-Toxin | 100.00% (10/10) | DNA-directed RNA polymerase |
| 232 | B*15:01 | 965 | 973 | 9 | YMQFAKQCY | 1.34 | 1.37 | -1.61 | 2.71 | 1.09 | 41.1 | -0.21396 | Non-Toxin | 100.00% (10/10) | DNA-directed RNA polymerase |
| 233 | A*01:01 | 321 | 330 | 10 | YTPDQLKGFY | 1.4 | 1.31 | -0.9 | 2.71 | 1.8 | 8 | -0.21088 | Non-Toxin | 100.00% (10/10) | DNA-directed RNA polymerase |
| 234 | A*26:01 | 321 | 330 | 10 | YTPDQLKGFY | 1.4 | 1.31 | -1.58 | 2.71 | 1.13 | 37.9 | -0.21088 | Non-Toxin | 100.00% (10/10) | DNA-directed RNA polymerase |
| 235 | A*02:03 | 671 | 679 | 9 | YVASSLVGI | 1.42 | 0.28 | -0.66 | 1.7 | 1.03 | 4.6 | -0.27067 | Non-Toxin | 100.00% (10/10) | DNA-directed RNA polymerase |
| 236 | A*02:06 | 671 | 679 | 9 | YVASSLVGI | 1.42 | 0.28 | -0.67 | 1.7 | 1.02 | 4.7 | -0.27067 | Non-Toxin | 100.00% (10/10) | DNA-directed RNA polymerase |
| 237 | A*23:01 | 516 | 524 | 9 | YYKDDISYF | 1.34 | 1.23 | -1.56 | 2.57 | 1.01 | 36 | -0.04258 | Non-Toxin | 100.00% (10/10) | DNA-directed RNA polymerase |

| # | **Allele** | **Start** | **End** | **Peptide Length** | **Peptide** | **Proteasome Score** | **TAP Score** | **MHC Score** | **Processing Score** | **Total Score** | **MHC IC50[nM]** | **Immunogenicity** | **Toxicity** | Conservancy | **Protein** |
|---|---|---|---|---|---|---|---|---|---|---|---|---|---|---|---|
| 238 | B*44:03 | 2 | 10 | 9 | AEFEDQLVF | 1.48 | 1.17 | -1.22 | 2.65 | 1.42 | 16.7 | 0.06607 | Non-Toxin | 99.66% (580/582) | Telomere-binding protein |
| 239 | B*40:01 | 2 | 10 | 9 | AEFEDQLVF | 1.48 | 1.17 | -1.25 | 2.65 | 1.39 | 17.9 | 0.06607 | Non-Toxin | 99.66% (580/582) | Telomere-binding protein |
| 240 | B*44:02 | 2 | 10 | 9 | AEFEDQLVF | 1.48 | 1.17 | -1.61 | 2.65 | 1.04 | 40.3 | 0.06607 | Non-Toxin | 99.66% (580/582) | Telomere-binding protein |
| 241 | A*33:01 | 190 | 198 | 9 | DVLKTRLFR | 1.07 | 0.58 | -1.03 | 1.65 | 0.63 | 10.6 | -0.07504 | Non-Toxin | 99.83% (581/582) | Telomere-binding protein |
| 242 | A*33:01 | 214 | 222 | 9 | DYFTRVTKR | 0.92 | 0.65 | -0.85 | 1.56 | 0.71 | 7.1 | 0.07308 | Non-Toxin | 99.66% (580/582) | Telomere-binding protein |
| 243 | A*26:01 | 243 | 252 | 10 | ETVKMGAFMY | 1.22 | 1.23 | -1.4 | 2.45 | 1.05 | 25 | -0.30857 | Non-Toxin | 99.66% (580/582) | Telomere-binding protein |
| 244 | A*02:03 | 250 | 258 | 9 | FMYTKHSML | 1.47 | 0.58 | -0.69 | 2.04 | 1.35 | 4.9 | -0.38391 | Non-Toxin | 99.83% (581/582) | Telomere-binding protein |
| 245 | B*08:01 | 250 | 258 | 9 | FMYTKHSML | 1.47 | 0.58 | -1.12 | 2.04 | 0.92 | 13.2 | -0.38391 | Non-Toxin | 99.83% (581/582) | Telomere-binding protein |
| 246 | A*02:01 | 250 | 258 | 9 | FMYTKHSML | 1.47 | 0.58 | -1.36 | 2.04 | 0.69 | 22.7 | -0.38391 | Non-Toxin | 99.83% (581/582) | Telomere-binding protein |
| 247 | B*15:01 | 62 | 71 | 10 | FVKKFGLCNY | 1.37 | 1.31 | -1.99 | 2.68 | 0.69 | 96.7 | -0.19972 | Non-Toxin | 100.00% (582/582) | Telomere-binding protein |
| 248 | A*02:03 | 74 | 82 | 9 | ILISLINSL | 1.5 | 0.45 | -0.92 | 1.95 | 1.03 | 8.3 | -0.11091 | Non-Toxin | 99.83% (581/582) | Telomere-binding protein |
| 249 | A*02:01 | 74 | 82 | 9 | ILISLINSL | 1.5 | 0.45 | -1.22 | 1.95 | 0.73 | 16.5 | -0.11091 | Non-Toxin | 99.83% (581/582) | Telomere-binding protein |

| | | | | | | | | | | | | | | | |
|---|---|---|---|---|---|---|---|---|---|---|---|---|---|---|---|
| 250 | B*15:01 | 13 | 21 | 9 | ISARALKAY | 1.54 | 1.33 | -1.94 | 2.87 | 0.92 | 87.9 | -0.0667 | Non-Toxin | 99.48% (579/582) | Telomere-binding protein |
| 251 | A*30:02 | 13 | 21 | 9 | ISARALKAY | 1.54 | 1.33 | -1.97 | 2.87 | 0.9 | 92.6 | -0.0667 | Non-Toxin | 99.48% (579/582) | Telomere-binding protein |
| 252 | B*58:01 | 13 | 22 | 10 | ISARALKAYF | 1.12 | 1.16 | -1.57 | 2.27 | 0.7 | 37.1 | -0.08006 | Non-Toxin | 99.48% (579/582) | Telomere-binding protein |
| 253 | A*30:02 | 207 | 215 | 9 | ISRTRLYDY | 1.21 | 1.32 | -1.93 | 2.54 | 0.61 | 85.2 | 0.10566 | Non-Toxin | 99.83% (581/582) | Telomere-binding protein |
| 254 | A*30:02 | 174 | 182 | 9 | KSSIKDSMY | 1.35 | 1.26 | -1.67 | 2.61 | 0.93 | 47.2 | -0.35112 | Non-Toxin | 100.00% (582/582) | Telomere-binding protein |
| 255 | A*30:01 | 193 | 202 | 10 | KTRLFRSPQV | 1.28 | 0.21 | -0.71 | 1.5 | 0.79 | 5.1 | -0.06273 | Non-Toxin | 99.83% (581/582) | Telomere-binding protein |
| 256 | A*31:01 | 129 | 137 | 9 | KVLAARLKR | 1.12 | 0.78 | -1.12 | 1.9 | 0.78 | 13.3 | -0.01277 | Non-Toxin | 100.00% (582/582) | Telomere-binding protein |
| 257 | A*68:01 | 8 | 16 | 9 | LVFNSISAR | 1.06 | 0.73 | -0.89 | 1.79 | 0.9 | 7.7 | -0.12109 | Non-Toxin | 99.48% (579/582) | Telomere-binding protein |
| 258 | B*35:01 | 1 | 10 | 10 | MAEFEDQLVF | 1.48 | 1.06 | -1.2 | 2.53 | 1.33 | 15.9 | 0.17512 | Non-Toxin | 99.66% (580/582) | Telomere-binding protein |
| 259 | B*58:01 | 1 | 10 | 10 | MAEFEDQLVF | 1.48 | 1.06 | -1.38 | 2.53 | 1.15 | 24.1 | 0.17512 | Non-Toxin | 99.66% (580/582) | Telomere-binding protein |
| 260 | B*53:01 | 1 | 10 | 10 | MAEFEDQLVF | 1.48 | 1.06 | -1.71 | 2.53 | 0.82 | 51.6 | 0.17512 | Non-Toxin | 99.66% (580/582) | Telomere-binding protein |
| 261 | B*44:03 | 1 | 10 | 10 | MAEFEDQLVF | 1.48 | 1.06 | -1.76 | 2.53 | 0.77 | 58.2 | 0.17512 | Non-Toxin | 99.66% (580/582) | Telomere-binding protein |
| 262 | B*15:01 | 1 | 10 | 10 | MAEFEDQLVF | 1.48 | 1.06 | -1.94 | 2.53 | 0.59 | 87.2 | 0.17512 | Non-Toxin | 99.66% (580/582) | Telomere-binding protein |
| 263 | A*26:01 | 205 | 213 | 9 | NIISRTRLY | 1.15 | 1.31 | -1.86 | 2.46 | 0.6 | 71.7 | 0.00087 | Non-Toxin | 99.83% (581/582) | Telomere-binding protein |
| 264 | A*68:02 | 113 | 121 | 9 | NTIDKSSPL | 1.26 | 0.47 | -1.15 | 1.73 | 0.58 | 1 | -0.44631 | Non-Toxin | 99.48% (579/582) | Telomere-binding protein |
| 265 | A*01:01 | 113 | 122 | 10 | NTIDKSSPLY | 1.29 | 1.33 | -1.31 | 2.62 | 1.31 | 20.6 | -0.47715 | Non-Toxin | 99.48% (579/582) | Telomere-binding protein |
| 266 | A*26:01 | 113 | 122 | 10 | NTIDKSSPLY | 1.29 | 1.33 | -1.68 | 2.62 | 0.95 | 47.6 | -0.47715 | Non-Toxin | 99.48% (579/582) | Telomere-binding protein |
| 267 | B*40:01 | 222 | 231 | 10 | RDESSIYVIL | 1.76 | 0.37 | -1.36 | 2.13 | 0.77 | 23.1 | -0.05635 | Non-Toxin | 99.48% (579/582) | Telomere-binding protein |
| 268 | B*15:01 | 134 | 142 | 9 | RLKRSATQF | 1.48 | 1.28 | -1.58 | 2.76 | 1.18 | 38.3 | -0.17711 | Non-Toxin | 100.00% (582/582) | Telomere-binding protein |
| 269 | A*02:03 | 211 | 219 | 9 | RLYDYFTRV | 0.97 | 0.34 | -0.36 | 1.31 | 0.95 | 2.3 | 0.19072 | Non-Toxin | 99.83% (581/582) | Telomere-binding protein |
| 270 | A*02:01 | 211 | 219 | 9 | RLYDYFTRV | 0.97 | 0.34 | -0.46 | 1.31 | 0.85 | 2.9 | 0.19072 | Non-Toxin | 99.83% (581/582) | Telomere-binding protein |
| 271 | A*02:06 | 211 | 219 | 9 | RLYDYFTRV | 0.97 | 0.34 | -0.57 | 1.31 | 0.74 | 3.7 | 0.19072 | Non-Toxin | 99.83% (581/582) | Telomere-binding protein |
| 272 | A*31:01 | 209 | 218 | 10 | RTRLYDYFTR | 1.12 | 0.79 | -0.78 | 1.91 | 1.13 | 6 | 0.14164 | Non-Toxin | 99.83% (581/582) | Telomere-binding protein |
| 273 | A*24:02 | 268 | 277 | 10 | RYSKKFQESF | 1.31 | 1.26 | -1.44 | 2.57 | 1.12 | 27.7 | -0.4879 | Non-Toxin | 99.83% (581/582) | Telomere-binding protein |
| 274 | A*23:01 | 268 | 277 | 10 | RYSKKFQESF | 1.31 | 1.26 | -1.57 | 2.57 | 1 | 36.8 | -0.4879 | Non-Toxin | 99.83% (581/582) | Telomere-binding protein |
| 275 | A*23:01 | 276 | 284 | 9 | SFYEDIAEF | 1.09 | 1.2 | -1.63 | 2.3 | 0.67 | 42.4 | 0.33795 | Non-Toxin | 99.83% (581/582) | Telomere-binding protein |
| 276 | B*15:01 | 12 | 21 | 10 | SISARALKAY | 1.54 | 1.41 | -2.08 | 2.95 | 0.87 | 119.9 | -0.09541 | Non-Toxin | 99.48% (579/582) | Telomere-binding protein |
| 277 | A*30:02 | 12 | 21 | 10 | SISARALKAY | 1.54 | 1.41 | -2.33 | 2.95 | 0.61 | 215.4 | -0.09541 | Non-Toxin | 99.48% (579/582) | Telomere-binding protein |
| 278 | A*03:01 | 12 | 21 | 10 | SISARALKAY | 1.54 | 1.41 | -2.33 | 2.95 | 0.61 | 216.1 | -0.09541 | Non-Toxin | 99.48% (579/582) | Telomere-binding protein |
| 279 | A*01:01 | 114 | 122 | 9 | TIDKSSPLY | 1.29 | 1.3 | -1.49 | 2.6 | 1.1 | 31 | -0.54247 | Non-Toxin | 99.48% (579/582) | Telomere-binding protein |
| 280 | B*15:01 | 140 | 149 | 10 | TQFNFNGHTY | 1.45 | 1.41 | -1.35 | 2.86 | 1.52 | 22.3 | 0.22107 | Non-Toxin | 100.00% (582/582) | Telomere-binding protein |
| 281 | A*30:02 | 140 | 149 | 10 | TQFNFNGHTY | 1.45 | 1.41 | -1.72 | 2.86 | 1.14 | 52.5 | 0.22107 | Non-Toxin | 100.00% (582/582) | Telomere-binding protein |
| 282 | A*30:02 | 269 | 278 | 10 | YSKKFQESFY | 1.41 | 1.34 | -1.71 | 2.74 | 1.03 | 51.4 | -0.26277 | Non-Toxin | 99.83% (581/582) | Telomere-binding protein |
| **#** | **Allele** | **Start** | **End** | **Peptide Length** | **Peptide** | **Proteasome Score** | **TAP Score** | **MHC Score** | **Processing Score** | **Total Score** | **MHC IC50[nM]** | **Immunogenicity** | **Toxicity** | Conservancy | **Protein** |
| 283 | B*15:01 | 96 | 105 | 10 | IARTHTALIF | 1.39 | 1.17 | -1.89 | 2.56 | 0.67 | 78.3 | 0.23039 | Non-Toxin | 99.48% (578/581) | Profilin |
| 284 | B*58:01 | 96 | 105 | 10 | IARTHTALIF | 1.39 | 1.17 | -2.24 | 2.56 | 0.32 | 175.1 | 0.23039 | Non-Toxin | 99.48% (578/581) | Profilin |
| 285 | B*15:01 | 57 | 66 | 10 | ILKPLIGQKF | 1.51 | 1.18 | -2.06 | 2.69 | 0.63 | 115.2 | -0.15422 | Non-Toxin | 96.90% (563/581) | Profilin |
| 286 | B*15:01 | 32 | 40 | 9 | LAAIPNRTF | 1.49 | 1.15 | -2.32 | 2.64 | 0.32 | 208.7 | 0.19609 | Non-Toxin | 99.83% (580/581) | Profilin |
| 287 | A*02:01 | 74 | 82 | 9 | LMDENTYAM | 1.05 | 0.13 | -0.72 | 1.19 | 0.47 | 5.2 | 0.15793 | Non-Toxin | 100.00% (581/581) | Profilin |
| 288 | A*02:06 | 74 | 82 | 9 | LMDENTYAM | 1.05 | 0.13 | -0.87 | 1.19 | 0.32 | 7.4 | 0.15793 | Non-Toxin | 100.00% (581/581) | Profilin |
| 289 | B*35:01 | 72 | 80 | 9 | NSLMDENTY | 1.61 | 1.32 | -2.01 | 2.93 | 0.92 | 102.9 | -0.04723 | Non-Toxin | 99.83% (580/581) | Profilin |
| 290 | A*68:01 | 30 | 38 | 9 | NVLAAIPNR | 1.06 | 0.77 | -1.47 | 1.84 | 0.37 | 29.5 | 0.18601 | Non-Toxin | 99.83% (580/581) | Profilin |
| 291 | A*31:01 | 119 | 127 | 9 | RTCRDHATR | 0.99 | 0.72 | -1.16 | 1.71 | 0.55 | 14.5 | 0.14233 | Non-Toxin | 99.66% (579/581) | Profilin |
| 292 | A*30:01 | 98 | 106 | 9 | RTHTALIFL | 1.68 | 0.59 | -1.08 | 2.28 | 1.2 | 11.9 | 0.25794 | Non-Toxin | 99.48% (578/581) | Profilin |
| 293 | A*02:03 | 73 | 81 | 9 | SLMDENTYA | 1.18 | -0.11 | -0.73 | 1.07 | 0.34 | 5.4 | 0.08733 | Non-Toxin | 100.00% (581/581) | Profilin |
| 294 | A*68:01 | 111 | 119 | 9 | TTSRRDVYR | 0.98 | 0.56 | -0.86 | 1.54 | 0.68 | 7.2 | 0.10234 | Non-Toxin | 100.00% (581/581) | Profilin |
| 295 | B*15:01 | 79 | 88 | 10 | TYAMELLTGY | 1.34 | 1.44 | -2.15 | 2.78 | 0.64 | 140.2 | -0.03518 | Non-Toxin | 99.83% (580/581) | Profilin |
| 296 | B*35:01 | 79 | 88 | 10 | TYAMELLTGY | 1.34 | 1.44 | -2.18 | 2.78 | 0.6 | 151.2 | -0.03518 | Non-Toxin | 99.83% (580/581) | Profilin |
| 297 | B*15:01 | 31 | 40 | 10 | VLAAIPNRTF | 1.49 | 1.2 | -1.69 | 2.7 | 1 | 49.5 | 0.23114 | Non-Toxin | 99.83% (580/581) | Profilin |
| 298 | B*35:01 | 80 | 88 | 9 | YAMELLTGY | 1.34 | 1.32 | -0.46 | 2.66 | 2.19 | 2.9 | 0.07507 | Non-Toxin | 99.83% (580/581) | Profilin |
| 299 | B*15:01 | 80 | 88 | 9 | YAMELLTGY | 1.34 | 1.32 | -1.24 | 2.66 | 1.42 | 17.4 | 0.07507 | Non-Toxin | 99.83% (580/581) | Profilin |
| 300 | A*26:01 | 80 | 88 | 9 | YAMELLTGY | 1.34 | 1.32 | -1.4 | 2.66 | 1.26 | 24.9 | 0.07507 | Non-Toxin | 99.83% (580/581) | Profilin |

| | | | | | | | | | | | | | | | |
|---|---|---|---|---|---|---|---|---|---|---|---|---|---|---|---|
| 301 | B*53:01 | 80 | 88 | 9 | YAMELLTGY | 1.34 | 1.32 | -1.81 | 2.66 | 0.84 | 65.2 | 0.07507 | Non-Toxin | 99.83% (580/581) | Profilin |
| 302 | A*68:01 | 80 | 88 | 9 | YAMELLTGY | 1.34 | 1.32 | -2.12 | 2.66 | 0.54 | 131.5 | 0.07507 | Non-Toxin | 99.83% (580/581) | Profilin |
| 303 | A*30:02 | 80 | 88 | 9 | YAMELLTGY | 1.34 | 1.32 | -2.21 | 2.66 | 0.44 | 163.3 | 0.07507 | Non-Toxin | 99.83% (580/581) | Profilin |
| 304 | B*58:01 | 80 | 88 | 9 | YAMELLTGY | 1.34 | 1.32 | -2.3 | 2.66 | 0.36 | 200.1 | 0.07507 | Non-Toxin | 99.83% (580/581) | Profilin |
| 305 | A*01:01 | 70 | 80 | 11 | YTNSLMDENTY | 1.61 | 1.23 | -1.87 | 2.85 | 0.98 | 74.3 | -0.2173 | Non-Toxin | 99.83% (580/581) | Profilin |

| # | Allele | Start | End | Peptide Length | Peptide | Proteasome Score | TAP Score | MHC Score | Processing Score | Total Score | MHC IC50[nM] | Immunogenicity | Toxicity | Conservancy | Protein |
|---|---|---|---|---|---|---|---|---|---|---|---|---|---|---|---|
| 306 | A*23:01 | 13 | 22 | 10 | DRYARTIFNF | 1.31 | 1.06 | -1.1 | 2.37 | 1.27 | 12.6 | 0.34667 | Non-Toxin | 100.00% (9/9) | Intermediate transcription factor 3 |
| 307 | A*24:02 | 13 | 22 | 10 | DRYARTIFNF | 1.31 | 1.06 | -1.18 | 2.37 | 1.19 | 15.2 | 0.34667 | Non-Toxin | 100.00% (9/9) | Intermediate transcription factor 3 |
| 308 | A*68:02 | 260 | 268 | 9 | EVANVVNHV | 1.01 | 0.16 | -0.45 | 1.17 | 0.72 | 2.8 | 0.09869 | Non-Toxin | 100.00% (9/9) | Intermediate transcription factor 3 |
| 309 | A*68:02 | 260 | 269 | 10 | EVANVVNHVL | 1.68 | 0.44 | -1.45 | 2.12 | 0.67 | 28.3 | 0.13192 | Non-Toxin | 100.00% (9/9) | Intermediate transcription factor 3 |
| 310 | B*15:01 | 304 | 313 | 10 | FIKSLDHTVF | 1.39 | 1.12 | -1.47 | 2.51 | 1.05 | 29.2 | -0.13834 | Non-Toxin | 22.22% (2/9) | Intermediate transcription factor 3 |
| 311 | A*68:01 | 5 | 14 | 10 | FTFLHEIEDR | 0.9 | 0.66 | -0.9 | 1.56 | 0.66 | 7.9 | 0.37858 | Non-Toxin | 100.00% (9/9) | Intermediate transcription factor 3 |
| 312 | A*23:01 | 239 | 249 | 11 | FYVYYNLFLLF | 1.43 | 1.23 | -1.53 | 2.66 | 1.13 | 33.9 | 0.08334 | Non-Toxin | 100.00% (9/9) | Intermediate transcription factor 3 |
| 313 | B*35:01 | 57 | 65 | 9 | HPAIKKLVY | 1.67 | 1.11 | -1.51 | 2.78 | 1.28 | 32.2 | -0.25162 | Non-Toxin | 100.00% (9/9) | Intermediate transcription factor 3 |
| 314 | B*15:01 | 74 | 83 | 10 | IINQNTYPVF | 1.7 | 1.11 | -1.79 | 2.81 | 1.03 | 61.2 | -0.07588 | Non-Toxin | 100.00% (9/9) | Intermediate transcription factor 3 |
| 315 | A*02:01 | 232 | 241 | 10 | INMIDKLFYV | 1.06 | 0.1 | -0.43 | 1.17 | 0.74 | 2.7 | -0.02528 | Non-Toxin | 100.00% (9/9) | Intermediate transcription factor 3 |
| 316 | A*02:03 | 232 | 241 | 10 | INMIDKLFYV | 1.06 | 0.1 | -0.51 | 1.17 | 0.66 | 3.2 | -0.02528 | Non-Toxin | 100.00% (9/9) | Intermediate transcription factor 3 |
| 317 | A*02:06 | 232 | 241 | 10 | INMIDKLFYV | 1.06 | 0.1 | -0.56 | 1.17 | 0.61 | 3.6 | -0.02528 | Non-Toxin | 100.00% (9/9) | Intermediate transcription factor 3 |
| 318 | A*02:01 | 232 | 242 | 11 | INMIDKLFYVY | 1.59 | 1.24 | -1.41 | 2.83 | 1.41 | 26 | 0.00892 | Non-Toxin | 100.00% (9/9) | Intermediate transcription factor 3 |
| 319 | A*02:03 | 232 | 242 | 11 | INMIDKLFYVY | 1.59 | 1.24 | -1.76 | 2.83 | 1.06 | 57.9 | 0.00892 | Non-Toxin | 100.00% (9/9) | Intermediate transcription factor 3 |
| 320 | A*02:06 | 232 | 242 | 11 | INMIDKLFYVY | 1.59 | 1.24 | -1.76 | 2.83 | 1.06 | 58.2 | 0.00892 | Non-Toxin | 100.00% (9/9) | Intermediate transcription factor 3 |
| 321 | A*01:01 | 25 | 34 | 10 | ISCDEIGDIY | 1.31 | 1.32 | -1.47 | 2.63 | 1.16 | 29.4 | 0.3603 | Non-Toxin | 100.00% (9/9) | Intermediate transcription factor 3 |
| 322 | B*58:01 | 104 | 112 | 9 | ISSRTVEIF | 1.32 | 1.19 | -1.4 | 2.51 | 1.11 | 25 | 0.2373 | Non-Toxin | 100.00% (9/9) | Intermediate transcription factor 3 |
| 323 | A*32:01 | 296 | 304 | 9 | KIGMFNLTF | 1.17 | 1.09 | -1.07 | 2.26 | 1.19 | 11.7 | -0.04447 | Non-Toxin | 100.00% (9/9) | Intermediate transcription factor 3 |
| 324 | A*32:01 | 237 | 245 | 9 | KLFYVYYNL | 1.61 | 0.46 | -0.77 | 2.07 | 1.29 | 5.9 | 0.0641 | Non-Toxin | 100.00% (9/9) | Intermediate transcription factor 3 |
| 325 | A*02:01 | 237 | 245 | 9 | KLFYVYYNL | 1.61 | 0.46 | -1.1 | 2.07 | 0.97 | 12.5 | 0.0641 | Non-Toxin | 100.00% (9/9) | Intermediate transcription factor 3 |
| 326 | A*23:01 | 237 | 246 | 10 | KLFYVYYNLF | 1.13 | 1.14 | -1.22 | 2.27 | 1.05 | 16.7 | 0.05546 | Non-Toxin | 100.00% (9/9) | Intermediate transcription factor 3 |
| 327 | A*32:01 | 237 | 246 | 10 | KLFYVYYNLF | 1.13 | 1.14 | -1.31 | 2.27 | 0.97 | 20.2 | 0.05546 | Non-Toxin | 100.00% (9/9) | Intermediate transcription factor 3 |
| 328 | A*02:03 | 283 | 292 | 10 | KMAEHDVYGV | 1.13 | 0.29 | -0.52 | 1.41 | 0.9 | 3.3 | 0.22209 | Non-Toxin | 100.00% (9/9) | Intermediate transcription factor 3 |
| 329 | A*02:01 | 283 | 292 | 10 | KMAEHDVYGV | 1.13 | 0.29 | -0.83 | 1.41 | 0.59 | 6.7 | 0.22209 | Non-Toxin | 100.00% (9/9) | Intermediate transcription factor 3 |
| 330 | A*23:01 | 238 | 246 | 9 | LFYVYYNLF | 1.13 | 1.3 | -1.36 | 2.43 | 1.06 | 23.1 | 0.02132 | Non-Toxin | 100.00% (9/9) | Intermediate transcription factor 3 |
| 331 | B*58:01 | 270 | 279 | 10 | MATALDEKLF | 1.35 | 1.15 | -1.41 | 2.5 | 1.09 | 25.7 | -0.03146 | Non-Toxin | 100.00% (9/9) | Intermediate transcription factor 3 |
| 332 | B*35:01 | 234 | 242 | 9 | MIDKLFYVY | 1.59 | 1.33 | -1.26 | 2.92 | 1.66 | 18 | -0.0894 | Non-Toxin | 100.00% (9/9) | Intermediate transcription factor 3 |
| 333 | A*01:01 | 234 | 242 | 9 | MIDKLFYVY | 1.59 | 1.33 | -1.52 | 2.92 | 1.4 | 32.9 | -0.0894 | Non-Toxin | 100.00% (9/9) | Intermediate transcription factor 3 |
| 334 | A*30:02 | 234 | 242 | 9 | MIDKLFYVY | 1.59 | 1.33 | -2.09 | 2.92 | 0.83 | 122 | -0.0894 | Non-Toxin | 100.00% (9/9) | Intermediate transcription factor 3 |
| 335 | A*01:01 | 234 | 243 | 10 | MIDKLFYVYY | 1.5 | 1.33 | -1.57 | 2.83 | 1.26 | 37 | -0.0774 | Non-Toxin | 100.00% (9/9) | Intermediate transcription factor 3 |
| 336 | B*08:01 | 355 | 363 | 9 | MMKERSTIL | 1.48 | 0.42 | -0.69 | 1.9 | 1.21 | 4.9 | 0.03594 | Non-Toxin | 100.00% (9/9) | Intermediate transcription factor 3 |
| 337 | A*02:01 | 3 | 11 | 9 | NLFTFLHEI | 1.27 | 0.24 | -0.63 | 1.52 | 0.89 | 4.3 | 0.26642 | Non-Toxin | 100.00% (9/9) | Intermediate transcription factor 3 |
| 338 | A*02:03 | 3 | 11 | 9 | NLFTFLHEI | 1.27 | 0.24 | -0.72 | 1.52 | 0.8 | 5.2 | 0.26642 | Non-Toxin | 100.00% (9/9) | Intermediate transcription factor 3 |
| 339 | A*02:06 | 3 | 11 | 9 | NLFTFLHEI | 1.27 | 0.24 | -0.84 | 1.52 | 0.68 | 6.9 | 0.26642 | Non-Toxin | 100.00% (9/9) | Intermediate transcription factor 3 |
| 340 | A*02:01 | 233 | 241 | 9 | NMIDKLFYV | 1.06 | 0.24 | -0.2 | 1.3 | 1.1 | 1.6 | -0.05828 | Non-Toxin | 100.00% (9/9) | Intermediate transcription factor 3 |
| 341 | A*02:06 | 233 | 241 | 9 | NMIDKLFYV | 1.06 | 0.24 | -0.23 | 1.3 | 1.07 | 1.7 | -0.05828 | Non-Toxin | 100.00% (9/9) | Intermediate transcription factor 3 |
| 342 | A*02:03 | 233 | 241 | 9 | NMIDKLFYV | 1.06 | 0.24 | -0.28 | 1.3 | 1.02 | 1.9 | -0.05828 | Non-Toxin | 100.00% (9/9) | Intermediate transcription factor 3 |
| 343 | A*02:01 | 233 | 242 | 10 | NMIDKLFYVY | 1.59 | 1.37 | -0.99 | 2.96 | 1.97 | 9.8 | -0.02408 | Non-Toxin | 100.00% (9/9) | Intermediate transcription factor 3 |
| 344 | A*02:03 | 233 | 242 | 10 | NMIDKLFYVY | 1.59 | 1.37 | -1.1 | 2.96 | 1.86 | 12.7 | -0.02408 | Non-Toxin | 100.00% (9/9) | Intermediate transcription factor 3 |
| 345 | A*02:06 | 233 | 242 | 10 | NMIDKLFYVY | 1.59 | 1.37 | -1.16 | 2.96 | 1.8 | 14.5 | -0.02408 | Non-Toxin | 100.00% (9/9) | Intermediate transcription factor 3 |
| 346 | B*15:01 | 233 | 242 | 10 | NMIDKLFYVY | 1.59 | 1.37 | -2.05 | 2.96 | 0.91 | 112.7 | -0.02408 | Non-Toxin | 100.00% (9/9) | Intermediate transcription factor 3 |
| 347 | A*02:01 | 233 | 243 | 11 | NMIDKLFYVYY | 1.5 | 1.37 | -2.22 | 2.87 | 0.66 | 164.1 | -0.01208 | Non-Toxin | 100.00% (9/9) | Intermediate transcription factor 3 |
| 348 | A*68:01 | 198 | 207 | 10 | QTIEIIFTNR | 0.97 | 0.69 | -0.83 | 1.66 | 0.83 | 6.8 | 0.54233 | Non-Toxin | 100.00% (9/9) | Intermediate transcription factor 3 |
| 349 | A*30:02 | 337 | 345 | 9 | RSLEDCTNY | 1.29 | 1.42 | -1.77 | 2.71 | 0.94 | 59.3 | 0.09698 | Non-Toxin | 100.00% (9/9) | Intermediate transcription factor 3 |
| 350 | B*58:01 | 337 | 345 | 9 | RSLEDCTNY | 1.29 | 1.42 | -2.07 | 2.71 | 0.64 | 117.2 | 0.09698 | Non-Toxin | 100.00% (9/9) | Intermediate transcription factor 3 |
| 351 | A*23:01 | 14 | 22 | 9 | RYARTIFNF | 1.31 | 1.27 | -0.53 | 2.58 | 2.05 | 3.4 | 0.32288 | Non-Toxin | 100.00% (9/9) | Intermediate transcription factor 3 |

| | | | | | | | | | | | | | | | |
|---|---|---|---|---|---|---|---|---|---|---|---|---|---|---|---|
| 352 | A*24:02 | 14 | 22 | 9 | RYARTIFNF | 1.31 | 1.27 | -0.54 | 2.58 | 2.04 | 3.5 | 0.32288 | Non-Toxin | 100.00% (9/9) | Intermediate transcription factor 3 |
| 353 | B*44:03 | 43 | 52 | 10 | SEDMFDNIVY | 1.47 | 1.18 | -1.7 | 2.65 | 0.95 | 50.2 | 0.09645 | Non-Toxin | 100.00% (9/9) | Intermediate transcription factor 3 |
| 354 | B*44:02 | 43 | 52 | 10 | SEDMFDNIVY | 1.47 | 1.18 | -1.87 | 2.65 | 0.79 | 73.6 | 0.09645 | Non-Toxin | 100.00% (9/9) | Intermediate transcription factor 3 |
| 355 | B*44:02 | 114 | 122 | 9 | SEKSSLVSY | 1.19 | 1.16 | -1.77 | 2.35 | 0.58 | 59 | -0.46983 | Non-Toxin | 100.00% (9/9) | Intermediate transcription factor 3 |
| 356 | B*15:01 | 87 | 95 | 9 | SQVKCCHYF | 1.29 | 1.21 | -1.28 | 2.5 | 1.22 | 19.2 | -0.28171 | Non-Toxin | 100.00% (9/9) | Intermediate transcription factor 3 |
| 357 | B*15:01 | 86 | 95 | 10 | SSQVKCCHYF | 1.29 | 1.07 | -1.76 | 2.36 | 0.6 | 57.6 | -0.28417 | Non-Toxin | 100.00% (9/9) | Intermediate transcription factor 3 |
| 358 | A*23:01 | 241 | 249 | 9 | VYYNLFLLF | 1.43 | 1.3 | -0.57 | 2.73 | 2.16 | 3.7 | 0.07585 | Non-Toxin | 100.00% (9/9) | Intermediate transcription factor 3 |
| 359 | A*24:02 | 241 | 249 | 9 | VYYNLFLLF | 1.43 | 1.3 | -0.94 | 2.73 | 1.79 | 8.7 | 0.07585 | Non-Toxin | 100.00% (9/9) | Intermediate transcription factor 3 |
| 360 | A*02:06 | 240 | 248 | 9 | YVYYNLFLL | 1.32 | 0.5 | -0.76 | 1.82 | 1.06 | 5.8 | 0.07066 | Non-Toxin | 100.00% (9/9) | Intermediate transcription factor 3 |
| 361 | A*02:01 | 240 | 248 | 9 | YVYYNLFLL | 1.32 | 0.5 | -0.86 | 1.82 | 0.96 | 7.3 | 0.07066 | Non-Toxin | 100.00% (9/9) | Intermediate transcription factor 3 |
| 362 | A*68:02 | 240 | 248 | 9 | YVYYNLFLL | 1.32 | 0.5 | -1.21 | 1.82 | 0.61 | 16.1 | 0.07066 | Non-Toxin | 100.00% (9/9) | Intermediate transcription factor 3 |
| 363 | A*02:03 | 240 | 248 | 9 | YVYYNLFLL | 1.32 | 0.5 | -1.25 | 1.82 | 0.57 | 17.8 | 0.07066 | Non-Toxin | 100.00% (9/9) | Intermediate transcription factor 3 |
| 364 | A*23:01 | 240 | 249 | 10 | YVYYNLFLLF | 1.43 | 1.19 | -0.72 | 2.61 | 1.9 | 5.2 | 0.07234 | Non-Toxin | 100.00% (9/9) | Intermediate transcription factor 3 |
| 365 | A*24:02 | 240 | 249 | 10 | YVYYNLFLLF | 1.43 | 1.19 | -1.2 | 2.61 | 1.41 | 16 | 0.07234 | Non-Toxin | 100.00% (9/9) | Intermediate transcription factor 3 |
| 366 | B*15:01 | 240 | 249 | 10 | YVYYNLFLLF | 1.43 | 1.19 | -1.92 | 2.61 | 0.7 | 82.9 | 0.07234 | Non-Toxin | 100.00% (9/9) | Intermediate transcription factor 3 |
| 367 | A*23:01 | 242 | 249 | 8 | YYNLFLLF | 1.43 | 1.19 | -1.23 | 2.61 | 1.38 | 17.1 | 0.08094 | Non-Toxin | 100.00% (9/9) | Intermediate transcription factor 3 |
| 368 | A*24:02 | 242 | 249 | 8 | YYNLFLLF | 1.43 | 1.19 | -1.66 | 2.61 | 0.95 | 45.7 | 0.08094 | Non-Toxin | 100.00% (9/9) | Intermediate transcription factor 3 |

| # | Allele | Start | End | Peptide Length | Peptide | Proteasome Score | TAP Score | MHC Score | Processing Score | Total Score | MHC IC50[nM] | Immunogenicity | Toxicity | Conservancy | Protein |
|---|---|---|---|---|---|---|---|---|---|---|---|---|---|---|---|
| 369 | A*31:01 | 16 | 24 | 9 | ALRNLQLHR | 1.37 | 0.74 | -1.3 | 2.11 | 0.81 | 19.9 | -0.10001 | Non-Toxin | 100.00% (1/1) | Probable host range protein 2 |
| 370 | B*35:01 | 106 | 114 | 9 | ELFKHYYPY | 0.87 | 1.3 | -1.64 | 2.17 | 0.52 | 43.9 | -0.16058 | Non-Toxin | 100.00% (1/1) | Probable host range protein 2 |
| 371 | A*30:02 | 115 | 124 | 10 | ISLNMISKKY | 1.5 | 1.28 | -1.98 | 2.78 | 0.8 | 96 | -0.51524 | Non-Toxin | 100.00% (1/1) | Probable host range protein 2 |
| 372 | A*30:02 | 59 | 68 | 10 | KGLTVFANNY | 1.43 | 1.23 | -2.07 | 2.66 | 0.58 | 118.2 | 0.21725 | Non-Toxin | 100.00% (1/1) | Probable host range protein 2 |
| 373 | A*30:02 | 31 | 39 | 9 | KLKIISNDY | 1.08 | 1.32 | -1.71 | 2.4 | 0.69 | 51.8 | 0.04529 | Non-Toxin | 100.00% (1/1) | Probable host range protein 2 |
| 374 | A*31:01 | 41 | 49 | 9 | KLKLRFVIR | 1.29 | 0.74 | -1.13 | 2.03 | 0.9 | 13.4 | 0.19204 | Non-Toxin | 100.00% (1/1) | Probable host range protein 2 |
| 375 | A*32:01 | 71 | 79 | 9 | KVNKVDYTL | 1.59 | 0.5 | -1.21 | 2.09 | 0.89 | 16.2 | -0.13846 | Non-Toxin | 100.00% (1/1) | Probable host range protein 2 |
| 376 | A*30:02 | 71 | 80 | 10 | KVNKVDYTLY | 1.27 | 1.36 | -1.49 | 2.63 | 1.13 | 31.1 | -0.1345 | Non-Toxin | 100.00% (1/1) | Probable host range protein 2 |
| 377 | A*30:02 | 71 | 81 | 11 | KVNKVDYTLYY | 1.48 | 1.36 | -1.89 | 2.83 | 0.95 | 76.9 | -0.13588 | Non-Toxin | 100.00% (1/1) | Probable host range protein 2 |
| 378 | A*11:01 | 71 | 81 | 11 | KVNKVDYTLYY | 1.48 | 1.36 | -2.31 | 2.83 | 0.52 | 205.2 | -0.13588 | Non-Toxin | 100.00% (1/1) | Probable host range protein 2 |
| 379 | A*30:02 | 116 | 124 | 9 | SLNMISKKY | 1.5 | 1.26 | -2.19 | 2.77 | 0.58 | 153.3 | -0.51293 | Non-Toxin | 100.00% (1/1) | Probable host range protein 2 |
| 380 | A*68:02 | 62 | 70 | 9 | TVFANNYAV | 1.1 | 0.26 | -0.61 | 1.36 | 0.74 | 4.1 | 0.08472 | Non-Toxin | 100.00% (1/1) | Probable host range protein 2 |
| 381 | A*02:01 | 82 | 90 | 9 | VIYEAVIHL | 1.57 | 0.56 | -1.51 | 2.13 | 0.62 | 32.4 | 0.30773 | Non-Toxin | 100.00% (1/1) | Probable host range protein 2 |
| 382 | A*02:03 | 82 | 90 | 9 | VIYEAVIHL | 1.57 | 0.56 | -1.56 | 2.13 | 0.57 | 36.2 | 0.30773 | Non-Toxin | 100.00% (1/1) | Probable host range protein 2 |
| 383 | A*02:06 | 82 | 90 | 9 | VIYEAVIHL | 1.57 | 0.56 | -1.62 | 2.13 | 0.52 | 41.3 | 0.30773 | Non-Toxin | 100.00% (1/1) | Probable host range protein 2 |
| 384 | A*30:02 | 82 | 91 | 10 | VIYEAVIHLY | 1.16 | 1.41 | -1.97 | 2.57 | 0.6 | 94.4 | 0.32395 | Non-Toxin | 100.00% (1/1) | Probable host range protein 2 |
| 385 | B*15:01 | 82 | 91 | 10 | VIYEAVIHLY | 1.16 | 1.41 | -2.03 | 2.57 | 0.55 | 106.5 | 0.32395 | Non-Toxin | 100.00% (1/1) | Probable host range protein 2 |
| 386 | A*03:01 | 82 | 91 | 10 | VIYEAVIHLY | 1.16 | 1.41 | -2.07 | 2.57 | 0.51 | 116.7 | 0.32395 | Non-Toxin | 100.00% (1/1) | Probable host range protein 2 |
| 387 | B*35:01 | 135 | 143 | 9 | YIEHPLIPY | 1.36 | 1.19 | -1.83 | 2.55 | 0.72 | 68.3 | 0.14965 | Non-Toxin | 100.00% (1/1) | Probable host range protein 2 |
| 388 | A*01:01 | 100 | 108 | 9 | YSDDENELF | 1.46 | 1.04 | -1.99 | 2.5 | 0.5 | 98.8 | 0.19895 | Non-Toxin | 100.00% (1/1) | Probable host range protein 2 |
| 389 | A*02:06 | 81 | 90 | 10 | YVIYEAVIHL | 1.57 | 0.48 | -1.33 | 2.05 | 0.72 | 21.4 | 0.34516 | Non-Toxin | 100.00% (1/1) | Probable host range protein 2 |
| 390 | A*02:03 | 81 | 90 | 10 | YVIYEAVIHL | 1.57 | 0.48 | -1.38 | 2.05 | 0.67 | 23.9 | 0.34516 | Non-Toxin | 100.00% (1/1) | Probable host range protein 2 |
| 391 | A*02:01 | 81 | 90 | 10 | YVIYEAVIHL | 1.57 | 0.48 | -1.49 | 2.05 | 0.56 | 31.1 | 0.34516 | Non-Toxin | 100.00% (1/1) | Probable host range protein 2 |

**Supplementary Table S2:** High Scoring CTL epitopes-HLA allele pairs screened from eleven proteins of Monkeypox virus by the "MHC-I Binding Predictions" tool of IEDB. Out of these epitopes, the top scoring epitopes were used to design Multi-epitope vaccine.

| # | allele | start | end | length | peptide | score | percentile_rank | Immunogenicity | Toxicity | Conservancy | Protein |
|---|---|---|---|---|---|---|---|---|---|---|---|
| 1 | A*11:01 | 187 | 196 | 10 | AVLPIPNLVK | 0.953745 | 0.01 | 0.11271 | Non-Toxin | 77.78% (7/9) | Poxin-Schlafen |
| 2 | A*26:01 | 484 | 492 | 9 | DTSGSMKKY | 0.947791 | 0.01 | -0.654 | Non-Toxin | 77.78% (7/9) | Poxin-Schlafen |
| 3 | A*68:02 | 241 | 249 | 9 | ESIHHLWSV | 0.966138 | 0.01 | 0.18709 | Non-Toxin | 66.67% (6/9) | Poxin-Schlafen |
| 4 | B*15:01 | 425 | 433 | 9 | IQKLPVVHF | 0.967539 | 0.01 | 0.00064 | Non-Toxin | 66.67% (6/9) | Poxin-Schlafen |
| 5 | A*23:01 | 423 | 433 | 11 | KYIQKLPVVHF | 0.949306 | 0.01 | -0.21236 | Non-Toxin | 66.67% (6/9) | Poxin-Schlafen |
| 6 | B*51:01 | 189 | 197 | 9 | LPIPNLVKV | 0.978854 | 0.01 | -0.07586 | Non-Toxin | 77.78% (7/9) | Poxin-Schlafen |
| 7 | B*15:01 | 376 | 385 | 10 | QQLPSILSSF | 0.925319 | 0.01 | -0.29298 | Non-Toxin | 100.00% (9/9) | Poxin-Schlafen |

| | | | | | | | | | | | |
|---|---|---|---|---|---|---|---|---|---|---|---|
| 8 | A*32:01 | 239 | 247 | 9 | RLESIHHLW | 0.903663 | 0.01 | 0.0469 | Non-Toxin | 66.67% (6/9) | Poxin-Schlafen |
| 9 | A*23:01 | 58 | 67 | 10 | RYIGALLPMF | 0.931298 | 0.01 | -0.0178 | Non-Toxin | 77.78% (7/9) | Poxin-Schlafen |
| 10 | A*11:01 | 32 | 40 | 9 | SVVSVYNKK | 0.931494 | 0.01 | -0.24781 | Non-Toxin | 66.67% (6/9) | Poxin-Schlafen |
| 11 | A*02:06 | 249 | 257 | 9 | VVYDHLNVV | 0.962428 | 0.01 | 0.06084 | Non-Toxin | 66.67% (6/9) | Poxin-Schlafen |
| 12 | B*15:01 | 148 | 157 | 10 | ALKPGPIIDY | 0.919805 | 0.02 | 0.1916 | Non-Toxin | 66.67% (6/9) | Poxin-Schlafen |
| 13 | A*03:01 | 187 | 196 | 10 | AVLPIPNLVK | 0.940371 | 0.02 | 0.11271 | Non-Toxin | 77.78% (7/9) | Poxin-Schlafen |
| 14 | A*24:02 | 9 | 17 | 9 | GYDENLHAF | 0.910205 | 0.02 | 0.14137 | Non-Toxin | 77.78% (7/9) | Poxin-Schlafen |
| 15 | B*51:01 | 191 | 199 | 9 | IPNLVKVKV | 0.915243 | 0.02 | -0.26722 | Non-Toxin | 66.67% (6/9) | Poxin-Schlafen |
| 16 | A*24:02 | 423 | 433 | 11 | KYIQKLPVVHF | 0.93425 | 0.02 | -0.21236 | Non-Toxin | 66.67% (6/9) | Poxin-Schlafen |
| 17 | B*35:01 | 64 | 72 | 9 | LPMFECNEY | 0.945552 | 0.02 | 0.16059 | Non-Toxin | 77.78% (7/9) | Poxin-Schlafen |
| 18 | B*15:01 | 346 | 354 | 9 | LQAGESVKF | 0.908634 | 0.02 | -0.10259 | Non-Toxin | 88.89% (8/9) | Poxin-Schlafen |
| 19 | A*31:01 | 335 | 343 | 9 | RLAEEFFNR | 0.907069 | 0.02 | 0.41617 | Non-Toxin | 88.89% (8/9) | Poxin-Schlafen |
| 20 | A*24:02 | 58 | 67 | 10 | RYIGALLPMF | 0.922624 | 0.02 | -0.0178 | Non-Toxin | 77.78% (7/9) | Poxin-Schlafen |
| 21 | B*08:01 | 357 | 365 | 9 | SINVKHTSV | 0.880113 | 0.02 | -0.20401 | Non-Toxin | 100.00% (9/9) | Poxin-Schlafen |
| 22 | B*08:01 | 138 | 146 | 9 | TIKEKAKEM | 0.888431 | 0.02 | -0.26592 | Non-Toxin | 77.78% (7/9) | Poxin-Schlafen |
| 23 | A*02:03 | 249 | 257 | 9 | VVYDHLNVV | 0.954048 | 0.02 | 0.06084 | Non-Toxin | 66.67% (6/9) | Poxin-Schlafen |
| 24 | A*02:06 | 187 | 195 | 9 | AVLPIPNLV | 0.931685 | 0.03 | 0.09246 | Non-Toxin | 77.78% (7/9) | Poxin-Schlafen |
| 25 | B*53:01 | 152 | 160 | 9 | GPIIDYHVW | 0.881278 | 0.03 | 0.24666 | Non-Toxin | 77.78% (7/9) | Poxin-Schlafen |
| 26 | A*01:01 | 293 | 302 | 10 | QMDSMEALEY | 0.910938 | 0.03 | -0.1468 | Non-Toxin | 88.89% (8/9) | Poxin-Schlafen |
| 27 | B*07:02 | 313 | 321 | 9 | SPRPELQKF | 0.949859 | 0.03 | -0.13106 | Non-Toxin | 66.67% (6/9) | Poxin-Schlafen |
| 28 | A*26:01 | 23 | 31 | 9 | TVANDVRKY | 0.876327 | 0.03 | -0.01567 | Non-Toxin | 77.78% (7/9) | Poxin-Schlafen |
| 29 | A*01:01 | 482 | 492 | 11 | YMDTSGSMKKY | 0.909264 | 0.03 | -0.71624 | Non-Toxin | 77.78% (7/9) | Poxin-Schlafen |
| 30 | B*44:03 | 495 | 503 | 9 | DEWVSHIKF | 0.904805 | 0.04 | -0.03089 | Non-Toxin | 77.78% (7/9) | Poxin-Schlafen |
| 31 | B*58:01 | 239 | 247 | 9 | RLESIHHLW | 0.946579 | 0.04 | 0.0469 | Non-Toxin | 66.67% (6/9) | Poxin-Schlafen |
| 32 | B*44:03 | 344 | 354 | 11 | SELQAGESVKF | 0.891153 | 0.04 | -0.19845 | Non-Toxin | 88.89% (8/9) | Poxin-Schlafen |
| 33 | B*35:01 | 313 | 321 | 9 | SPRPELQKF | 0.904377 | 0.04 | -0.13106 | Non-Toxin | 66.67% (6/9) | Poxin-Schlafen |
| **#** | **allele** | **start** | **end** | **length** | **peptide** | **score** | **percentile_rank** | **Immunogenicity** | **Toxicity** | **Conservancy** | **Protein** |
| 34 | B*15:01 | 44 | 52 | 9 | KLNDDTQVY | 0.949449 | 0.01 | 0.00472 | Non-Toxin | 100.00% (7/7) | Cell surface-binding protein |
| 35 | A*30:02 | 44 | 52 | 9 | KLNDDTQVY | 0.857415 | 0.01 | 0.00472 | Non-Toxin | 100.00% (7/7) | Cell surface-binding protein |
| 36 | A*24:02 | 28 | 36 | 9 | KYIEGNKTF | 0.980752 | 0.01 | 0.01154 | Non-Toxin | 100.00% (7/7) | Cell surface-binding protein |
| 37 | A*23:01 | 28 | 36 | 9 | KYIEGNKTF | 0.976944 | 0.01 | 0.01154 | Non-Toxin | 100.00% (7/7) | Cell surface-binding protein |
| 38 | B*15:01 | 20 | 28 | 9 | RLKTLDIHY | 0.960197 | 0.01 | 0.11036 | Non-Toxin | 100.00% (7/7) | Cell surface-binding protein |
| 39 | A*11:01 | 4 | 12 | 9 | STIHIYWGK | 0.921271 | 0.01 | 0.40861 | Non-Toxin | 100.00% (7/7) | Cell surface-binding protein |
| 40 | B*08:01 | 14 | 22 | 9 | DQLSKFRTL | 0.873398 | 0.02 | -0.20351 | Non-Toxin | 100.00% (7/7) | Cell surface-binding protein |
| 41 | A*01:01 | 44 | 53 | 10 | KLNDDTQVYY | 0.927876 | 0.02 | 0.00326 | Non-Toxin | 100.00% (7/7) | Cell surface-binding protein |
| 42 | A*30:02 | 44 | 53 | 10 | KLNDDTQVYY | 0.825858 | 0.02 | 0.00326 | Non-Toxin | 100.00% (7/7) | Cell surface-binding protein |
| 43 | B*15:01 | 1 | 10 | 10 | LQVSDHKNVY | 0.900017 | 0.02 | -0.28431 | Non-Toxin | 100.00% (7/7) | Cell surface-binding protein |
| 44 | A*01:01 | 45 | 53 | 9 | LNDDTQVYY | 0.898587 | 0.03 | -0.00904 | Non-Toxin | 100.00% (7/7) | Cell surface-binding protein |
| 45 | A*30:02 | 20 | 28 | 9 | RLKTLDIHY | 0.795893 | 0.03 | 0.11036 | Non-Toxin | 100.00% (7/7) | Cell surface-binding protein |
| 46 | A*30:02 | 2 | 10 | 9 | TTSPVRENY | 0.791927 | 0.03 | 0.10478 | Non-Toxin | 100.00% (7/7) | Cell surface-binding protein |
| 47 | A*24:02 | 26 | 36 | 11 | YQKYIEGNKTF | 0.894951 | 0.03 | 0.02097 | Non-Toxin | 100.00% (7/7) | Cell surface-binding protein |
| 48 | A*23:01 | 26 | 36 | 11 | YQKYIEGNKTF | 0.867888 | 0.03 | 0.02097 | Non-Toxin | 100.00% (7/7) | Cell surface-binding protein |
| 49 | B*58:01 | 27 | 36 | 10 | YSGEINLVHW | 0.96297 | 0.03 | 0.27835 | Non-Toxin | 100.00% (7/7) | Cell surface-binding protein |
| 50 | A*30:02 | 1 | 9 | 9 | YVLSTIHIY | 0.759971 | 0.03 | 0.09807 | Non-Toxin | 100.00% (7/7) | Cell surface-binding protein |
| 51 | B*15:01 | 44 | 53 | 10 | KLNDDTQVYY | 0.848004 | 0.04 | 0.00326 | Non-Toxin | 100.00% (7/7) | Cell surface-binding protein |
| 52 | A*02:06 | 42 | 50 | 9 | STLDYFTYL | 0.902861 | 0.04 | 0.15592 | Non-Toxin | 100.00% (7/7) | Cell surface-binding protein |
| 53 | B*57:01 | 27 | 36 | 10 | YSGEINLVHW | 0.966675 | 0.04 | 0.27835 | Non-Toxin | 100.00% (7/7) | Cell surface-binding protein |
| 54 | B*35:01 | 1 | 9 | 9 | YVLSTIHIY | 0.904252 | 0.04 | 0.09807 | Non-Toxin | 100.00% (7/7) | Cell surface-binding protein |
| 55 | B*44:03 | 29 | 36 | 8 | GEINLVHW | 0.864485 | 0.05 | 0.09205 | Non-Toxin | 100.00% (7/7) | Cell surface-binding protein |
| 56 | B*35:01 | 31 | 39 | 9 | KPHYITENY | 0.879993 | 0.05 | 0.25364 | Non-Toxin | 100.00% (7/7) | Cell surface-binding protein |
| 57 | A*01:01 | 2 | 10 | 9 | TTSPVRENY | 0.851515 | 0.05 | 0.10478 | Non-Toxin | 100.00% (7/7) | Cell surface-binding protein |
| 58 | A*26:01 | 2 | 10 | 9 | TTSPVRENY | 0.779031 | 0.05 | 0.10478 | Non-Toxin | 100.00% (7/7) | Cell surface-binding protein |
| 59 | A*23:01 | 44 | 52 | 9 | VFILTAILF | 0.76673 | 0.05 | 0.21251 | Non-Toxin | 100.00% (7/7) | Cell surface-binding protein |

| | | | | | | | | | | | |
|---|---|---|---|---|---|---|---|---|---|---|---|
| 60 | A*26:01 | 1 | 9 | 9 | YVLSTIHIY | 0.768904 | 0.05 | 0.09807 | Non-Toxin | 100.00% (7/7) | Cell surface-binding protein |
| 61 | A*68:01 | 11 | 20 | 10 | ETKKAISDTR | 0.915876 | 0.06 | -0.23363 | Non-Toxin | 14.29% (1/7) | Cell surface-binding protein |
| 62 | A*02:06 | 3 | 11 | 9 | IIFPTPINI | 0.852122 | 0.06 | 0.16274 | Non-Toxin | 100.00% (7/7) | Cell surface-binding protein |
| 63 | A*02:03 | 3 | 11 | 9 | IIFPTPINI | 0.782931 | 0.06 | 0.16274 | Non-Toxin | 100.00% (7/7) | Cell surface-binding protein |
| 64 | B*35:01 | 40 | 49 | 10 | LPSTLDYFTY | 0.856222 | 0.06 | 0.11416 | Non-Toxin | 100.00% (7/7) | Cell surface-binding protein |
| 65 | A*01:01 | 15 | 25 | 11 | LSDLREACFSY | 0.811055 | 0.06 | 0.13343 | Non-Toxin | 100.00% (7/7) | Cell surface-binding protein |
| 66 | B*58:01 | 26 | 34 | 9 | MSAPFDSVF | 0.899602 | 0.06 | 0.02092 | Non-Toxin | 100.00% (7/7) | Cell surface-binding protein |
| 67 | A*01:01 | 26 | 35 | 10 | MSAPFDSVFY | 0.831188 | 0.06 | 0.08465 | Non-Toxin | 100.00% (7/7) | Cell surface-binding protein |
| 68 | A*24:02 | 27 | 36 | 10 | QKYIEGNKTF | 0.806491 | 0.06 | 0.09781 | Non-Toxin | 100.00% (7/7) | Cell surface-binding protein |
| 69 | A*68:01 | 4 | 12 | 9 | STIHIYWGK | 0.920676 | 0.06 | 0.40861 | Non-Toxin | 100.00% (7/7) | Cell surface-binding protein |
| 70 | B*58:01 | 2 | 10 | 9 | VLSTIHIYW | 0.9107 | 0.06 | 0.25557 | Non-Toxin | 100.00% (7/7) | Cell surface-binding protein |
| 71 | B*15:01 | 26 | 36 | 11 | YQKYIEGNKTF | 0.804677 | 0.06 | 0.02097 | Non-Toxin | 100.00% (7/7) | Cell surface-binding protein |
| 72 | B*57:01 | 1 | 10 | 10 | YVLSTIHIYW | 0.948703 | 0.06 | 0.13794 | Non-Toxin | 100.00% (7/7) | Cell surface-binding protein |
| **#** | **allele** | **start** | **end** | **length** | **peptide** | **score** | **percentile_rank** | **Immunogenicity** | **Toxicity** | **Conservancy** | **Protein** |
| 73 | A*26:01 | 153 | 161 | 9 | DIIKILDKY | 0.952028 | 0.01 | -0.16192 | Non-Toxin | 100.00% (19/19) | E3 ubiquitin-protein ligase |
| 74 | A*31:01 | 49 | 57 | 9 | KINPHLANR | 0.984402 | 0.01 | 0.03704 | Non-Toxin | 100.00% (19/19) | E3 ubiquitin-protein ligase |
| 75 | A*30:01 | 228 | 236 | 9 | RFRKITMSK | 0.95173 | 0.01 | -0.27892 | Non-Toxin | 73.68% (14/19) | E3 ubiquitin-protein ligase |
| 76 | B*40:01 | 150 | 158 | 9 | SELDIIKIL | 0.987141 | 0.01 | 0.16936 | Non-Toxin | 100.00% (19/19) | E3 ubiquitin-protein ligase |
| 77 | B*44:03 | 150 | 158 | 9 | SELDIIKIL | 0.974234 | 0.01 | 0.16936 | Non-Toxin | 100.00% (19/19) | E3 ubiquitin-protein ligase |
| 78 | A*01:01 | 157 | 165 | 9 | ILDKYEDMY | 0.941737 | 0.02 | -0.20303 | Non-Toxin | 100.00% (19/19) | E3 ubiquitin-protein ligase |
| 79 | A*11:01 | 41 | 49 | 9 | ITYYINITK | 0.893764 | 0.02 | 0.25359 | Non-Toxin | 100.00% (19/19) | E3 ubiquitin-protein ligase |
| 80 | B*44:02 | 150 | 158 | 9 | SELDIIKIL | 0.935239 | 0.02 | 0.16936 | Non-Toxin | 100.00% (19/19) | E3 ubiquitin-protein ligase |
| 81 | A*01:01 | 12 | 21 | 10 | SIDHVTILQY | 0.932625 | 0.02 | 0.16599 | Non-Toxin | 100.00% (19/19) | E3 ubiquitin-protein ligase |
| 82 | A*32:01 | 123 | 131 | 9 | SILRGLVNW | 0.77592 | 0.02 | 0.1021 | Non-Toxin | 100.00% (19/19) | E3 ubiquitin-protein ligase |
| 83 | A*68:01 | 118 | 126 | 9 | DVIVQSILR | 0.950306 | 0.03 | -0.07795 | Non-Toxin | 100.00% (19/19) | E3 ubiquitin-protein ligase |
| 84 | B*57:01 | 122 | 131 | 10 | QSILRGLVNW | 0.977383 | 0.03 | 0.13606 | Non-Toxin | 100.00% (19/19) | E3 ubiquitin-protein ligase |
| 85 | A*30:01 | 226 | 236 | 11 | RTRFRKITMSK | 0.750405 | 0.03 | -0.10372 | Non-Toxin | 73.68% (14/19) | E3 ubiquitin-protein ligase |
| 86 | A*24:02 | 42 | 50 | 9 | TYYINITKI | 0.884661 | 0.03 | 0.15846 | Non-Toxin | 100.00% (19/19) | E3 ubiquitin-protein ligase |
| 87 | A*33:01 | 118 | 126 | 9 | DVIVQSILR | 0.770918 | 0.04 | -0.07795 | Non-Toxin | 100.00% (19/19) | E3 ubiquitin-protein ligase |
| 88 | B*51:01 | 24 | 32 | 9 | EPNDIRLTV | 0.808159 | 0.04 | 0.21186 | Non-Toxin | 100.00% (19/19) | E3 ubiquitin-protein ligase |
| 89 | A*03:01 | 41 | 49 | 9 | ITYYINITK | 0.888965 | 0.04 | 0.25359 | Non-Toxin | 100.00% (19/19) | E3 ubiquitin-protein ligase |
| 90 | A*03:01 | 49 | 57 | 9 | KINPHLANR | 0.881824 | 0.04 | 0.03704 | Non-Toxin | 100.00% (19/19) | E3 ubiquitin-protein ligase |
| 91 | A*31:01 | 49 | 59 | 11 | KINPHLANRFR | 0.879088 | 0.04 | 0.15153 | Non-Toxin | 100.00% (19/19) | E3 ubiquitin-protein ligase |
| 92 | A*30:01 | 226 | 234 | 9 | RTRFRKITM | 0.721339 | 0.04 | 0.117 | Non-Toxin | 73.68% (14/19) | E3 ubiquitin-protein ligase |
| 93 | A*23:01 | 42 | 50 | 9 | TYYINITKI | 0.85097 | 0.04 | 0.15846 | Non-Toxin | 100.00% (19/19) | E3 ubiquitin-protein ligase |
| 94 | A*02:06 | 111 | 119 | 9 | VVIDWITDV | 0.905217 | 0.04 | 0.45222 | Non-Toxin | 100.00% (19/19) | E3 ubiquitin-protein ligase |
| 95 | A*32:01 | 53 | 61 | 9 | HLANRFRAW | 0.642605 | 0.05 | 0.23333 | Non-Toxin | 100.00% (19/19) | E3 ubiquitin-protein ligase |
| 96 | A*32:01 | 232 | 240 | 9 | ITMSKFYKL | 0.630692 | 0.05 | -0.45239 | Non-Toxin | 100.00% (19/19) | E3 ubiquitin-protein ligase |
| 97 | A*26:01 | 97 | 105 | 9 | NIYGLYIHY | 0.751724 | 0.05 | 0.14984 | Non-Toxin | 100.00% (19/19) | E3 ubiquitin-protein ligase |
| 98 | A*31:01 | 156 | 166 | 11 | KILDKYEDMYR | 0.802952 | 0.06 | -0.22686 | Non-Toxin | 100.00% (19/19) | E3 ubiquitin-protein ligase |
| 99 | B*58:01 | 122 | 131 | 10 | QSILRGLVNW | 0.905181 | 0.06 | 0.13606 | Non-Toxin | 100.00% (19/19) | E3 ubiquitin-protein ligase |
| 100 | A*30:02 | 36 | 44 | 9 | RNINNITYY | 0.680854 | 0.06 | 0.18627 | Non-Toxin | 52.63% (10/19) | E3 ubiquitin-protein ligase |
| 101 | A*30:02 | 11 | 21 | 11 | SSIDHVTILQY | 0.703336 | 0.06 | 0.22326 | Non-Toxin | 100.00% (19/19) | E3 ubiquitin-protein ligase |
| 102 | A*31:01 | 59 | 68 | 10 | RAWKKRIAGR | 0.773522 | 0.07 | -0.1266 | Non-Toxin | 100.00% (19/19) | E3 ubiquitin-protein ligase |
| 103 | B*44:03 | 169 | 177 | 9 | KEKECGICY | 0.795454 | 0.08 | 0.09097 | Non-Toxin | 100.00% (19/19) | E3 ubiquitin-protein ligase |
| 104 | B*57:01 | 123 | 131 | 9 | SILRGLVNW | 0.928323 | 0.08 | 0.1021 | Non-Toxin | 100.00% (19/19) | E3 ubiquitin-protein ligase |
| 105 | B*58:01 | 123 | 131 | 9 | SILRGLVNW | 0.859598 | 0.08 | 0.1021 | Non-Toxin | 100.00% (19/19) | E3 ubiquitin-protein ligase |
| 106 | B*08:01 | 232 | 240 | 9 | ITMSKFYKL | 0.619446 | 0.09 | -0.45239 | Non-Toxin | 100.00% (19/19) | E3 ubiquitin-protein ligase |
| 107 | B*44:02 | 169 | 177 | 9 | KEKECGICY | 0.720011 | 0.09 | 0.09097 | Non-Toxin | 100.00% (19/19) | E3 ubiquitin-protein ligase |
| 108 | A*30:02 | 94 | 102 | 9 | KNKNIYGLY | 0.625291 | 0.09 | 0.07173 | Non-Toxin | 100.00% (19/19) | E3 ubiquitin-protein ligase |
| **#** | **allele** | **start** | **end** | **length** | **peptide** | **score** | **percentile_rank** | **Immunogenicity** | **Toxicity** | **Conservancy** | **Protein** |
| 109 | A*26:01 | 158 | 166 | 9 | EIIGGNDMY | 0.932141 | 0.01 | 0.02033 | Non-Toxin | 100.00% (607/607) | Thymidine kinase |
| 110 | A*23:01 | 85 | 93 | 9 | QFFPDIVEF | 0.96163 | 0.01 | 0.26706 | Non-Toxin | 99.67% (605/607) | Thymidine kinase |

| | | | | | | | | | | | |
|---|---|---|---|---|---|---|---|---|---|---|---|
| 111 | B*15:01 | 31 | 40 | 10 | AQYKCVTIKY | 0.90655 | 0.02 | -0.20764 | Non-Toxin | 100.00% (607/607) | Thymidine kinase |
| 112 | B*15:01 | 84 | 93 | 10 | GQFFPDIVEF | 0.903174 | 0.02 | 0.38522 | Non-Toxin | 99.67% (605/607) | Thymidine kinase |
| 113 | A*24:02 | 85 | 93 | 9 | QFFPDIVEF | 0.930703 | 0.02 | 0.26706 | Non-Toxin | 99.67% (605/607) | Thymidine kinase |
| 114 | B*44:02 | 143 | 151 | 9 | KEASFSKRL | 0.852596 | 0.03 | -0.34726 | Non-Toxin | 100.00% (607/607) | Thymidine kinase |
| 115 | A*26:01 | 20 | 28 | 9 | ELIRRVRRY | 0.827186 | 0.04 | 0.25846 | Non-Toxin | 99.84% (606/607) | Thymidine kinase |
| 116 | B*44:03 | 143 | 151 | 9 | KEASFSKRL | 0.8979 | 0.04 | -0.34726 | Non-Toxin | 100.00% (607/607) | Thymidine kinase |
| 117 | B*07:02 | 116 | 124 | 9 | RPFNNILNL | 0.897797 | 0.04 | 0.13733 | Non-Toxin | 99.67% (605/607) | Thymidine kinase |
| 118 | A*03:01 | 24 | 34 | 11 | RVRRYQIAQYK | 0.88713 | 0.04 | 0.01899 | Non-Toxin | 100.00% (607/607) | Thymidine kinase |
| 119 | A*30:01 | 24 | 34 | 11 | RVRRYQIAQYK | 0.703275 | 0.04 | 0.01899 | Non-Toxin | 100.00% (607/607) | Thymidine kinase |
| 120 | B*51:01 | 71 | 79 | 9 | EAITDFSVI | 0.758154 | 0.05 | 0.09856 | Non-Toxin | 99.67% (605/607) | Thymidine kinase |
| 121 | B*07:02 | 11 | 21 | 11 | GPMLSGKSTEL | 0.869426 | 0.05 | -0.47073 | Non-Toxin | 0.00% (0/607) | Thymidine kinase |
| 122 | A*02:03 | 13 | 21 | 9 | MLSGKSTEL | 0.81437 | 0.05 | -0.29407 | Non-Toxin | 0.00% (0/607) | Thymidine kinase |
| 123 | A*31:01 | 107 | 115 | 9 | AALDGTFQR | 0.822754 | 0.06 | 0.11938 | Non-Toxin | 100.00% (607/607) | Thymidine kinase |
| 124 | A*26:01 | 68 | 76 | 9 | DVLEAITDF | 0.666283 | 0.06 | 0.30625 | Non-Toxin | 99.67% (605/607) | Thymidine kinase |
| 125 | B*40:01 | 143 | 151 | 9 | KEASFSKRL | 0.91881 | 0.06 | -0.34726 | Non-Toxin | 100.00% (607/607) | Thymidine kinase |
| 126 | A*30:02 | 31 | 40 | 10 | AQYKCVTIKY | 0.663558 | 0.07 | -0.20764 | Non-Toxin | 100.00% (607/607) | Thymidine kinase |
| 127 | B*08:01 | 129 | 137 | 9 | EMVVKLTAV | 0.689913 | 0.07 | -0.10988 | Non-Toxin | 100.00% (607/607) | Thymidine kinase |
| 128 | B*51:01 | 125 | 132 | 8 | IPLSEMVV | 0.720798 | 0.07 | -0.20303 | Non-Toxin | 100.00% (607/607) | Thymidine kinase |
| 129 | B*08:01 | 13 | 21 | 9 | MLSGKSTEL | 0.693376 | 0.07 | -0.29407 | Non-Toxin | 0.00% (0/607) | Thymidine kinase |
| 130 | A*11:01 | 107 | 115 | 9 | AALDGTFQR | 0.793233 | 0.08 | 0.11938 | Non-Toxin | 100.00% (607/607) | Thymidine kinase |
| 131 | A*30:01 | 140 | 149 | 10 | KCFKEASFSK | 0.601969 | 0.08 | -0.19699 | Non-Toxin | 100.00% (607/607) | Thymidine kinase |
| 132 | B*51:01 | 97 | 105 | 9 | MANEGKIVI | 0.655983 | 0.09 | 0.06509 | Non-Toxin | 100.00% (607/607) | Thymidine kinase |
| 133 | A*30:01 | 145 | 153 | 9 | ASFSKRLGA | 0.546245 | 0.1 | -0.27931 | Non-Toxin | 3.95% (24/607) | Thymidine kinase |
| 134 | A*02:01 | 65 | 73 | 9 | KLCDVLEAI | 0.750096 | 0.1 | 0.14194 | Non-Toxin | 99.84% (606/607) | Thymidine kinase |
| 135 | A*02:06 | 123 | 131 | 9 | NLIPLSEMV | 0.69242 | 0.11 | -0.15259 | Non-Toxin | 100.00% (607/607) | Thymidine kinase |
| 136 | A*02:01 | 123 | 131 | 9 | NLIPLSEMV | 0.717546 | 0.12 | -0.15259 | Non-Toxin | 100.00% (607/607) | Thymidine kinase |
| 137 | A*03:01 | 31 | 39 | 9 | AQYKCVTIK | 0.756942 | 0.13 | -0.12132 | Non-Toxin | 100.00% (607/607) | Thymidine kinase |
| 138 | A*02:06 | 65 | 73 | 9 | KLCDVLEAI | 0.667753 | 0.13 | 0.14194 | Non-Toxin | 99.84% (606/607) | Thymidine kinase |
| 139 | A*03:01 | 9 | 17 | 9 | IIGPMLSGK | 0.728889 | 0.14 | -0.30142 | Non-Toxin | 0.00% (0/607) | Thymidine kinase |
| 140 | A*03:01 | 8 | 17 | 10 | LIIGPMLSGK | 0.725536 | 0.14 | -0.23476 | Non-Toxin | 0.00% (0/607) | Thymidine kinase |
| 141 | A*02:01 | 13 | 21 | 9 | MLSGKSTEL | 0.666953 | 0.15 | -0.29407 | Non-Toxin | 0.00% (0/607) | Thymidine kinase |
| 142 | A*02:03 | 123 | 131 | 9 | NLIPLSEMV | 0.601274 | 0.15 | -0.15259 | Non-Toxin | 100.00% (607/607) | Thymidine kinase |
| 143 | A*23:01 | 84 | 93 | 10 | GQFFPDIVEF | 0.505602 | 0.16 | 0.38522 | Non-Toxin | 99.67% (605/607) | Thymidine kinase |
| 144 | A*11:01 | 24 | 34 | 11 | RVRRYQIAQYK | 0.680152 | 0.16 | 0.01899 | Non-Toxin | 100.00% (607/607) | Thymidine kinase |
| 145 | A*68:01 | 107 | 115 | 9 | AALDGTFQR | 0.824072 | 0.19 | 0.11938 | Non-Toxin | 100.00% (607/607) | Thymidine kinase |

| # | allele | start | end | length | peptide | score | percentile_rank | Immunogenicity | Toxicity | Conservancy | Protein |
|---|---|---|---|---|---|---|---|---|---|---|---|
| 146 | A*31:01 | 36 | 44 | 9 | GTYSLIIHR | 0.9613 | 0.01 | 0.07803 | Non-Toxin | 100.00% (86/86) | Cu-Zn superoxide dismutase-like protein |
| 147 | A*11:01 | 36 | 44 | 9 | GTYSLIIHR | 0.958563 | 0.01 | 0.07803 | Non-Toxin | 100.00% (86/86) | Cu-Zn superoxide dismutase-like protein |
| 148 | A*24:02 | 48 | 57 | 10 | SYINEKIIHF | 0.972814 | 0.01 | 0.18069 | Non-Toxin | 100.00% (86/86) | Cu-Zn superoxide dismutase-like protein |
| 149 | A*23:01 | 48 | 57 | 10 | SYINEKIIHF | 0.968015 | 0.01 | 0.18069 | Non-Toxin | 100.00% (86/86) | Cu-Zn superoxide dismutase-like protein |
| 150 | A*03:01 | 25 | 34 | 10 | KVLGSVIGLK | 0.945271 | 0.02 | 0.057 | Non-Toxin | 100.00% (86/86) | Cu-Zn superoxide dismutase-like protein |
| 151 | A*30:02 | 37 | 45 | 9 | TYSLIIHRY | 0.785996 | 0.03 | 0.24756 | Non-Toxin | 100.00% (86/86) | Cu-Zn superoxide dismutase-like protein |
| 152 | B*15:01 | 49 | 57 | 9 | YINEKIIHF | 0.87105 | 0.03 | 0.14515 | Non-Toxin | 100.00% (86/86) | Cu-Zn superoxide dismutase-like protein |
| 153 | B*08:01 | 49 | 57 | 9 | YINEKIIHF | 0.816421 | 0.03 | 0.14515 | Non-Toxin | 100.00% (86/86) | Cu-Zn superoxide dismutase-like protein |
| 154 | A*68:01 | 36 | 44 | 9 | GTYSLIIHR | 0.940165 | 0.04 | 0.07803 | Non-Toxin | 100.00% (86/86) | Cu-Zn superoxide dismutase-like protein |
| 155 | A*31:01 | 34 | 44 | 11 | KSGTYSLIIHR | 0.85456 | 0.04 | 0.13106 | Non-Toxin | 100.00% (86/86) | Cu-Zn superoxide dismutase-like protein |
| 156 | A*11:01 | 25 | 34 | 10 | KVLGSVIGLK | 0.87655 | 0.04 | 0.057 | Non-Toxin | 100.00% (86/86) | Cu-Zn superoxide dismutase-like protein |
| 157 | B*35:01 | 6 | 14 | 9 | FVNRYGVAY | 0.878096 | 0.05 | 0.13598 | Non-Toxin | 100.00% (86/86) | Cu-Zn superoxide dismutase-like protein |
| 158 | B*15:01 | 6 | 14 | 9 | FVNRYGVAY | 0.824564 | 0.05 | 0.13598 | Non-Toxin | 100.00% (86/86) | Cu-Zn superoxide dismutase-like protein |
| 159 | A*02:03 | 32 | 40 | 9 | GLKSGTYSL | 0.823261 | 0.05 | -0.26671 | Non-Toxin | 100.00% (86/86) | Cu-Zn superoxide dismutase-like protein |
| 160 | A*03:01 | 36 | 44 | 9 | GTYSLIIHR | 0.852567 | 0.05 | 0.07803 | Non-Toxin | 100.00% (86/86) | Cu-Zn superoxide dismutase-like protein |
| 161 | A*30:02 | 36 | 45 | 10 | GTYSLIIHRY | 0.717042 | 0.05 | 0.13395 | Non-Toxin | 100.00% (86/86) | Cu-Zn superoxide dismutase-like protein |
| 162 | A*32:01 | 25 | 33 | 9 | KVLGSVIGL | 0.661664 | 0.05 | 0.04038 | Non-Toxin | 100.00% (86/86) | Cu-Zn superoxide dismutase-like protein |

| | | | | | | | | | | | |
|---|---|---|---|---|---|---|---|---|---|---|---|
| 163 | A*26:01 | 29 | 38 | 10 | SVIGLKSGTY | 0.772906 | 0.05 | -0.24795 | Non-Toxin | 100.00% (86/86) | Cu-Zn superoxide dismutase-like protein |
| 164 | A*32:01 | 49 | 57 | 9 | YINEKIIHF | 0.639719 | 0.05 | 0.14515 | Non-Toxin | 100.00% (86/86) | Cu-Zn superoxide dismutase-like protein |
| 165 | A*02:06 | 25 | 33 | 9 | KVLGSVIGL | 0.851332 | 0.06 | 0.04038 | Non-Toxin | 100.00% (86/86) | Cu-Zn superoxide dismutase-like protein |
| 166 | A*24:02 | 9 | 17 | 9 | RYGVAYVYL | 0.798815 | 0.06 | 0.11984 | Non-Toxin | 100.00% (86/86) | Cu-Zn superoxide dismutase-like protein |
| 167 | A*31:01 | 35 | 44 | 10 | SGTYSLIIHR | 0.817554 | 0.06 | 0.09348 | Non-Toxin | 100.00% (86/86) | Cu-Zn superoxide dismutase-like protein |
| 168 | A*24:02 | 15 | 23 | 9 | VYLDTDVNI | 0.771787 | 0.06 | 0.10846 | Non-Toxin | 100.00% (86/86) | Cu-Zn superoxide dismutase-like protein |
| 169 | A*26:01 | 49 | 57 | 9 | YINEKIIHF | 0.733693 | 0.06 | 0.14515 | Non-Toxin | 100.00% (86/86) | Cu-Zn superoxide dismutase-like protein |
| 170 | A*26:01 | 6 | 14 | 9 | FVNRYGVAY | 0.639742 | 0.07 | 0.13598 | Non-Toxin | 100.00% (86/86) | Cu-Zn superoxide dismutase-like protein |
| 171 | A*68:01 | 35 | 44 | 10 | SGTYSLIIHR | 0.907589 | 0.07 | 0.09348 | Non-Toxin | 100.00% (86/86) | Cu-Zn superoxide dismutase-like protein |
| 172 | A*23:01 | 48 | 58 | 11 | SYINEKIIHFL | 0.703303 | 0.08 | 0.27477 | Non-Toxin | 100.00% (86/86) | Cu-Zn superoxide dismutase-like protein |
| 173 | A*23:01 | 15 | 23 | 9 | VYLDTDVNI | 0.69722 | 0.08 | 0.10846 | Non-Toxin | 100.00% (86/86) | Cu-Zn superoxide dismutase-like protein |
| 174 | B*15:01 | 29 | 38 | 10 | SVIGLKSGTY | 0.761117 | 0.09 | -0.24795 | Non-Toxin | 100.00% (86/86) | Cu-Zn superoxide dismutase-like protein |
| 175 | A*24:02 | 48 | 58 | 11 | SYINEKIIHFL | 0.717031 | 0.09 | 0.27477 | Non-Toxin | 100.00% (86/86) | Cu-Zn superoxide dismutase-like protein |
| 176 | A*23:01 | 37 | 45 | 9 | TYSLIIHRY | 0.656855 | 0.09 | 0.24756 | Non-Toxin | 100.00% (86/86) | Cu-Zn superoxide dismutase-like protein |
| 177 | A*23:01 | 9 | 17 | 9 | RYGVAYVYL | 0.635589 | 0.1 | 0.11984 | Non-Toxin | 100.00% (86/86) | Cu-Zn superoxide dismutase-like protein |
| 178 | A*30:02 | 29 | 38 | 10 | SVIGLKSGTY | 0.5916 | 0.11 | -0.24795 | Non-Toxin | 100.00% (86/86) | Cu-Zn superoxide dismutase-like protein |
| 179 | A*24:02 | 37 | 45 | 9 | TYSLIIHRY | 0.694382 | 0.11 | 0.24756 | Non-Toxin | 100.00% (86/86) | Cu-Zn superoxide dismutase-like protein |
| 180 | A*03:01 | 26 | 34 | 9 | VLGSVIGLK | 0.770565 | 0.11 | 0.03213 | Non-Toxin | 100.00% (86/86) | Cu-Zn superoxide dismutase-like protein |
| **#** | **allele** | **start** | **end** | **length** | **peptide** | **score** | **percentile_rank** | **Immunogenicity** | **Toxicity** | **Conservancy** | **Protein** |
| 181 | A*68:01 | 4 | 12 | 9 | DTISDVKQK | 0.958 | 0.03 | -0.31249 | Non-Toxin | 100.00% (574/574) | Envelope protein A28 |
| 182 | B*15:01 | 10 | 18 | 9 | KQKWRCVVY | 0.862449 | 0.03 | 0.2115 | Non-Toxin | 100.00% (574/574) | Envelope protein A28 |
| 183 | A*31:01 | 42 | 50 | 9 | SIRKFNTMR | 0.898278 | 0.03 | -0.16213 | Non-Toxin | 100.00% (574/574) | Envelope protein A28 |
| 184 | A*30:02 | 22 | 30 | 9 | QSYSIYENY | 0.751595 | 0.04 | 0.03917 | Non-Toxin | 100.00% (574/574) | Envelope protein A28 |
| 185 | A*02:06 | 27 | 35 | 9 | SIFGFQAEV | 0.888479 | 0.04 | 0.16858 | Non-Toxin | 100.00% (574/574) | Envelope protein A28 |
| 186 | A*02:01 | 27 | 35 | 9 | SIFGFQAEV | 0.876949 | 0.04 | 0.16858 | Non-Toxin | 100.00% (574/574) | Envelope protein A28 |
| 187 | A*02:03 | 27 | 35 | 9 | SIFGFQAEV | 0.872211 | 0.04 | 0.16858 | Non-Toxin | 100.00% (574/574) | Envelope protein A28 |
| 188 | B*58:01 | 5 | 13 | 9 | TISDVKQKW | 0.931204 | 0.04 | -0.41794 | Non-Toxin | 100.00% (574/574) | Envelope protein A28 |
| 189 | A*23:01 | 26 | 36 | 11 | IYENYGNIKEF | 0.788913 | 0.05 | 0.05087 | Non-Toxin | 100.00% (574/574) | Envelope protein A28 |
| 190 | B*35:01 | 37 | 45 | 9 | NATHAAFEY | 0.890759 | 0.05 | 0.27738 | Non-Toxin | 100.00% (574/574) | Envelope protein A28 |
| 191 | A*68:02 | 27 | 35 | 9 | SIFGFQAEV | 0.820302 | 0.05 | 0.16858 | Non-Toxin | 100.00% (574/574) | Envelope protein A28 |
| 192 | A*31:01 | 41 | 50 | 10 | RSIRKFNTMR | 0.80963 | 0.06 | -0.07665 | Non-Toxin | 100.00% (574/574) | Envelope protein A28 |
| 193 | B*53:01 | 5 | 13 | 9 | TISDVKQKW | 0.68864 | 0.06 | -0.41794 | Non-Toxin | 100.00% (574/574) | Envelope protein A28 |
| 194 | B*58:01 | 6 | 13 | 8 | ISDVKQKW | 0.888271 | 0.07 | -0.4523 | Non-Toxin | 100.00% (574/574) | Envelope protein A28 |
| 195 | A*24:02 | 26 | 36 | 11 | IYENYGNIKEF | 0.770246 | 0.07 | 0.05087 | Non-Toxin | 100.00% (574/574) | Envelope protein A28 |
| 196 | B*57:01 | 5 | 13 | 9 | TISDVKQKW | 0.932363 | 0.08 | -0.41794 | Non-Toxin | 100.00% (574/574) | Envelope protein A28 |
| 197 | B*44:03 | 27 | 36 | 10 | YENYGNIKEF | 0.79744 | 0.08 | 0.02266 | Non-Toxin | 100.00% (574/574) | Envelope protein A28 |
| 198 | B*44:02 | 27 | 36 | 10 | YENYGNIKEF | 0.732977 | 0.08 | 0.02266 | Non-Toxin | 100.00% (574/574) | Envelope protein A28 |
| 199 | B*51:01 | 41 | 49 | 9 | AAFEYSKSI | 0.661134 | 0.09 | -0.29924 | Non-Toxin | 99.83% (573/574) | Envelope protein A28 |
| 200 | B*44:03 | 34 | 43 | 10 | KEFNATHAAF | 0.779999 | 0.09 | 0.19372 | Non-Toxin | 100.00% (574/574) | Envelope protein A28 |
| 201 | A*32:01 | 5 | 13 | 9 | TISDVKQKW | 0.534263 | 0.09 | -0.41794 | Non-Toxin | 100.00% (574/574) | Envelope protein A28 |
| 202 | A*11:01 | 38 | 47 | 10 | ATHAAFEYSK | 0.764078 | 0.1 | 0.19644 | Non-Toxin | 99.83% (573/574) | Envelope protein A28 |
| 203 | B*08:01 | 42 | 49 | 8 | SIRKFNTM | 0.578945 | 0.11 | -0.05953 | Non-Toxin | 100.00% (574/574) | Envelope protein A28 |
| 204 | B*44:02 | 34 | 43 | 10 | KEFNATHAAF | 0.667246 | 0.12 | 0.19372 | Non-Toxin | 100.00% (574/574) | Envelope protein A28 |
| 205 | A*26:01 | 22 | 30 | 9 | QSYSIYENY | 0.473192 | 0.13 | 0.03917 | Non-Toxin | 100.00% (574/574) | Envelope protein A28 |
| 206 | A*68:02 | 1 | 9 | 9 | DVNDTISDV | 0.57861 | 0.14 | 0.05664 | Non-Toxin | 100.00% (574/574) | Envelope protein A28 |
| 207 | B*57:01 | 6 | 13 | 8 | ISDVKQKW | 0.875809 | 0.14 | -0.4523 | Non-Toxin | 100.00% (574/574) | Envelope protein A28 |
| 208 | B*40:01 | 34 | 43 | 10 | KEFNATHAAF | 0.696897 | 0.15 | 0.19372 | Non-Toxin | 100.00% (574/574) | Envelope protein A28 |
| 209 | A*68:01 | 32 | 41 | 10 | QAEVGPNNTR | 0.849929 | 0.15 | 0.10737 | Non-Toxin | 100.00% (574/574) | Envelope protein A28 |
| 210 | B*40:01 | 33 | 43 | 11 | AEVGPNNTRSI | 0.664546 | 0.17 | 0.00766 | Non-Toxin | 100.00% (574/574) | Envelope protein A28 |
| 211 | A*30:02 | 10 | 18 | 9 | KQKWRCVVY | 0.486796 | 0.18 | 0.2115 | Non-Toxin | 100.00% (574/574) | Envelope protein A28 |
| 212 | A*68:01 | 4 | 14 | 11 | DTISDVKQKWR | 0.825768 | 0.19 | -0.43309 | Non-Toxin | 100.00% (574/574) | Envelope protein A28 |
| 213 | B*08:01 | 8 | 16 | 9 | DVKQKWRCV | 0.455079 | 0.19 | -0.17587 | Non-Toxin | 100.00% (574/574) | Envelope protein A28 |
| 214 | A*33:01 | 42 | 50 | 9 | SIRKFNTMR | 0.471976 | 0.19 | -0.16213 | Non-Toxin | 100.00% (574/574) | Envelope protein A28 |

| | | | | | | | | | | | |
|---|---|---|---|---|---|---|---|---|---|---|---|
| 215 | A*68:01 | 5 | 14 | 10 | TISDVKQKWR | 0.819013 | 0.19 | -0.3628 | Non-Toxin | 100.00% (574/574) | Envelope protein A28 |
| 216 | A*68:02 | 16 | 24 | 9 | VVYPGNGFV | 0.510667 | 0.19 | 0.11155 | Non-Toxin | 100.00% (574/574) | Envelope protein A28 |
| **#** | **allele** | **start** | **end** | **length** | **peptide** | **score** | **percentile_rank** | **Immunogenicity** | **Toxicity** | **Conservancy** | **Protein** |
| 217 | A*11:01 | 36 | 44 | 9 | ATFTVNIFK | 0.982136 | 0.01 | 0.29189 | Non-Toxin | 100.00% (10/10) | DNA-directed RNA polymerase |
| 218 | B*51:01 | 33 | 40 | 8 | DALATFTV | 0.924732 | 0.01 | 0.21653 | Non-Toxin | 100.00% (10/10) | DNA-directed RNA polymerase |
| 219 | A*33:01 | 8 | 16 | 9 | DMYKSRVER | 0.942288 | 0.01 | -0.23724 | Non-Toxin | 100.00% (10/10) | DNA-directed RNA polymerase |
| 220 | B*35:01 | 3 | 11 | 9 | FPAEFRDGY | 0.991885 | 0.01 | 0.31469 | Non-Toxin | 100.00% (10/10) | DNA-directed RNA polymerase |
| 221 | B*35:01 | 48 | 56 | 9 | FPITIENAL | 0.977844 | 0.01 | 0.32351 | Non-Toxin | 100.00% (10/10) | DNA-directed RNA polymerase |
| 222 | A*30:02 | 4 | 12 | 9 | GSYRTHPHY | 0.845882 | 0.01 | 0.12867 | Non-Toxin | 100.00% (10/10) | DNA-directed RNA polymerase |
| 223 | A*01:01 | 38 | 46 | 9 | ITDFDIDTY | 0.988568 | 0.01 | 0.31328 | Non-Toxin | 60.00% (6/10) | DNA-directed RNA polymerase |
| 224 | B*40:01 | 21 | 29 | 9 | KEFPHVIEM | 0.992796 | 0.01 | 0.26802 | Non-Toxin | 100.00% (10/10) | DNA-directed RNA polymerase |
| 225 | A*03:01 | 28 | 37 | 10 | KLNLSPLLTK | 0.983572 | 0.01 | -0.18228 | Non-Toxin | 100.00% (10/10) | DNA-directed RNA polymerase |
| 226 | B*58:01 | 30 | 38 | 9 | KSYDALATF | 0.978211 | 0.01 | 0.10448 | Non-Toxin | 100.00% (10/10) | DNA-directed RNA polymerase |
| 227 | A*32:01 | 30 | 38 | 9 | KSYDALATF | 0.928526 | 0.01 | 0.10448 | Non-Toxin | 100.00% (10/10) | DNA-directed RNA polymerase |
| 228 | A*03:01 | 47 | 55 | 9 | KVFFGPIYY | 0.955581 | 0.01 | 0.28852 | Non-Toxin | 100.00% (10/10) | DNA-directed RNA polymerase |
| 229 | A*30:02 | 47 | 55 | 9 | KVFFGPIYY | 0.951959 | 0.01 | 0.28852 | Non-Toxin | 100.00% (10/10) | DNA-directed RNA polymerase |
| 230 | A*11:01 | 47 | 55 | 9 | KVFFGPIYY | 0.933312 | 0.01 | 0.28852 | Non-Toxin | 100.00% (10/10) | DNA-directed RNA polymerase |
| 231 | A*32:01 | 47 | 55 | 9 | KVFFGPIYY | 0.866851 | 0.01 | 0.28852 | Non-Toxin | 100.00% (10/10) | DNA-directed RNA polymerase |
| 232 | A*30:02 | 41 | 49 | 9 | KYHPSQYLY | 0.96213 | 0.01 | -0.2804 | Non-Toxin | 100.00% (10/10) | DNA-directed RNA polymerase |
| 233 | A*24:02 | 41 | 50 | 10 | KYHPSQYLYF | 0.984281 | 0.01 | -0.28956 | Non-Toxin | 100.00% (10/10) | DNA-directed RNA polymerase |
| 234 | A*23:01 | 41 | 50 | 10 | KYHPSQYLYF | 0.974592 | 0.01 | -0.28956 | Non-Toxin | 100.00% (10/10) | DNA-directed RNA polymerase |
| 235 | B*51:01 | 16 | 24 | 9 | MPPEVVYLV | 0.948739 | 0.01 | 0.16661 | Non-Toxin | 100.00% (10/10) | DNA-directed RNA polymerase |
| 236 | A*30:02 | 40 | 49 | 10 | RKYHPSQYLY | 0.841155 | 0.01 | -0.25919 | Non-Toxin | 100.00% (10/10) | DNA-directed RNA polymerase |
| 237 | A*03:01 | 45 | 53 | 9 | RLHEILTVK | 0.96551 | 0.01 | 0.28729 | Non-Toxin | 100.00% (10/10) | DNA-directed RNA polymerase |
| 238 | B*44:03 | 7 | 15 | 9 | SEMDQRLGY | 0.993779 | 0.01 | -0.08832 | Non-Toxin | 100.00% (10/10) | DNA-directed RNA polymerase |
| 239 | B*44:02 | 7 | 15 | 9 | SEMDQRLGY | 0.986859 | 0.01 | -0.08832 | Non-Toxin | 100.00% (10/10) | DNA-directed RNA polymerase |
| 240 | B*44:03 | 7 | 17 | 11 | SEMDQRLGYKF | 0.971043 | 0.01 | -0.2051 | Non-Toxin | 100.00% (10/10) | DNA-directed RNA polymerase |
| 241 | B*44:02 | 7 | 17 | 11 | SEMDQRLGYKF | 0.963669 | 0.01 | -0.2051 | Non-Toxin | 100.00% (10/10) | DNA-directed RNA polymerase |
| 242 | B*15:01 | 48 | 55 | 8 | SQKGTVAY | 0.948209 | 0.01 | 0.07378 | Non-Toxin | 100.00% (10/10) | DNA-directed RNA polymerase |
| 243 | A*23:01 | 35 | 44 | 10 | SYYKDDISYF | 0.929358 | 0.01 | -0.1915 | Non-Toxin | 100.00% (10/10) | DNA-directed RNA polymerase |
| 244 | B*08:01 | 31 | 39 | 9 | TMKERRPIL | 0.945208 | 0.01 | 0.19827 | Non-Toxin | 100.00% (10/10) | DNA-directed RNA polymerase |
| 245 | A*24:02 | 1 | 9 | 9 | TYGKYFETL | 0.96099 | 0.01 | 0.00778 | Non-Toxin | 100.00% (10/10) | DNA-directed RNA polymerase |
| 246 | A*01:01 | 2 | 10 | 9 | VSEKYTDMY | 0.980424 | 0.01 | -0.23544 | Non-Toxin | 100.00% (10/10) | DNA-directed RNA polymerase |
| 247 | A*01:01 | 21 | 30 | 10 | YTPDQLKGFY | 0.968586 | 0.01 | -0.21088 | Non-Toxin | 100.00% (10/10) | DNA-directed RNA polymerase |
| 248 | A*24:02 | 36 | 44 | 9 | YYKDDISYF | 0.976663 | 0.01 | -0.04258 | Non-Toxin | 100.00% (10/10) | DNA-directed RNA polymerase |
| 249 | A*23:01 | 36 | 44 | 9 | YYKDDISYF | 0.970683 | 0.01 | -0.04258 | Non-Toxin | 100.00% (10/10) | DNA-directed RNA polymerase |
| 250 | A*03:01 | 36 | 44 | 9 | ATFTVNIFK | 0.945804 | 0.02 | 0.29189 | Non-Toxin | 100.00% (10/10) | DNA-directed RNA polymerase |
| 251 | A*68:01 | 38 | 46 | 9 | EASNISSTK | 0.965415 | 0.02 | -0.20328 | Non-Toxin | 100.00% (10/10) | DNA-directed RNA polymerase |
| 252 | B*35:01 | 52 | 60 | 9 | FARDNQISF | 0.946137 | 0.02 | -0.06056 | Non-Toxin | 100.00% (10/10) | DNA-directed RNA polymerase |
| 253 | A*33:01 | 49 | 57 | 9 | FFGPIYYLR | 0.849406 | 0.02 | 0.11636 | Non-Toxin | 100.00% (10/10) | DNA-directed RNA polymerase |
| 254 | B*51:01 | 23 | 30 | 8 | FPHVIEMV | 0.915641 | 0.02 | 0.12769 | Non-Toxin | 100.00% (10/10) | DNA-directed RNA polymerase |
| 255 | A*32:01 | 25 | 33 | 9 | GVFYRPLHF | 0.772951 | 0.02 | 0.08378 | Non-Toxin | 100.00% (10/10) | DNA-directed RNA polymerase |
| 256 | A*31:01 | 39 | 47 | 9 | IIKKQFIQR | 0.923194 | 0.02 | -0.24496 | Non-Toxin | 100.00% (10/10) | DNA-directed RNA polymerase |
| 257 | A*30:02 | 52 | 60 | 9 | ISITKISSY | 0.802309 | 0.02 | -0.23874 | Non-Toxin | 100.00% (10/10) | DNA-directed RNA polymerase |
| 258 | A*30:02 | 14 | 22 | 9 | ITNILTSEY | 0.823801 | 0.02 | 0.07644 | Non-Toxin | 100.00% (10/10) | DNA-directed RNA polymerase |
| 259 | B*57:01 | 30 | 38 | 9 | KSYDALATF | 0.985443 | 0.02 | 0.10448 | Non-Toxin | 100.00% (10/10) | DNA-directed RNA polymerase |
| 260 | A*02:06 | 24 | 32 | 9 | KTISMLIEV | 0.936668 | 0.02 | -0.13389 | Non-Toxin | 100.00% (10/10) | DNA-directed RNA polymerase |
| 261 | A*31:01 | 47 | 57 | 11 | KVFFGPIYYLR | 0.910813 | 0.02 | 0.29452 | Non-Toxin | 100.00% (10/10) | DNA-directed RNA polymerase |
| 262 | B*08:01 | 24 | 32 | 9 | KVKVRVLTM | 0.870399 | 0.02 | 0.07412 | Non-Toxin | 100.00% (10/10) | DNA-directed RNA polymerase |
| 263 | B*51:01 | 16 | 23 | 8 | MPPEVVYL | 0.852183 | 0.02 | 0.17309 | Non-Toxin | 100.00% (10/10) | DNA-directed RNA polymerase |
| 264 | B*44:02 | 34 | 44 | 11 | QEDGIIIKKQF | 0.912305 | 0.02 | -0.02258 | Non-Toxin | 100.00% (10/10) | DNA-directed RNA polymerase |
| 265 | A*30:01 | 45 | 53 | 9 | RLHEILTVK | 0.81383 | 0.02 | 0.28729 | Non-Toxin | 100.00% (10/10) | DNA-directed RNA polymerase |
| 266 | B*57:01 | 7 | 14 | 8 | RTHPHYSW | 0.98452 | 0.02 | -0.11226 | Non-Toxin | 100.00% (10/10) | DNA-directed RNA polymerase |

| | | | | | | | | | | | |
|---|---|---|---|---|---|---|---|---|---|---|---|
| 267 | A*32:01 | 1 | 10 | 10 | RVVKPNSFTF | 0.781209 | 0.02 | -0.25495 | Non-Toxin | 100.00% (10/10) | DNA-directed RNA polymerase |
| 268 | A*01:01 | 37 | 46 | 10 | SITDFDIDTY | 0.936788 | 0.02 | 0.3372 | Non-Toxin | 60.00% (6/10) | DNA-directed RNA polymerase |
| 269 | A*33:01 | 37 | 45 | 9 | SYSNFILHR | 0.819403 | 0.02 | 0.18861 | Non-Toxin | 100.00% (10/10) | DNA-directed RNA polymerase |
| 270 | A*24:02 | 35 | 44 | 10 | SYYKDDISYF | 0.92973 | 0.02 | -0.1915 | Non-Toxin | 100.00% (10/10) | DNA-directed RNA polymerase |
| 271 | A*24:02 | 34 | 42 | 9 | TYAVNIHVL | 0.929761 | 0.02 | 0.22464 | Non-Toxin | 100.00% (10/10) | DNA-directed RNA polymerase |
| 272 | A*23:01 | 1 | 9 | 9 | TYGKYFETL | 0.896453 | 0.02 | 0.00778 | Non-Toxin | 100.00% (10/10) | DNA-directed RNA polymerase |
| 273 | A*11:01 | 22 | 30 | 9 | VTIALMSYK | 0.907142 | 0.02 | -0.23531 | Non-Toxin | 100.00% (10/10) | DNA-directed RNA polymerase |
| 274 | B*15:01 | 2 | 10 | 9 | VVKPNSFTF | 0.913946 | 0.02 | -0.12171 | Non-Toxin | 100.00% (10/10) | DNA-directed RNA polymerase |
| 275 | A*32:01 | 2 | 10 | 9 | VVKPNSFTF | 0.809306 | 0.02 | -0.12171 | Non-Toxin | 100.00% (10/10) | DNA-directed RNA polymerase |
| 276 | A*68:02 | 38 | 46 | 9 | FTVNIFKEV | 0.880902 | 0.03 | 0.12319 | Non-Toxin | 100.00% (10/10) | DNA-directed RNA polymerase |
| 277 | A*30:02 | 27 | 35 | 9 | FYRPLHFQY | 0.763641 | 0.03 | 0.05641 | Non-Toxin | 100.00% (10/10) | DNA-directed RNA polymerase |
| 278 | A*01:01 | 14 | 22 | 9 | ITNILTSEY | 0.916606 | 0.03 | 0.07644 | Non-Toxin | 100.00% (10/10) | DNA-directed RNA polymerase |
| 279 | B*44:03 | 21 | 29 | 9 | KEFPHVIEM | 0.937788 | 0.03 | 0.26802 | Non-Toxin | 100.00% (10/10) | DNA-directed RNA polymerase |
| 280 | B*44:02 | 21 | 29 | 9 | KEFPHVIEM | 0.849906 | 0.03 | 0.26802 | Non-Toxin | 100.00% (10/10) | DNA-directed RNA polymerase |
| 281 | A*01:01 | 48 | 57 | 10 | KIDNGIHLMY | 0.918963 | 0.03 | 0.08178 | Non-Toxin | 100.00% (10/10) | DNA-directed RNA polymerase |
| 282 | A*30:02 | 48 | 57 | 10 | KIDNGIHLMY | 0.77521 | 0.03 | 0.08178 | Non-Toxin | 100.00% (10/10) | DNA-directed RNA polymerase |
| 283 | B*53:01 | 4 | 12 | 9 | KPNSFTFSF | 0.810527 | 0.03 | -0.01589 | Non-Toxin | 100.00% (10/10) | DNA-directed RNA polymerase |
| 284 | B*40:01 | 21 | 29 | 9 | LESNGLVRL | 0.950514 | 0.03 | 0.02743 | Non-Toxin | 100.00% (10/10) | DNA-directed RNA polymerase |
| 285 | A*02:01 | 28 | 36 | 9 | MLIEVILTA | 0.911823 | 0.03 | 0.32275 | Non-Toxin | 90.00% (9/10) | DNA-directed RNA polymerase |
| 286 | A*02:03 | 30 | 38 | 9 | NLSPLLTKI | 0.888015 | 0.03 | -0.17934 | Non-Toxin | 100.00% (10/10) | DNA-directed RNA polymerase |
| 287 | B*58:01 | 7 | 14 | 8 | RTHPHYSW | 0.961252 | 0.03 | -0.11226 | Non-Toxin | 100.00% (10/10) | DNA-directed RNA polymerase |
| 288 | A*02:01 | 11 | 20 | 10 | SLSYDMPPEV | 0.90696 | 0.03 | -0.16812 | Non-Toxin | 100.00% (10/10) | DNA-directed RNA polymerase |
| 289 | A*02:03 | 11 | 20 | 10 | SLSYDMPPEV | 0.881998 | 0.03 | -0.16812 | Non-Toxin | 100.00% (10/10) | DNA-directed RNA polymerase |
| 290 | A*31:01 | 37 | 45 | 9 | SYSNFILHR | 0.89425 | 0.03 | 0.18861 | Non-Toxin | 100.00% (10/10) | DNA-directed RNA polymerase |
| 291 | B*51:01 | 16 | 24 | 9 | TPPDYSPII | 0.838912 | 0.03 | -0.07221 | Non-Toxin | 100.00% (10/10) | DNA-directed RNA polymerase |
| 292 | A*02:03 | 34 | 42 | 9 | ALATFTVNI | 0.873717 | 0.04 | 0.23336 | Non-Toxin | 100.00% (10/10) | DNA-directed RNA polymerase |
| 293 | A*24:02 | 19 | 27 | 9 | DYSPIIASI | 0.837371 | 0.04 | 0.12638 | Non-Toxin | 100.00% (10/10) | DNA-directed RNA polymerase |
| 294 | B*53:01 | 48 | 56 | 9 | FPITIENAL | 0.774221 | 0.04 | 0.32351 | Non-Toxin | 100.00% (10/10) | DNA-directed RNA polymerase |
| 295 | B*57:01 | 4 | 14 | 11 | GSYRTHPHYSW | 0.974566 | 0.04 | 0.04005 | Non-Toxin | 100.00% (10/10) | DNA-directed RNA polymerase |
| 296 | B*51:01 | 42 | 50 | 9 | IAHGAANTI | 0.801841 | 0.04 | 0.13675 | Non-Toxin | 100.00% (10/10) | DNA-directed RNA polymerase |
| 297 | B*40:01 | 52 | 60 | 9 | IENALVASL | 0.947398 | 0.04 | 0.00169 | Non-Toxin | 100.00% (10/10) | DNA-directed RNA polymerase |
| 298 | B*15:01 | 52 | 60 | 9 | ISITKISSY | 0.855727 | 0.04 | -0.23874 | Non-Toxin | 100.00% (10/10) | DNA-directed RNA polymerase |
| 299 | A*03:01 | 37 | 45 | 9 | KIDTTHVSK | 0.880032 | 0.04 | 0.05269 | Non-Toxin | 100.00% (10/10) | DNA-directed RNA polymerase |
| 300 | B*35:01 | 4 | 12 | 9 | KPNSFTFSF | 0.916555 | 0.04 | -0.01589 | Non-Toxin | 100.00% (10/10) | DNA-directed RNA polymerase |
| 301 | B*07:02 | 42 | 50 | 9 | KPYESKVFF | 0.917249 | 0.04 | -0.16131 | Non-Toxin | 90.00% (9/10) | DNA-directed RNA polymerase |
| 302 | A*02:03 | 28 | 36 | 9 | MLIEVILTA | 0.854539 | 0.04 | 0.32275 | Non-Toxin | 90.00% (9/10) | DNA-directed RNA polymerase |
| 303 | A*68:01 | 4 | 12 | 9 | NTVSEMDQR | 0.941128 | 0.04 | -0.26983 | Non-Toxin | 60.00% (6/10) | DNA-directed RNA polymerase |
| 304 | B*44:03 | 34 | 44 | 11 | QEDGIIIKKQF | 0.906355 | 0.04 | -0.02258 | Non-Toxin | 100.00% (10/10) | DNA-directed RNA polymerase |
| 305 | A*31:01 | 28 | 36 | 9 | RVLTMKERR | 0.868209 | 0.04 | -0.2238 | Non-Toxin | 100.00% (10/10) | DNA-directed RNA polymerase |
| 306 | A*31:01 | 2 | 10 | 9 | RVSLEFIRR | 0.854859 | 0.04 | 0.2854 | Non-Toxin | 100.00% (10/10) | DNA-directed RNA polymerase |
| 307 | A*68:01 | 14 | 24 | 11 | SVSPPNVLPTR | 0.941775 | 0.04 | -0.03888 | Non-Toxin | 90.00% (9/10) | DNA-directed RNA polymerase |
| 308 | A*23:01 | 34 | 42 | 9 | TYAVNIHVL | 0.840086 | 0.04 | 0.22464 | Non-Toxin | 100.00% (10/10) | DNA-directed RNA polymerase |
| 309 | A*11:01 | 32 | 40 | 9 | VIGGVFINK | 0.861663 | 0.04 | 0.30404 | Non-Toxin | 100.00% (10/10) | DNA-directed RNA polymerase |
| 310 | B*07:02 | 19 | 27 | 9 | VPDPKAGVF | 0.900146 | 0.04 | -0.12441 | Non-Toxin | 100.00% (10/10) | DNA-directed RNA polymerase |
| 311 | B*35:01 | 19 | 27 | 9 | VPDPKAGVF | 0.899213 | 0.04 | -0.12441 | Non-Toxin | 100.00% (10/10) | DNA-directed RNA polymerase |
| 312 | B*53:01 | 19 | 27 | 9 | VPDPKAGVF | 0.791685 | 0.04 | -0.12441 | Non-Toxin | 100.00% (10/10) | DNA-directed RNA polymerase |
| 313 | A*01:01 | 6 | 15 | 10 | VSEMDQRLGY | 0.893825 | 0.04 | -0.17624 | Non-Toxin | 60.00% (6/10) | DNA-directed RNA polymerase |
| 314 | A*68:02 | 11 | 19 | 9 | YVASSLVGI | 0.850254 | 0.04 | -0.27067 | Non-Toxin | 100.00% (10/10) | DNA-directed RNA polymerase |
| 315 | A*02:06 | 41 | 49 | 9 | FILHRLHEI | 0.885851 | 0.05 | 0.15471 | Non-Toxin | 100.00% (10/10) | DNA-directed RNA polymerase |
| 316 | A*02:03 | 17 | 26 | 10 | FLVPDPKAGV | 0.80856 | 0.05 | -0.13714 | Non-Toxin | 100.00% (10/10) | DNA-directed RNA polymerase |
| 317 | B*15:01 | 33 | 41 | 9 | FQYVSYSNF | 0.828876 | 0.05 | -0.26764 | Non-Toxin | 100.00% (10/10) | DNA-directed RNA polymerase |
| 318 | A*03:01 | 33 | 42 | 10 | KILYDPETDK | 0.862622 | 0.05 | 0.14345 | Non-Toxin | 100.00% (10/10) | DNA-directed RNA polymerase |
| 319 | A*31:01 | 32 | 40 | 9 | KISKMFSVR | 0.824663 | 0.05 | -0.447 | Non-Toxin | 100.00% (10/10) | DNA-directed RNA polymerase |

| | | | | | | | | | | | |
|---|---|---|---|---|---|---|---|---|---|---|---|
| 320 | B*07:02 | 4 | 12 | 9 | KPNSFTFSF | 0.878057 | 0.05 | -0.01589 | Non-Toxin | 100.00% (10/10) | DNA-directed RNA polymerase |
| 321 | B*15:01 | 30 | 38 | 9 | KSYDALATF | 0.832661 | 0.05 | 0.10448 | Non-Toxin | 100.00% (10/10) | DNA-directed RNA polymerase |
| 322 | B*15:01 | 47 | 55 | 9 | KVFFGPIYY | 0.821272 | 0.05 | 0.28852 | Non-Toxin | 100.00% (10/10) | DNA-directed RNA polymerase |
| 323 | A*24:02 | 41 | 49 | 9 | KYHPSQYLY | 0.834549 | 0.05 | -0.2804 | Non-Toxin | 100.00% (10/10) | DNA-directed RNA polymerase |
| 324 | A*23:01 | 41 | 49 | 9 | KYHPSQYLY | 0.809752 | 0.05 | -0.2804 | Non-Toxin | 100.00% (10/10) | DNA-directed RNA polymerase |
| 325 | A*02:06 | 28 | 36 | 9 | MLIEVILTA | 0.874505 | 0.05 | 0.32275 | Non-Toxin | 90.00% (9/10) | DNA-directed RNA polymerase |
| 326 | B*44:02 | 21 | 30 | 10 | SEEELSDKIF | 0.7945 | 0.05 | -0.12201 | Non-Toxin | 100.00% (10/10) | DNA-directed RNA polymerase |
| 327 | A*31:01 | 14 | 24 | 11 | SVSPPNVLPTR | 0.837827 | 0.05 | -0.03888 | Non-Toxin | 90.00% (9/10) | DNA-directed RNA polymerase |
| 328 | A*24:02 | 15 | 23 | 9 | TFSDIQKEF | 0.817245 | 0.05 | -0.13432 | Non-Toxin | 100.00% (10/10) | DNA-directed RNA polymerase |
| 329 | A*23:01 | 15 | 23 | 9 | TFSDIQKEF | 0.786468 | 0.05 | -0.13432 | Non-Toxin | 100.00% (10/10) | DNA-directed RNA polymerase |
| 330 | B*15:01 | 51 | 59 | 9 | TVKRPLLSF | 0.840032 | 0.05 | -0.14518 | Non-Toxin | 100.00% (10/10) | DNA-directed RNA polymerase |
| 331 | A*23:01 | 48 | 56 | 9 | VFFGPIYYL | 0.785658 | 0.05 | 0.1813 | Non-Toxin | 100.00% (10/10) | DNA-directed RNA polymerase |
| 332 | A*03:01 | 32 | 40 | 9 | VIGGVFINK | 0.839758 | 0.05 | 0.30404 | Non-Toxin | 100.00% (10/10) | DNA-directed RNA polymerase |
| 333 | A*23:01 | 2 | 10 | 9 | VVKPNSFTF | 0.770061 | 0.05 | -0.12171 | Non-Toxin | 100.00% (10/10) | DNA-directed RNA polymerase |
| 334 | A*02:01 | 34 | 42 | 9 | ALATFTVNI | 0.83955 | 0.06 | 0.23336 | Non-Toxin | 100.00% (10/10) | DNA-directed RNA polymerase |
| 335 | A*01:01 | 36 | 46 | 11 | ESITDFDIDTY | 0.807884 | 0.06 | 0.40614 | Non-Toxin | 60.00% (6/10) | DNA-directed RNA polymerase |
| 336 | A*68:02 | 19 | 27 | 9 | EVVYLVNAI | 0.77186 | 0.06 | 0.05514 | Non-Toxin | 100.00% (10/10) | DNA-directed RNA polymerase |
| 337 | B*40:01 | 6 | 14 | 9 | FETLAHDEL | 0.904424 | 0.06 | 0.14721 | Non-Toxin | 100.00% (10/10) | DNA-directed RNA polymerase |
| 338 | A*02:01 | 41 | 49 | 9 | FILHRLHEI | 0.847747 | 0.06 | 0.15471 | Non-Toxin | 100.00% (10/10) | DNA-directed RNA polymerase |
| 339 | A*02:03 | 41 | 49 | 9 | FILHRLHEI | 0.786816 | 0.06 | 0.15471 | Non-Toxin | 100.00% (10/10) | DNA-directed RNA polymerase |
| 340 | B*44:02 | 10 | 18 | 9 | KERDRSNAY | 0.779628 | 0.06 | -0.04881 | Non-Toxin | 100.00% (10/10) | DNA-directed RNA polymerase |
| 341 | A*11:01 | 37 | 45 | 9 | KIDTTHVSK | 0.832586 | 0.06 | 0.05269 | Non-Toxin | 100.00% (10/10) | DNA-directed RNA polymerase |
| 342 | A*24:02 | 8 | 15 | 8 | KYEMLHNF | 0.778216 | 0.06 | -0.13001 | Non-Toxin | 100.00% (10/10) | DNA-directed RNA polymerase |
| 343 | A*24:02 | 20 | 29 | 10 | NYTPDQLKGF | 0.784791 | 0.06 | -0.2624 | Non-Toxin | 100.00% (10/10) | DNA-directed RNA polymerase |
| 344 | A*24:02 | 34 | 44 | 11 | RSYYKDDISYF | 0.78494 | 0.06 | -0.18822 | Non-Toxin | 100.00% (10/10) | DNA-directed RNA polymerase |
| 345 | B*44:03 | 21 | 30 | 10 | SEEELSDKIF | 0.850975 | 0.06 | -0.12201 | Non-Toxin | 100.00% (10/10) | DNA-directed RNA polymerase |
| 346 | A*02:06 | 6 | 14 | 9 | SQLSVLSSI | 0.844927 | 0.06 | -0.37659 | Non-Toxin | 100.00% (10/10) | DNA-directed RNA polymerase |
| 347 | A*01:01 | 47 | 55 | 9 | TSQKGTVAY | 0.832126 | 0.06 | -0.12736 | Non-Toxin | 100.00% (10/10) | DNA-directed RNA polymerase |
| 348 | A*03:01 | 51 | 60 | 10 | TVKRPLLSFK | 0.831582 | 0.06 | -0.12118 | Non-Toxin | 100.00% (10/10) | DNA-directed RNA polymerase |
| 349 | B*44:03 | 5 | 15 | 11 | TVSEMDQRLGY | 0.843416 | 0.06 | -0.15599 | Non-Toxin | 60.00% (6/10) | DNA-directed RNA polymerase |
| 350 | B*35:01 | 19 | 28 | 10 | VPDPKAGVFY | 0.855051 | 0.06 | -0.04072 | Non-Toxin | 100.00% (10/10) | DNA-directed RNA polymerase |
| 351 | A*24:02 | 2 | 10 | 9 | VVKPNSFTF | 0.780197 | 0.06 | -0.12171 | Non-Toxin | 100.00% (10/10) | DNA-directed RNA polymerase |
| 352 | B*40:01 | 44 | 52 | 9 | FETGFPITI | 0.881194 | 0.07 | 0.28526 | Non-Toxin | 100.00% (10/10) | DNA-directed RNA polymerase |
| 353 | B*35:01 | 9 | 17 | 9 | LAHDELENY | 0.814324 | 0.07 | 0.2006 | Non-Toxin | 100.00% (10/10) | DNA-directed RNA polymerase |
| 354 | A*02:06 | 12 | 20 | 9 | LSYDMPPEV | 0.821978 | 0.07 | -0.11118 | Non-Toxin | 100.00% (10/10) | DNA-directed RNA polymerase |
| 355 | B*07:02 | 5 | 13 | 9 | RPPSFYKPL | 0.841412 | 0.07 | -0.24803 | Non-Toxin | 100.00% (10/10) | DNA-directed RNA polymerase |
| 356 | B*44:02 | 5 | 15 | 11 | TVSEMDQRLGY | 0.761914 | 0.07 | -0.15599 | Non-Toxin | 60.00% (6/10) | DNA-directed RNA polymerase |
| 357 | B*35:01 | 29 | 37 | 9 | VPNVIGGVF | 0.829872 | 0.07 | 0.25366 | Non-Toxin | 100.00% (10/10) | DNA-directed RNA polymerase |
| 358 | A*03:01 | 22 | 30 | 9 | VTIALMSYK | 0.813476 | 0.07 | -0.23531 | Non-Toxin | 100.00% (10/10) | DNA-directed RNA polymerase |
| 359 | A*24:02 | 42 | 50 | 9 | YHPSQYLYF | 0.76082 | 0.07 | -0.29787 | Non-Toxin | 100.00% (10/10) | DNA-directed RNA polymerase |
| 360 | A*02:06 | 11 | 19 | 9 | YVASSLVGI | 0.823538 | 0.07 | -0.27067 | Non-Toxin | 100.00% (10/10) | DNA-directed RNA polymerase |
| 361 | B*15:01 | 19 | 29 | 11 | GQHVTIALMSY | 0.771176 | 0.08 | 0.00224 | Non-Toxin | 100.00% (10/10) | DNA-directed RNA polymerase |
| 362 | B*57:01 | 18 | 26 | 9 | ISIPRSVGF | 0.933533 | 0.08 | -0.01865 | Non-Toxin | 100.00% (10/10) | DNA-directed RNA polymerase |
| 363 | B*58:01 | 18 | 26 | 9 | ISIPRSVGF | 0.864471 | 0.08 | -0.01865 | Non-Toxin | 100.00% (10/10) | DNA-directed RNA polymerase |
| 364 | B*57:01 | 14 | 23 | 10 | MTFSDIQKEF | 0.932541 | 0.08 | -0.20981 | Non-Toxin | 100.00% (10/10) | DNA-directed RNA polymerase |
| 365 | A*02:01 | 30 | 38 | 9 | NLSPLLTKI | 0.802461 | 0.08 | -0.17934 | Non-Toxin | 100.00% (10/10) | DNA-directed RNA polymerase |
| 366 | A*31:01 | 45 | 54 | 10 | RLHEILTVKR | 0.766608 | 0.08 | 0.17543 | Non-Toxin | 100.00% (10/10) | DNA-directed RNA polymerase |
| 367 | B*58:01 | 34 | 44 | 11 | RSYYKDDISYF | 0.871147 | 0.08 | -0.18822 | Non-Toxin | 100.00% (10/10) | DNA-directed RNA polymerase |
| 368 | B*35:01 | 47 | 55 | 9 | TSQKGTVAY | 0.793277 | 0.08 | -0.12736 | Non-Toxin | 100.00% (10/10) | DNA-directed RNA polymerase |
| 369 | A*31:01 | 48 | 57 | 10 | VFFGPIYYLR | 0.760368 | 0.08 | 0.17782 | Non-Toxin | 100.00% (10/10) | DNA-directed RNA polymerase |
| 370 | B*15:01 | 13 | 21 | 9 | VIINSTSIF | 0.773279 | 0.08 | -0.14973 | Non-Toxin | 100.00% (10/10) | DNA-directed RNA polymerase |
| 371 | A*02:06 | 17 | 25 | 9 | VQVELTDKV | 0.781212 | 0.08 | 0.03261 | Non-Toxin | 100.00% (10/10) | DNA-directed RNA polymerase |
| 372 | B*44:03 | 6 | 15 | 10 | VSEMDQRLGY | 0.789083 | 0.08 | -0.17624 | Non-Toxin | 60.00% (6/10) | DNA-directed RNA polymerase |

| | | | | | | | | | | | |
|---|---|---|---|---|---|---|---|---|---|---|---|
| 373 | B*58:01 | 4 | 14 | 11 | GSYRTHPHYSW | 0.852699 | 0.09 | 0.04005 | Non-Toxin | 100.00% (10/10) | DNA-directed RNA polymerase |
| 374 | B*57:01 | 28 | 38 | 11 | KGKSYDALATF | 0.915542 | 0.09 | -0.13511 | Non-Toxin | 100.00% (10/10) | DNA-directed RNA polymerase |
| 375 | B*35:01 | 42 | 50 | 9 | KPYESKVFF | 0.76041 | 0.09 | -0.16131 | Non-Toxin | 90.00% (9/10) | DNA-directed RNA polymerase |
| 376 | A*11:01 | 41 | 49 | 9 | NIFKEVMTK | 0.76758 | 0.09 | -0.16816 | Non-Toxin | 100.00% (10/10) | DNA-directed RNA polymerase |
| 377 | A*68:01 | 31 | 40 | 10 | NVIGGVFINK | 0.887608 | 0.09 | 0.36924 | Non-Toxin | 100.00% (10/10) | DNA-directed RNA polymerase |
| 378 | B*57:01 | 34 | 44 | 11 | RSYYKDDISYF | 0.926193 | 0.09 | -0.18822 | Non-Toxin | 100.00% (10/10) | DNA-directed RNA polymerase |
| 379 | A*11:01 | 14 | 24 | 11 | SVSPPNVLPTR | 0.774259 | 0.09 | -0.03888 | Non-Toxin | 90.00% (9/10) | DNA-directed RNA polymerase |
| 380 | B*35:01 | 22 | 30 | 9 | TPDQLKGFY | 0.773885 | 0.09 | -0.22616 | Non-Toxin | 100.00% (10/10) | DNA-directed RNA polymerase |
| 381 | B*07:02 | 29 | 37 | 9 | VPNVIGGVF | 0.781965 | 0.09 | 0.25366 | Non-Toxin | 100.00% (10/10) | DNA-directed RNA polymerase |
| 382 | B*40:01 | 13 | 20 | 8 | YEYSNPKL | 0.844526 | 0.09 | -0.36641 | Non-Toxin | 100.00% (10/10) | DNA-directed RNA polymerase |
| 383 | B*35:01 | 27 | 36 | 10 | YPDQVKISKM | 0.769711 | 0.09 | -0.4195 | Non-Toxin | 100.00% (10/10) | DNA-directed RNA polymerase |
| 384 | A*03:01 | 34 | 42 | 9 | ILYDPETDK | 0.784949 | 0.1 | 0.15029 | Non-Toxin | 100.00% (10/10) | DNA-directed RNA polymerase |
| 385 | B*58:01 | 14 | 23 | 10 | MTFSDIQKEF | 0.828738 | 0.1 | -0.20981 | Non-Toxin | 100.00% (10/10) | DNA-directed RNA polymerase |
| 386 | A*68:01 | 16 | 24 | 9 | NSTSIFSRK | 0.881612 | 0.1 | -0.02345 | Non-Toxin | 100.00% (10/10) | DNA-directed RNA polymerase |
| **#** | **allele** | **start** | **end** | **length** | **peptide** | **score** | **percentile_rank** | **Immunogenicity** | **Toxicity** | **Conservancy** | **Protein** |
| 387 | B*44:03 | 2 | 10 | 9 | AEFEDQLVF | 0.983805 | 0.01 | 0.06607 | Non-Toxin | 99.66% (580/582) | Telomere-binding protein |
| 388 | B*40:01 | 2 | 10 | 9 | AEFEDQLVF | 0.974324 | 0.01 | 0.06607 | Non-Toxin | 99.66% (580/582) | Telomere-binding protein |
| 389 | B*44:02 | 2 | 10 | 9 | AEFEDQLVF | 0.954767 | 0.01 | 0.06607 | Non-Toxin | 99.66% (580/582) | Telomere-binding protein |
| 390 | A*33:01 | 10 | 18 | 9 | DVLKTRLFR | 0.92971 | 0.01 | -0.07504 | Non-Toxin | 99.83% (581/582) | Telomere-binding protein |
| 391 | A*33:01 | 34 | 42 | 9 | DYFTRVTKR | 0.970491 | 0.01 | 0.07308 | Non-Toxin | 99.66% (580/582) | Telomere-binding protein |
| 392 | A*33:01 | 7 | 15 | 9 | ELIDVLKTR | 0.866345 | 0.01 | -0.06404 | Non-Toxin | 100.00% (582/582) | Telomere-binding protein |
| 393 | B*08:01 | 10 | 18 | 9 | FMYTKHSML | 0.934619 | 0.01 | -0.38391 | Non-Toxin | 99.83% (581/582) | Telomere-binding protein |
| 394 | A*68:02 | 22 | 30 | 9 | FTAKINEMV | 0.937921 | 0.01 | -0.09889 | Non-Toxin | 99.66% (580/582) | Telomere-binding protein |
| 395 | A*31:01 | 9 | 17 | 9 | KVLAARLKR | 0.937614 | 0.01 | -0.01277 | Non-Toxin | 100.00% (582/582) | Telomere-binding protein |
| 396 | A*33:01 | 21 | 29 | 9 | QVKDNIISR | 0.864474 | 0.01 | 0.08696 | Non-Toxin | 99.83% (581/582) | Telomere-binding protein |
| 397 | A*03:01 | 15 | 23 | 9 | RLFRSPQVK | 0.975832 | 0.01 | -0.1551 | Non-Toxin | 99.83% (581/582) | Telomere-binding protein |
| 398 | A*30:01 | 15 | 23 | 9 | RLFRSPQVK | 0.830003 | 0.01 | -0.1551 | Non-Toxin | 99.83% (581/582) | Telomere-binding protein |
| 399 | B*15:01 | 14 | 22 | 9 | RLKRSATQF | 0.944417 | 0.01 | -0.17711 | Non-Toxin | 100.00% (582/582) | Telomere-binding protein |
| 400 | A*02:03 | 31 | 39 | 9 | RLYDYFTRV | 0.989846 | 0.01 | 0.19072 | Non-Toxin | 99.83% (581/582) | Telomere-binding protein |
| 401 | A*02:01 | 31 | 39 | 9 | RLYDYFTRV | 0.989046 | 0.01 | 0.19072 | Non-Toxin | 99.83% (581/582) | Telomere-binding protein |
| 402 | A*02:06 | 31 | 39 | 9 | RLYDYFTRV | 0.969493 | 0.01 | 0.19072 | Non-Toxin | 99.83% (581/582) | Telomere-binding protein |
| 403 | A*03:01 | 31 | 41 | 11 | RLYDYFTRVTK | 0.974287 | 0.01 | 0.27556 | Non-Toxin | 99.83% (581/582) | Telomere-binding protein |
| 404 | B*15:01 | 20 | 29 | 10 | TQFNFNGHTY | 0.942285 | 0.01 | 0.22107 | Non-Toxin | 100.00% (582/582) | Telomere-binding protein |
| 405 | A*68:01 | 7 | 15 | 9 | ELIDVLKTR | 0.961953 | 0.02 | -0.06404 | Non-Toxin | 100.00% (582/582) | Telomere-binding protein |
| 406 | B*08:01 | 10 | 17 | 8 | FMYTKHSM | 0.867224 | 0.02 | -0.28131 | Non-Toxin | 99.83% (581/582) | Telomere-binding protein |
| 407 | A*68:01 | 21 | 29 | 9 | QVKDNIISR | 0.965789 | 0.02 | 0.08696 | Non-Toxin | 99.83% (581/582) | Telomere-binding protein |
| 408 | A*31:01 | 21 | 29 | 9 | QVKDNIISR | 0.904855 | 0.02 | 0.08696 | Non-Toxin | 99.83% (581/582) | Telomere-binding protein |
| 409 | A*26:01 | 25 | 33 | 9 | NIISRTRLY | 0.83762 | 0.03 | 0.00087 | Non-Toxin | 99.83% (581/582) | Telomere-binding protein |
| 410 | A*23:01 | 36 | 44 | 9 | SFYEDIAEF | 0.870243 | 0.03 | 0.33795 | Non-Toxin | 99.83% (581/582) | Telomere-binding protein |
| 411 | A*03:01 | 17 | 25 | 9 | ALKAYFTAK | 0.872584 | 0.04 | 0.13159 | Non-Toxin | 99.48% (579/582) | Telomere-binding protein |
| 412 | A*32:01 | 14 | 22 | 9 | RLKRSATQF | 0.683726 | 0.04 | -0.17711 | Non-Toxin | 100.00% (582/582) | Telomere-binding protein |
| 413 | A*24:02 | 28 | 37 | 10 | RYSKKFQESF | 0.876034 | 0.04 | -0.4879 | Non-Toxin | 99.83% (581/582) | Telomere-binding protein |
| 414 | A*30:01 | 23 | 31 | 9 | SSRVDRYSK | 0.713668 | 0.04 | 0.02888 | Non-Toxin | 99.66% (580/582) | Telomere-binding protein |
| 415 | A*30:02 | 20 | 29 | 10 | TQFNFNGHTY | 0.734645 | 0.04 | 0.22107 | Non-Toxin | 100.00% (582/582) | Telomere-binding protein |
| 416 | A*26:01 | 3 | 12 | 10 | ETVKMGAFMY | 0.775091 | 0.05 | -0.30857 | Non-Toxin | 99.66% (580/582) | Telomere-binding protein |
| 417 | A*02:03 | 10 | 18 | 9 | FMYTKHSML | 0.842432 | 0.05 | -0.38391 | Non-Toxin | 99.83% (581/582) | Telomere-binding protein |
| 418 | A*02:01 | 14 | 22 | 9 | ILISLINSL | 0.86778 | 0.05 | -0.11091 | Non-Toxin | 99.83% (581/582) | Telomere-binding protein |
| 419 | A*02:03 | 14 | 22 | 9 | ILISLINSL | 0.808331 | 0.05 | -0.11091 | Non-Toxin | 99.83% (581/582) | Telomere-binding protein |
| 420 | B*53:01 | 4 | 12 | 9 | IPDELIDVL | 0.718116 | 0.05 | 0.26527 | Non-Toxin | 100.00% (582/582) | Telomere-binding protein |
| 421 | A*03:01 | 13 | 23 | 11 | KTRLFRSPQVK | 0.851995 | 0.05 | -0.07514 | Non-Toxin | 99.83% (581/582) | Telomere-binding protein |
| 422 | A*30:01 | 13 | 23 | 11 | KTRLFRSPQVK | 0.70184 | 0.05 | -0.07514 | Non-Toxin | 99.83% (581/582) | Telomere-binding protein |
| 423 | A*68:01 | 8 | 16 | 9 | LVFNSISAR | 0.925269 | 0.05 | -0.12109 | Non-Toxin | 99.48% (579/582) | Telomere-binding protein |
| 424 | A*23:01 | 28 | 37 | 10 | RYSKKFQESF | 0.764602 | 0.05 | -0.4879 | Non-Toxin | 99.83% (581/582) | Telomere-binding protein |

| | | | | | | | | | | | |
|---|---|---|---|---|---|---|---|---|---|---|---|
| 425 | A*24:02 | 36 | 44 | 9 | SFYEDIAEF | 0.81721 | 0.05 | 0.33795 | Non-Toxin | 99.83% (581/582) | Telomere-binding protein |
| 426 | A*33:01 | 4 | 14 | 11 | DISDVKVLAAR | 0.711811 | 0.06 | -0.11554 | Non-Toxin | 100.00% (582/582) | Telomere-binding protein |
| 427 | A*26:01 | 3 | 11 | 9 | ETVKMGAFM | 0.70488 | 0.06 | -0.24128 | Non-Toxin | 99.66% (580/582) | Telomere-binding protein |
| 428 | A*30:01 | 31 | 41 | 11 | RLYDYFTRVTK | 0.643003 | 0.06 | 0.27556 | Non-Toxin | 99.83% (581/582) | Telomere-binding protein |
| 429 | A*30:01 | 34 | 42 | 9 | VTRKCPQKK | 0.674778 | 0.06 | -0.4869 | Non-Toxin | 100.00% (582/582) | Telomere-binding protein |
| 430 | A*68:01 | 10 | 18 | 9 | DVLKTRLFR | 0.910916 | 0.07 | -0.07504 | Non-Toxin | 99.83% (581/582) | Telomere-binding protein |
| 431 | A*68:01 | 28 | 36 | 9 | EMVDELVTR | 0.91305 | 0.07 | 0.1803 | Non-Toxin | 99.66% (580/582) | Telomere-binding protein |
| 432 | A*33:01 | 28 | 36 | 9 | EMVDELVTR | 0.708943 | 0.07 | 0.1803 | Non-Toxin | 99.66% (580/582) | Telomere-binding protein |
| 433 | B*44:03 | 34 | 44 | 11 | QESFYEDIAEF | 0.818557 | 0.07 | 0.3964 | Non-Toxin | 99.83% (581/582) | Telomere-binding protein |
| 434 | B*57:01 | 29 | 37 | 9 | YSKKFQESF | 0.944012 | 0.07 | -0.2942 | Non-Toxin | 99.83% (581/582) | Telomere-binding protein |
| 435 | A*33:01 | 6 | 16 | 11 | DQLVFNSISAR | 0.659602 | 0.08 | -0.00694 | Non-Toxin | 99.48% (579/582) | Telomere-binding protein |
| 436 | A*03:01 | 6 | 14 | 9 | KMGAFMYTK | 0.800283 | 0.08 | 0.01863 | Non-Toxin | 99.83% (581/582) | Telomere-binding protein |
| 437 | A*24:02 | 10 | 18 | 9 | NYGGILISL | 0.736025 | 0.08 | 0.17992 | Non-Toxin | 99.83% (581/582) | Telomere-binding protein |
| 438 | A*33:01 | 33 | 42 | 10 | YDYFTRVTKR | 0.681938 | 0.08 | 0.15042 | Non-Toxin | 99.66% (580/582) | Telomere-binding protein |
| 439 | A*68:01 | 6 | 16 | 11 | DQLVFNSISAR | 0.886961 | 0.09 | -0.00694 | Non-Toxin | 99.48% (579/582) | Telomere-binding protein |
| 440 | B*40:01 | 31 | 39 | 9 | LENDKIEDL | 0.831321 | 0.09 | 0.03296 | Non-Toxin | 99.66% (580/582) | Telomere-binding protein |
| 441 | A*31:01 | 8 | 16 | 9 | LVFNSISAR | 0.752055 | 0.09 | -0.12109 | Non-Toxin | 99.48% (579/582) | Telomere-binding protein |
| 442 | B*58:01 | 29 | 37 | 9 | YSKKFQESF | 0.852768 | 0.09 | -0.2942 | Non-Toxin | 99.83% (581/582) | Telomere-binding protein |
| 443 | A*68:01 | 44 | 52 | 9 | ESSIYVILK | 0.882503 | 0.1 | 0.22132 | Non-Toxin | 99.83% (581/582) | Telomere-binding protein |
| 444 | A*68:02 | 50 | 58 | 9 | EVRIPVDLV | 0.668298 | 0.1 | 0.19102 | Non-Toxin | 99.66% (580/582) | Telomere-binding protein |
| 445 | A*02:06 | 14 | 22 | 9 | ILISLINSL | 0.732618 | 0.1 | -0.11091 | Non-Toxin | 99.83% (581/582) | Telomere-binding protein |
| 446 | B*35:01 | 4 | 12 | 9 | IPDELIDVL | 0.749922 | 0.1 | 0.26527 | Non-Toxin | 100.00% (582/582) | Telomere-binding protein |
| 447 | A*68:02 | 18 | 26 | 9 | LTNAISSRV | 0.653604 | 0.1 | -0.09824 | Non-Toxin | 99.83% (581/582) | Telomere-binding protein |
| 448 | A*11:01 | 29 | 37 | 9 | MVDELVTRK | 0.758585 | 0.1 | 0.19901 | Non-Toxin | 99.66% (580/582) | Telomere-binding protein |
| 449 | B*44:02 | 34 | 44 | 11 | QESFYEDIAEF | 0.695334 | 0.1 | 0.3964 | Non-Toxin | 99.83% (581/582) | Telomere-binding protein |
| 450 | A*31:01 | 19 | 29 | 11 | SPQVKDNIISR | 0.735926 | 0.1 | -0.04982 | Non-Toxin | 99.83% (581/582) | Telomere-binding protein |
| **#** | **allele** | **start** | **end** | **length** | **peptide** | **score** | **percentile_rank** | **Immunogenicity** | **Toxicity** | **Conservancy** | **Protein** |
| 451 | B*35:01 | 20 | 28 | 9 | YAMELLTGY | 0.982787 | 0.01 | 0.07507 | Non-Toxin | 99.83% (580/581) | Profilin |
| 452 | B*51:01 | 49 | 57 | 9 | IPLITNHNI | 0.915576 | 0.02 | 0.18555 | Non-Toxin | 99.48% (578/581) | Profilin |
| 453 | B*51:01 | 35 | 43 | 9 | IPNRTFAKI | 0.892859 | 0.02 | 0.105 | Non-Toxin | 99.83% (580/581) | Profilin |
| 454 | B*51:01 | 44 | 52 | 9 | NPGEVIPLI | 0.896042 | 0.02 | 0.26139 | Non-Toxin | 99.48% (578/581) | Profilin |
| 455 | A*26:01 | 20 | 28 | 9 | YAMELLTGY | 0.876874 | 0.03 | 0.07507 | Non-Toxin | 99.83% (580/581) | Profilin |
| 456 | A*11:01 | 21 | 29 | 9 | AIVDYKTTK | 0.866982 | 0.04 | -0.11544 | Non-Toxin | 99.31% (577/581) | Profilin |
| 457 | A*26:01 | 47 | 55 | 9 | EVIPLITNH | 0.809492 | 0.04 | 0.1755 | Non-Toxin | 99.48% (578/581) | Profilin |
| 458 | A*02:01 | 14 | 22 | 9 | LMDENTYAM | 0.864084 | 0.05 | 0.15793 | Non-Toxin | 100.00% (581/581) | Profilin |
| 459 | A*30:01 | 38 | 46 | 9 | RTHTALIFL | 0.70211 | 0.05 | 0.25794 | Non-Toxin | 99.48% (578/581) | Profilin |
| 460 | A*01:01 | 10 | 20 | 11 | YTNSLMDENTY | 0.807938 | 0.06 | -0.2173 | Non-Toxin | 99.83% (580/581) | Profilin |
| 461 | A*02:01 | 13 | 21 | 9 | SLMDENTYA | 0.819223 | 0.07 | 0.08733 | Non-Toxin | 100.00% (581/581) | Profilin |
| 462 | A*02:03 | 13 | 21 | 9 | SLMDENTYA | 0.772016 | 0.07 | 0.08733 | Non-Toxin | 100.00% (581/581) | Profilin |
| 463 | B*51:01 | 28 | 35 | 8 | YAPVSPIV | 0.707726 | 0.07 | -0.02128 | Non-Toxin | 99.83% (580/581) | Profilin |
| 464 | B*51:01 | 28 | 36 | 9 | YAPVSPIVI | 0.696762 | 0.07 | 0.00284 | Non-Toxin | 99.83% (580/581) | Profilin |
| 465 | A*02:03 | 45 | 53 | 9 | FLMGKPTTS | 0.745553 | 0.08 | -0.1879 | Non-Toxin | 99.66% (579/581) | Profilin |
| 466 | A*68:01 | 30 | 38 | 9 | NVLAAIPNR | 0.893638 | 0.08 | 0.18601 | Non-Toxin | 99.83% (580/581) | Profilin |
| 467 | A*68:01 | 30 | 38 | 9 | PVSPIVIAR | 0.894476 | 0.08 | 0.23878 | Non-Toxin | 99.83% (580/581) | Profilin |
| 468 | A*33:01 | 30 | 38 | 9 | NVLAAIPNR | 0.638681 | 0.09 | 0.18601 | Non-Toxin | 99.83% (580/581) | Profilin |
| 469 | A*11:01 | 34 | 42 | 9 | AIPNRTFAK | 0.747748 | 0.1 | 0.19849 | Non-Toxin | 99.83% (580/581) | Profilin |
| 470 | A*31:01 | 29 | 38 | 10 | APVSPIVIAR | 0.73873 | 0.1 | 0.13977 | Non-Toxin | 99.83% (580/581) | Profilin |
| 471 | A*68:02 | 47 | 57 | 11 | EVIPLITNHNI | 0.658038 | 0.1 | 0.20607 | Non-Toxin | 99.48% (578/581) | Profilin |
| 472 | A*02:06 | 14 | 22 | 9 | LMDENTYAM | 0.734083 | 0.1 | 0.15793 | Non-Toxin | 100.00% (581/581) | Profilin |
| 473 | B*53:01 | 44 | 52 | 9 | NPGEVIPLI | 0.566588 | 0.1 | 0.26139 | Non-Toxin | 99.48% (578/581) | Profilin |
| 474 | A*02:06 | 13 | 21 | 9 | SLMDENTYA | 0.737021 | 0.1 | 0.08733 | Non-Toxin | 100.00% (581/581) | Profilin |
| 475 | A*68:01 | 51 | 59 | 9 | TTSRRDVYR | 0.87996 | 0.1 | 0.10234 | Non-Toxin | 100.00% (581/581) | Profilin |
| 476 | A*03:01 | 21 | 29 | 9 | AIVDYKTTK | 0.770577 | 0.11 | -0.11544 | Non-Toxin | 99.31% (577/581) | Profilin |

| | | | | | | | | | | | |
|---|---|---|---|---|---|---|---|---|---|---|---|
| 477 | A*26:01 | 18 | 28 | 11 | NTYAMELLTGY | 0.501916 | 0.11 | -0.00401 | Non-Toxin | 99.83% (580/581) | Profilin |
| 478 | A*11:01 | 33 | 42 | 10 | AAIPNRTFAK | 0.725752 | 0.12 | 0.23434 | Non-Toxin | 99.83% (580/581) | Profilin |
| 479 | A*68:01 | 29 | 38 | 10 | APVSPIVIAR | 0.871398 | 0.12 | 0.13977 | Non-Toxin | 99.83% (580/581) | Profilin |
| 480 | A*68:02 | 27 | 35 | 9 | TTKNVLAAI | 0.631859 | 0.12 | 0.00913 | Non-Toxin | 99.83% (580/581) | Profilin |
| 481 | A*31:01 | 47 | 55 | 9 | MGKPTTSRR | 0.697499 | 0.13 | -0.1162 | Non-Toxin | 100.00% (581/581) | Profilin |
| 482 | A*33:01 | 47 | 55 | 9 | MGKPTTSRR | 0.543366 | 0.13 | -0.1162 | Non-Toxin | 100.00% (581/581) | Profilin |
| 483 | B*53:01 | 20 | 28 | 9 | YAMELLTGY | 0.492822 | 0.13 | 0.07507 | Non-Toxin | 99.83% (580/581) | Profilin |
| 484 | B*08:01 | 36 | 43 | 8 | IARTHTAL | 0.513058 | 0.14 | 0.15692 | Non-Toxin | 99.66% (579/581) | Profilin |
| 485 | B*07:02 | 49 | 57 | 9 | KPTTSRRDV | 0.661691 | 0.14 | -0.00408 | Non-Toxin | 100.00% (581/581) | Profilin |
| 486 | B*58:01 | 32 | 40 | 9 | LAAIPNRTF | 0.755926 | 0.14 | 0.19609 | Non-Toxin | 99.83% (580/581) | Profilin |
| 487 | A*33:01 | 30 | 38 | 9 | PVSPIVIAR | 0.530721 | 0.15 | 0.23878 | Non-Toxin | 99.83% (580/581) | Profilin |
| 488 | B*15:01 | 20 | 28 | 9 | YAMELLTGY | 0.68142 | 0.15 | 0.07507 | Non-Toxin | 99.83% (580/581) | Profilin |
| 489 | A*68:01 | 28 | 38 | 11 | YAPVSPIVIAR | 0.845383 | 0.15 | 0.16968 | Non-Toxin | 99.83% (580/581) | Profilin |
| 490 | A*30:02 | 20 | 28 | 9 | YAMELLTGY | 0.511031 | 0.16 | 0.07507 | Non-Toxin | 99.83% (580/581) | Profilin |
| **#** | **allele** | **start** | **end** | **length** | **peptide** | **score** | **percentile_rank** | **Immunogenicity** | **Toxicity** | **Conservancy** | **Protein** |
| 491 | A*33:01 | 32 | 40 | 9 | DIYGLMKER | 0.870262 | 0.01 | -0.2667 | Non-Toxin | 100.00% (9/9) | Intermediate transcription factor 3 |
| 492 | A*68:01 | 48 | 56 | 9 | DVYGVSNFK | 0.985274 | 0.01 | -0.01969 | Non-Toxin | 100.00% (9/9) | Intermediate transcription factor 3 |
| 493 | A*68:02 | 20 | 28 | 9 | EVANVVNHV | 0.995135 | 0.01 | 0.09869 | Non-Toxin | 100.00% (9/9) | Intermediate transcription factor 3 |
| 494 | A*68:01 | 18 | 27 | 10 | QTIEIIFTNR | 0.974101 | 0.01 | 0.54233 | Non-Toxin | 100.00% (9/9) | Intermediate transcription factor 3 |
| 495 | A*24:02 | 14 | 22 | 9 | RYARTIFNF | 0.99209 | 0.01 | 0.32288 | Non-Toxin | 100.00% (9/9) | Intermediate transcription factor 3 |
| 496 | A*23:01 | 14 | 22 | 9 | RYARTIFNF | 0.987658 | 0.01 | 0.32288 | Non-Toxin | 100.00% (9/9) | Intermediate transcription factor 3 |
| 497 | A*11:01 | 36 | 44 | 9 | STMHIILSK | 0.993167 | 0.01 | 0.12441 | Non-Toxin | 100.00% (9/9) | Intermediate transcription factor 3 |
| 498 | A*03:01 | 36 | 44 | 9 | STMHIILSK | 0.975706 | 0.01 | 0.12441 | Non-Toxin | 100.00% (9/9) | Intermediate transcription factor 3 |
| 499 | A*30:01 | 36 | 44 | 9 | STMHIILSK | 0.853983 | 0.01 | 0.12441 | Non-Toxin | 100.00% (9/9) | Intermediate transcription factor 3 |
| 500 | A*23:01 | 1 | 9 | 9 | VYYNLFLLF | 0.986203 | 0.01 | 0.07585 | Non-Toxin | 100.00% (9/9) | Intermediate transcription factor 3 |
| 501 | A*24:02 | 1 | 9 | 9 | VYYNLFLLF | 0.964385 | 0.01 | 0.07585 | Non-Toxin | 100.00% (9/9) | Intermediate transcription factor 3 |
| 502 | A*02:01 | 33 | 41 | 9 | ALDEKLFLI | 0.974015 | 0.02 | -0.02017 | Non-Toxin | 100.00% (9/9) | Intermediate transcription factor 3 |
| 503 | A*01:01 | 25 | 34 | 10 | ISCDEIGDIY | 0.928486 | 0.02 | 0.3603 | Non-Toxin | 100.00% (9/9) | Intermediate transcription factor 3 |
| 504 | B*44:03 | 45 | 55 | 11 | AEHDVYGVSNF | 0.919549 | 0.03 | -0.00212 | Non-Toxin | 100.00% (9/9) | Intermediate transcription factor 3 |
| 505 | B*44:02 | 45 | 55 | 11 | AEHDVYGVSNF | 0.891188 | 0.03 | -0.00212 | Non-Toxin | 100.00% (9/9) | Intermediate transcription factor 3 |
| 506 | B*44:02 | 18 | 26 | 9 | DEDSKIKFF | 0.849535 | 0.03 | -0.35759 | Non-Toxin | 100.00% (9/9) | Intermediate transcription factor 3 |
| 507 | A*03:01 | 11 | 19 | 9 | KVNYGEIKK | 0.896046 | 0.03 | 0.10775 | Non-Toxin | 100.00% (9/9) | Intermediate transcription factor 3 |
| 508 | A*02:01 | 3 | 11 | 9 | NLFTFLHEI | 0.929389 | 0.03 | 0.26642 | Non-Toxin | 100.00% (9/9) | Intermediate transcription factor 3 |
| 509 | A*02:03 | 3 | 11 | 9 | NLFTFLHEI | 0.87675 | 0.03 | 0.26642 | Non-Toxin | 100.00% (9/9) | Intermediate transcription factor 3 |
| 510 | A*02:06 | 33 | 41 | 9 | ALDEKLFLI | 0.905321 | 0.04 | -0.02017 | Non-Toxin | 100.00% (9/9) | Intermediate transcription factor 3 |
| 511 | A*33:01 | 2 | 11 | 10 | DIINHSIVTR | 0.76861 | 0.04 | 0.08989 | Non-Toxin | 100.00% (9/9) | Intermediate transcription factor 3 |
| 512 | A*68:01 | 46 | 56 | 11 | EHDVYGVSNFK | 0.942076 | 0.04 | 0.02555 | Non-Toxin | 100.00% (9/9) | Intermediate transcription factor 3 |
| 513 | A*68:02 | 20 | 29 | 10 | EVANVVNHVL | 0.839505 | 0.04 | 0.13192 | Non-Toxin | 100.00% (9/9) | Intermediate transcription factor 3 |
| 514 | A*31:01 | 3 | 11 | 9 | IINHSIVTR | 0.854631 | 0.04 | 0.05215 | Non-Toxin | 100.00% (9/9) | Intermediate transcription factor 3 |
| 515 | A*11:01 | 11 | 19 | 9 | KVNYGEIKK | 0.871364 | 0.04 | 0.10775 | Non-Toxin | 100.00% (9/9) | Intermediate transcription factor 3 |
| 516 | A*03:01 | 3 | 12 | 10 | LVYCDIQLTK | 0.876516 | 0.04 | 3.00E-05 | Non-Toxin | 100.00% (9/9) | Intermediate transcription factor 3 |
| 517 | A*01:01 | 42 | 52 | 11 | SSEDMFDNIVY | 0.875577 | 0.04 | 0.14977 | Non-Toxin | 100.00% (9/9) | Intermediate transcription factor 3 |
| 518 | A*68:01 | 36 | 44 | 9 | STMHIILSK | 0.945591 | 0.04 | 0.12441 | Non-Toxin | 100.00% (9/9) | Intermediate transcription factor 3 |
| 519 | B*44:03 | 18 | 26 | 9 | DEDSKIKFF | 0.854262 | 0.05 | -0.35759 | Non-Toxin | 100.00% (9/9) | Intermediate transcription factor 3 |
| 520 | A*68:01 | 2 | 11 | 10 | DIINHSIVTR | 0.926668 | 0.05 | 0.08989 | Non-Toxin | 100.00% (9/9) | Intermediate transcription factor 3 |
| 521 | A*01:01 | 30 | 38 | 9 | FSGKKSDEY | 0.839766 | 0.05 | -0.49451 | Non-Toxin | 100.00% (9/9) | Intermediate transcription factor 3 |
| 522 | A*03:01 | 47 | 56 | 10 | HDVYGVSNFK | 0.848086 | 0.05 | -0.00991 | Non-Toxin | 100.00% (9/9) | Intermediate transcription factor 3 |
| 523 | B*40:01 | 8 | 16 | 9 | IETESVDRL | 0.938178 | 0.05 | 0.04007 | Non-Toxin | 100.00% (9/9) | Intermediate transcription factor 3 |
| 524 | A*03:01 | 2 | 12 | 11 | KLVYCDIQLTK | 0.863062 | 0.05 | 0.01266 | Non-Toxin | 100.00% (9/9) | Intermediate transcription factor 3 |
| 525 | A*68:02 | 4 | 13 | 10 | NSIDIETESV | 0.806695 | 0.05 | 0.317 | Non-Toxin | 100.00% (9/9) | Intermediate transcription factor 3 |
| 526 | A*26:01 | 20 | 29 | 10 | TVHGGTNANY | 0.767477 | 0.05 | 0.13855 | Non-Toxin | 100.00% (9/9) | Intermediate transcription factor 3 |
| 527 | A*68:01 | 32 | 40 | 9 | DIYGLMKER | 0.920526 | 0.06 | -0.2667 | Non-Toxin | 100.00% (9/9) | Intermediate transcription factor 3 |
| 528 | A*02:06 | 13 | 21 | 9 | IQNEYFKEV | 0.860651 | 0.06 | 0.08175 | Non-Toxin | 11.11% (1/9) | Intermediate transcription factor 3 |

| | | | | | | | | | | | |
|---|---|---|---|---|---|---|---|---|---|---|---|
| 529 | B*58:01 | 44 | 52 | 9 | ISSRTVEIF | 0.907077 | 0.06 | 0.2373 | Non-Toxin | 100.00% (9/9) | Intermediate transcription factor 3 |
| 530 | A*02:06 | 3 | 11 | 9 | NLFTFLHEI | 0.839567 | 0.06 | 0.26642 | Non-Toxin | 100.00% (9/9) | Intermediate transcription factor 3 |
| 531 | A*30:02 | 37 | 45 | 9 | RSLEDCTNY | 0.698124 | 0.06 | 0.09698 | Non-Toxin | 100.00% (9/9) | Intermediate transcription factor 3 |
| 532 | A*01:01 | 26 | 34 | 9 | SCDEIGDIY | 0.823496 | 0.06 | 0.36593 | Non-Toxin | 100.00% (9/9) | Intermediate transcription factor 3 |
| 533 | A*11:01 | 36 | 46 | 11 | STMHIILSKDK | 0.837064 | 0.06 | -0.10082 | Non-Toxin | 100.00% (9/9) | Intermediate transcription factor 3 |
| 534 | A*24:02 | 49 | 57 | 9 | VYGVSNFKI | 0.807805 | 0.06 | -0.14185 | Non-Toxin | 100.00% (9/9) | Intermediate transcription factor 3 |
| 535 | A*26:01 | 20 | 28 | 9 | EVANVVNHV | 0.641397 | 0.07 | 0.09869 | Non-Toxin | 100.00% (9/9) | Intermediate transcription factor 3 |
| 536 | A*68:01 | 47 | 56 | 10 | HDVYGVSNFK | 0.901838 | 0.07 | -0.00991 | Non-Toxin | 100.00% (9/9) | Intermediate transcription factor 3 |
| 537 | A*03:01 | 34 | 44 | 11 | KDSTMHIILSK | 0.816684 | 0.07 | -0.00528 | Non-Toxin | 100.00% (9/9) | Intermediate transcription factor 3 |
| 538 | A*02:03 | 43 | 52 | 10 | KMAEHDVYGV | 0.741657 | 0.08 | 0.22209 | Non-Toxin | 100.00% (9/9) | Intermediate transcription factor 3 |
| 539 | A*11:01 | 3 | 12 | 10 | LVYCDIQLTK | 0.798063 | 0.08 | 3.00E-05 | Non-Toxin | 100.00% (9/9) | Intermediate transcription factor 3 |
| 540 | A*11:01 | 47 | 56 | 10 | RTVEIFESEK | 0.801792 | 0.08 | 0.37088 | Non-Toxin | 100.00% (9/9) | Intermediate transcription factor 3 |
| 541 | A*23:01 | 2 | 9 | 8 | YYNLFLLF | 0.718242 | 0.08 | 0.08094 | Non-Toxin | 100.00% (9/9) | Intermediate transcription factor 3 |
| 542 | A*02:03 | 33 | 41 | 9 | ALDEKLFLI | 0.727538 | 0.09 | -0.02017 | Non-Toxin | 100.00% (9/9) | Intermediate transcription factor 3 |
| 543 | B*44:03 | 9 | 17 | 9 | FEDIIQNEY | 0.772058 | 0.1 | 0.21472 | Non-Toxin | 11.11% (1/9) | Intermediate transcription factor 3 |
| 544 | A*02:03 | 4 | 12 | 9 | FIKSLDHTV | 0.707385 | 0.1 | -0.17641 | Non-Toxin | 22.22% (2/9) | Intermediate transcription factor 3 |
| 545 | A*02:03 | 13 | 21 | 9 | IQNEYFKEV | 0.699147 | 0.1 | 0.08175 | Non-Toxin | 11.11% (1/9) | Intermediate transcription factor 3 |
| 546 | A*11:01 | 34 | 44 | 11 | KDSTMHIILSK | 0.75503 | 0.1 | -0.00528 | Non-Toxin | 100.00% (9/9) | Intermediate transcription factor 3 |
| 547 | B*57:01 | 41 | 51 | 11 | TTVRSNINQPW | 0.910669 | 0.1 | -0.08265 | Non-Toxin | 100.00% (9/9) | Intermediate transcription factor 3 |
| 548 | A*23:01 | 49 | 57 | 9 | VYGVSNFKI | 0.629465 | 0.1 | -0.14185 | Non-Toxin | 100.00% (9/9) | Intermediate transcription factor 3 |
| **#** | **allele** | **start** | **end** | **length** | **peptide** | **score** | **percentile_rank** | **Immunogenicity** | **Toxicity** | **Conservancy** | **Protein** |
| 549 | B*44:03 | 43 | 51 | 9 | DENELFKHY | 0.990298 | 0.01 | 0.03495 | Non-Toxin | 100.00% (1/1) | Probable host range protein 2 |
| 550 | B*44:02 | 43 | 51 | 9 | DENELFKHY | 0.970961 | 0.01 | 0.03495 | Non-Toxin | 100.00% (1/1) | Probable host range protein 2 |
| 551 | A*32:01 | 11 | 19 | 9 | KVNKVDYTL | 0.840145 | 0.01 | -0.13846 | Non-Toxin | 100.00% (1/1) | Probable host range protein 2 |
| 552 | B*51:01 | 13 | 21 | 9 | SPYIEHPLI | 0.944556 | 0.01 | 0.24483 | Non-Toxin | 100.00% (1/1) | Probable host range protein 2 |
| 553 | B*51:01 | 52 | 60 | 9 | YPYISLNMI | 0.977713 | 0.01 | -0.14688 | Non-Toxin | 100.00% (1/1) | Probable host range protein 2 |
| 554 | A*11:01 | 26 | 34 | 9 | AVIHLYNKK | 0.895191 | 0.02 | -0.06999 | Non-Toxin | 100.00% (1/1) | Probable host range protein 2 |
| 555 | B*44:03 | 43 | 52 | 10 | DENELFKHYY | 0.938701 | 0.03 | 0.02399 | Non-Toxin | 100.00% (1/1) | Probable host range protein 2 |
| 556 | B*44:02 | 43 | 52 | 10 | DENELFKHYY | 0.871278 | 0.03 | 0.02399 | Non-Toxin | 100.00% (1/1) | Probable host range protein 2 |
| 557 | B*08:01 | 14 | 22 | 9 | DIALRNLQL | 0.832428 | 0.03 | -0.03119 | Non-Toxin | 100.00% (1/1) | Probable host range protein 2 |
| 558 | B*08:01 | 29 | 37 | 9 | HLYNKKTEI | 0.846055 | 0.03 | -0.32945 | Non-Toxin | 100.00% (1/1) | Probable host range protein 2 |
| 559 | A*30:02 | 11 | 21 | 11 | KVNKVDYTLYY | 0.762121 | 0.03 | -0.13588 | Non-Toxin | 100.00% (1/1) | Probable host range protein 2 |
| 560 | B*44:03 | 41 | 51 | 11 | SDDENELFKHY | 0.929048 | 0.03 | 0.13545 | Non-Toxin | 100.00% (1/1) | Probable host range protein 2 |
| 561 | B*44:02 | 41 | 51 | 11 | SDDENELFKHY | 0.883271 | 0.03 | 0.13545 | Non-Toxin | 100.00% (1/1) | Probable host range protein 2 |
| 562 | A*02:06 | 22 | 30 | 9 | VIYEAVIHL | 0.920983 | 0.03 | 0.30773 | Non-Toxin | 100.00% (1/1) | Probable host range protein 2 |
| 563 | A*02:01 | 22 | 30 | 9 | VIYEAVIHL | 0.915662 | 0.03 | 0.30773 | Non-Toxin | 100.00% (1/1) | Probable host range protein 2 |
| 564 | A*30:02 | 23 | 31 | 9 | IYEAVIHLY | 0.725814 | 0.04 | 0.25817 | Non-Toxin | 100.00% (1/1) | Probable host range protein 2 |
| 565 | A*30:02 | 11 | 20 | 10 | KVNKVDYTLY | 0.748529 | 0.04 | -0.1345 | Non-Toxin | 100.00% (1/1) | Probable host range protein 2 |
| 566 | A*68:02 | 2 | 10 | 9 | TVFANNYAV | 0.860745 | 0.04 | 0.08472 | Non-Toxin | 100.00% (1/1) | Probable host range protein 2 |
| 567 | B*51:01 | 52 | 59 | 8 | YPYISLNM | 0.787288 | 0.04 | -0.04428 | Non-Toxin | 100.00% (1/1) | Probable host range protein 2 |
| 568 | A*31:01 | 16 | 24 | 9 | ALRNLQLHR | 0.844032 | 0.05 | -0.10001 | Non-Toxin | 100.00% (1/1) | Probable host range protein 2 |
| 569 | A*02:03 | 29 | 37 | 9 | HLYNKKTEI | 0.832788 | 0.05 | -0.32945 | Non-Toxin | 100.00% (1/1) | Probable host range protein 2 |
| 570 | B*35:01 | 21 | 29 | 9 | IPYRDYESM | 0.886212 | 0.05 | 0.05684 | Non-Toxin | 100.00% (1/1) | Probable host range protein 2 |
| 571 | A*24:02 | 39 | 48 | 10 | IYSDDENELF | 0.833531 | 0.05 | 0.15965 | Non-Toxin | 100.00% (1/1) | Probable host range protein 2 |
| 572 | B*15:01 | 31 | 39 | 9 | KLKIISNDY | 0.830264 | 0.05 | 0.04529 | Non-Toxin | 100.00% (1/1) | Probable host range protein 2 |
| 573 | A*02:03 | 22 | 30 | 9 | VIYEAVIHL | 0.834743 | 0.05 | 0.30773 | Non-Toxin | 100.00% (1/1) | Probable host range protein 2 |
| 574 | A*01:01 | 40 | 48 | 9 | YSDDENELF | 0.849408 | 0.05 | 0.19895 | Non-Toxin | 100.00% (1/1) | Probable host range protein 2 |
| 575 | A*26:01 | 46 | 54 | 9 | ELFKHYYPY | 0.706208 | 0.06 | -0.16058 | Non-Toxin | 100.00% (1/1) | Probable host range protein 2 |
| 576 | A*31:01 | 41 | 49 | 9 | KLKLRFVIR | 0.820976 | 0.06 | 0.19204 | Non-Toxin | 100.00% (1/1) | Probable host range protein 2 |
| 577 | B*44:03 | 42 | 51 | 10 | DDENELFKHY | 0.817449 | 0.07 | 0.05979 | Non-Toxin | 100.00% (1/1) | Probable host range protein 2 |
| 578 | B*35:01 | 18 | 26 | 9 | HPLIPYRDY | 0.790827 | 0.08 | 0.17268 | Non-Toxin | 100.00% (1/1) | Probable host range protein 2 |
| 579 | A*23:01 | 39 | 48 | 10 | IYSDDENELF | 0.712093 | 0.08 | 0.15965 | Non-Toxin | 100.00% (1/1) | Probable host range protein 2 |
| 580 | A*30:02 | 31 | 39 | 9 | KLKIISNDY | 0.650988 | 0.08 | 0.04529 | Non-Toxin | 100.00% (1/1) | Probable host range protein 2 |

| | | | | | | | | | | | |
|---|---|---|---|---|---|---|---|---|---|---|---|
| 581 | B*44:03 | 24 | 31 | 8 | YEAVIHLY | 0.791344 | 0.08 | 0.20493 | Non-Toxin | 100.00% (1/1) | Probable host range protein 2 |
| 582 | A*24:02 | 51 | 59 | 9 | YYPYISLNM | 0.738081 | 0.08 | -0.04659 | Non-Toxin | 100.00% (1/1) | Probable host range protein 2 |
| 583 | B*44:02 | 42 | 51 | 10 | DDENELFKHY | 0.722139 | 0.09 | 0.05979 | Non-Toxin | 100.00% (1/1) | Probable host range protein 2 |
| 584 | A*68:01 | 25 | 33 | 9 | EAVIHLYNK | 0.888796 | 0.09 | 0.16148 | Non-Toxin | 100.00% (1/1) | Probable host range protein 2 |
| 585 | A*23:01 | 23 | 31 | 9 | IYEAVIHLY | 0.685447 | 0.09 | 0.25817 | Non-Toxin | 100.00% (1/1) | Probable host range protein 2 |
| 586 | A*02:06 | 14 | 22 | 9 | KVDYTLYYV | 0.751163 | 0.09 | 0.02556 | Non-Toxin | 100.00% (1/1) | Probable host range protein 2 |
| 587 | A*24:02 | 30 | 38 | 9 | LYNKKTEIL | 0.715892 | 0.09 | -0.2303 | Non-Toxin | 100.00% (1/1) | Probable host range protein 2 |
| 588 | B*51:01 | 21 | 29 | 9 | IPYRDYESM | 0.640568 | 0.1 | 0.05684 | Non-Toxin | 100.00% (1/1) | Probable host range protein 2 |
| 589 | B*53:01 | 21 | 29 | 9 | IPYRDYESM | 0.547291 | 0.1 | 0.05684 | Non-Toxin | 100.00% (1/1) | Probable host range protein 2 |
| 590 | A*24:02 | 23 | 31 | 9 | IYEAVIHLY | 0.705229 | 0.1 | 0.25817 | Non-Toxin | 100.00% (1/1) | Probable host range protein 2 |
| 591 | A*32:01 | 22 | 30 | 9 | VIYEAVIHL | 0.501745 | 0.1 | 0.30773 | Non-Toxin | 100.00% (1/1) | Probable host range protein 2 |
| 592 | A*30:02 | 22 | 31 | 10 | VIYEAVIHLY | 0.616456 | 0.1 | 0.32395 | Non-Toxin | 100.00% (1/1) | Probable host range protein 2 |
| 593 | A*23:01 | 51 | 59 | 9 | YYPYISLNM | 0.639636 | 0.1 | -0.04659 | Non-Toxin | 100.00% (1/1) | Probable host range protein 2 |

**Supplementary Table S3:** High Percentile Ranking HTL epitopes-HLA allele pairs screened from eleven proteins of Monkeypox virus by the "MHC-II Binding Predictions" tool of IEDB. Out of these epitopes, the top scoring epitopes were used to design Multi-epitope vaccine.

| # | allele | start | end | length | method | peptide | Percentile rank | Protein | Toxicity | Conservancy |
|---|---|---|---|---|---|---|---|---|---|---|
| 1 | DPA1*02:01/DPB1*14:01 | 1 | 15 | 15 | NetMHCIIpan | MRFKRGAVLPIPNLV | 0.1 | Poxin-Schlafen | Non-Toxin | 66.67% (6/9) |
| 2 | DPA1*02:01/DPB1*14:01 | 2 | 16 | 15 | NetMHCIIpan | RFKRGAVLPIPNLVK | 0.23 | Poxin-Schlafen | Non-Toxin | 66.67% (6/9) |
| 3 | DRB1*09:01 | 32 | 46 | 15 | Consensus (comb.lib./smm/nn) | YRHKNYYALSGIGYE | 0.37 | Poxin-Schlafen | Non-Toxin | 77.78% (7/9) |
| 4 | DRB4*01:01 | 9 | 23 | 15 | Consensus (comb.lib./smm/nn) | QLRTRIRQQLPSILS | 0.38 | Poxin-Schlafen | Non-Toxin | 77.78% (7/9) |
| 5 | DRB4*01:01 | 8 | 22 | 15 | Consensus (comb.lib./smm/nn) | KQLRTRIRQQLPSIL | 0.52 | Poxin-Schlafen | Non-Toxin | 77.78% (7/9) |
| 6 | DRB1*09:01 | 31 | 45 | 15 | Consensus (comb.lib./smm/nn) | TYRHKNYYALSGIGY | 0.75 | Poxin-Schlafen | Non-Toxin | 77.78% (7/9) |
| 7 | DRB1*12:01 | 1 | 15 | 15 | Consensus (smm/nn) | IEKYIQKLPVVHFCK | 0.76 | Poxin-Schlafen | Non-Toxin | 66.67% (6/9) |
| 8 | DPA1*01/DPB1*04:01 | 33 | 47 | 15 | Consensus (comb.lib./smm) | IERLAEEFFNRSELQ | 0.81 | Poxin-Schlafen | Non-Toxin | 88.89% (8/9) |
| 9 | DPA1*01/DPB1*04:01 | 32 | 46 | 15 | Consensus (comb.lib./smm) | SIERLAEEFFNRSEL | 0.81 | Poxin-Schlafen | Non-Toxin | 88.89% (8/9) |
| 10 | DPA1*01/DPB1*04:01 | 34 | 48 | 15 | Consensus (comb.lib./smm) | ERLAEEFFNRSELQA | 0.85 | Poxin-Schlafen | Non-Toxin | 88.89% (8/9) |
| 11 | DPA1*01/DPB1*04:01 | 31 | 45 | 15 | Consensus (comb.lib./smm) | ESIERLAEEFFNRSE | 0.85 | Poxin-Schlafen | Non-Toxin | 88.89% (8/9) |
| 12 | DPA1*01/DPB1*04:01 | 30 | 44 | 15 | Consensus (comb.lib./smm) | TESIERLAEEFFNRS | 0.85 | Poxin-Schlafen | Non-Toxin | 88.89% (8/9) |
| 13 | DQA1*05:01/DQB1*02:01 | 27 | 41 | 15 | Consensus (comb.lib./smm/nn) | VKDTESIERLAEEFF | 0.95 | Poxin-Schlafen | Non-Toxin | 66.67% (6/9) |
| 14 | DQA1*05:01/DQB1*02:01 | 28 | 42 | 15 | Consensus (comb.lib./smm/nn) | KDTESIERLAEEFFN | 0.99 | Poxin-Schlafen | Non-Toxin | 66.67% (6/9) |
| 15 | DPA1*02:01/DPB1*05:01 | 31 | 45 | 15 | Consensus (comb.lib./smm/nn) | ESIERLAEEFFNRSE | 1.1 | Poxin-Schlafen | Non-Toxin | 88.89% (8/9) |
| 16 | DPA1*02:01/DPB1*05:01 | 33 | 47 | 15 | Consensus (comb.lib./smm/nn) | IERLAEEFFNRSELQ | 1.1 | Poxin-Schlafen | Non-Toxin | 88.89% (8/9) |
| 17 | DPA1*02:01/DPB1*05:01 | 32 | 46 | 15 | Consensus (comb.lib./smm/nn) | SIERLAEEFFNRSEL | 1.1 | Poxin-Schlafen | Non-Toxin | 88.89% (8/9) |
| 18 | DRB5*01:01 | 35 | 49 | 15 | Consensus (smm/nn/sturniolo) | SVYNKKYNIVKNKYM | 1.1 | Poxin-Schlafen | Non-Toxin | 66.67% (6/9) |
| 19 | DPA1*02:01/DPB1*05:01 | 30 | 44 | 15 | Consensus (comb.lib./smm/nn) | TESIERLAEEFFNRS | 1.1 | Poxin-Schlafen | Non-Toxin | 88.89% (8/9) |
| 20 | DPA1*02:01/DPB1*05:01 | 29 | 43 | 15 | Consensus (comb.lib./smm/nn) | DTESIERLAEEFFNR | 1.2 | Poxin-Schlafen | Non-Toxin | 88.89% (8/9) |
| 21 | DQA1*05:01/DQB1*02:01 | 29 | 43 | 15 | Consensus (comb.lib./smm/nn) | DTESIERLAEEFFNR | 1.2 | Poxin-Schlafen | Non-Toxin | 88.89% (8/9) |
| 22 | DPA1*02:01/DPB1*14:01 | 3 | 17 | 15 | NetMHCIIpan | FKRGAVLPIPNLVKV | 1.2 | Poxin-Schlafen | Non-Toxin | 66.67% (6/9) |
| 23 | DRB1*08:02 | 2 | 16 | 15 | Consensus (smm/nn/sturniolo) | EKYIQKLPVVHFCKK | 1.3 | Poxin-Schlafen | Non-Toxin | 66.67% (6/9) |
| 24 | DRB1*08:02 | 1 | 15 | 15 | Consensus (smm/nn/sturniolo) | IEKYIQKLPVVHFCK | 1.3 | Poxin-Schlafen | Non-Toxin | 66.67% (6/9) |
| 25 | DPA1*02:01/DPB1*14:01 | 28 | 42 | 15 | NetMHCIIpan | VRKYSVVSVYNKKYN | 1.3 | Poxin-Schlafen | Non-Toxin | 66.67% (6/9) |
| 26 | DRB1*12:01 | 2 | 16 | 15 | Consensus (smm/nn) | EKYIQKLPVVHFCKK | 1.36 | Poxin-Schlafen | Non-Toxin | 66.67% (6/9) |
| 27 | DRB4*01:01 | 7 | 21 | 15 | Consensus (comb.lib./smm/nn) | AKQLRTRIRQQLPSI | 1.4 | Poxin-Schlafen | Non-Toxin | 77.78% (7/9) |
| 28 | DRB4*01:01 | 10 | 24 | 15 | Consensus (comb.lib./smm/nn) | LRTRIRQQLPSILSS | 1.4 | Poxin-Schlafen | Non-Toxin | 77.78% (7/9) |
| 29 | DRB4*01:01 | 11 | 25 | 15 | Consensus (comb.lib./smm/nn) | RTRIRQQLPSILSSF | 1.4 | Poxin-Schlafen | Non-Toxin | 77.78% (7/9) |
| 30 | DQA1*05:01/DQB1*02:01 | 30 | 44 | 15 | Consensus (comb.lib./smm/nn) | TESIERLAEEFFNRS | 1.4 | Poxin-Schlafen | Non-Toxin | 88.89% (8/9) |
| **#** | **allele** | **start** | **end** | **length** | **method** | **peptide** | **percentile _rank** | **Protein** | **Toxicity** | **Conservancy** |
| 31 | DRB1*04:05 | 196 | 210 | 15 | Consensus (smm/nn/sturniolo) | LSKFRTLLSSSNHEG | 0.05 | Cell surface-binding protein | Non-Toxin | 100.00% (7/7) |
| 32 | DRB1*04:05 | 195 | 209 | 15 | Consensus (smm/nn/sturniolo) | QLSKFRTLLSSSNHE | 0.05 | Cell surface-binding protein | Non-Toxin | 100.00% (7/7) |

| # | allele | start | end | length | method | peptide | percentile _rank | Protein | Toxicity | Conservancy |
|---|---|---|---|---|---|---|---|---|---|---|
| 33 | DRB1*04:05 | 194 | 208 | 15 | Consensus (smm/nn/sturniolo) | DQLSKFRTLLSSSNH | 0.06 | Cell surface-binding protein | Non-Toxin | 100.00% (7/7) |
| 34 | DPA1*03:01/DPB1*04:02 | 283 | 297 | 15 | Consensus (comb.lib./smm/nn) | FVFILTAILFLMSQR | 0.06 | Cell surface-binding protein | Non-Toxin | 100.00% (7/7) |
| 35 | DPA1*03:01/DPB1*04:02 | 281 | 295 | 15 | Consensus (comb.lib./smm/nn) | IVFVFILTAILFLMS | 0.06 | Cell surface-binding protein | Non-Toxin | 100.00% (7/7) |
| 36 | DPA1*03:01/DPB1*04:02 | 282 | 296 | 15 | Consensus (comb.lib./smm/nn) | VFVFILTAILFLMSQ | 0.06 | Cell surface-binding protein | Non-Toxin | 100.00% (7/7) |
| 37 | DRB1*04:05 | 197 | 211 | 15 | Consensus (smm/nn/sturniolo) | SKFRTLLSSSNHEGK | 0.07 | Cell surface-binding protein | Non-Toxin | 100.00% (7/7) |
| 38 | DPA1*03:01/DPB1*04:02 | 280 | 294 | 15 | Consensus (comb.lib./smm/nn) | AIVFVFILTAILFLM | 0.08 | Cell surface-binding protein | Non-Toxin | 100.00% (7/7) |
| 39 | DPA1*03:01/DPB1*04:02 | 279 | 293 | 15 | Consensus (comb.lib./smm/nn) | IAIVFVFILTAILFL | 0.09 | Cell surface-binding protein | Non-Toxin | 100.00% (7/7) |
| 40 | DRB1*07:01 | 279 | 293 | 15 | Consensus (comb.lib./smm/nn) | IAIVFVFILTAILFL | 0.12 | Cell surface-binding protein | Non-Toxin | 100.00% (7/7) |
| 41 | DRB1*04:05 | 280 | 294 | 15 | Consensus (smm/nn/sturniolo) | AIVFVFILTAILFLM | 0.13 | Cell surface-binding protein | Non-Toxin | 100.00% (7/7) |
| 42 | DRB1*04:05 | 279 | 293 | 15 | Consensus (smm/nn/sturniolo) | IAIVFVFILTAILFL | 0.13 | Cell surface-binding protein | Non-Toxin | 100.00% (7/7) |
| 43 | DRB1*04:05 | 281 | 295 | 15 | Consensus (smm/nn/sturniolo) | IVFVFILTAILFLMS | 0.13 | Cell surface-binding protein | Non-Toxin | 100.00% (7/7) |
| 44 | DRB1*04:05 | 277 | 291 | 15 | Consensus (smm/nn/sturniolo) | AIIAIVFVFILTAIL | 0.16 | Cell surface-binding protein | Non-Toxin | 100.00% (7/7) |
| 45 | DRB1*04:05 | 278 | 292 | 15 | Consensus (smm/nn/sturniolo) | IIAIVFVFILTAILF | 0.16 | Cell surface-binding protein | Non-Toxin | 100.00% (7/7) |
| 46 | DRB1*04:05 | 193 | 207 | 15 | Consensus (smm/nn/sturniolo) | SDQLSKFRTLLSSSN | 0.18 | Cell surface-binding protein | Non-Toxin | 100.00% (7/7) |
| **#** | **allele** | **start** | **end** | **length** | **method** | **peptide** | **percentile _rank** | **Protein** | **Toxicity** | **Conservancy** |
| 47 | DRB1*13:02 | 31 | 45 | 15 | Consensus (smm/nn/sturniolo) | TVCIIRNINNITYYI | 0.09 | E3 ubiquitin-protein ligase | Non-Toxin | 52.63% (10/19) |
| 48 | DRB1*13:02 | 32 | 46 | 15 | Consensus (smm/nn/sturniolo) | VCIIRNINNITYYIN | 0.09 | E3 ubiquitin-protein ligase | Non-Toxin | 52.63% (10/19) |
| 49 | DRB1*13:02 | 30 | 44 | 15 | Consensus (smm/nn/sturniolo) | LTVCIIRNINNITYY | 0.1 | E3 ubiquitin-protein ligase | Non-Toxin | 52.63% (10/19) |
| 50 | DRB1*13:02 | 29 | 43 | 15 | Consensus (smm/nn/sturniolo) | RLTVCIIRNINNITY | 0.18 | E3 ubiquitin-protein ligase | Non-Toxin | 52.63% (10/19) |
| 51 | DRB1*07:01 | 226 | 240 | 15 | Consensus (comb.lib./smm/nn) | RTRFRKITMSKFYKL | 0.22 | E3 ubiquitin-protein ligase | Non-Toxin | 73.68% (14/19) |
| 52 | DRB3*02:02 | 31 | 45 | 15 | NetMHCIIpan | TVCIIRNINNITYYI | 0.22 | E3 ubiquitin-protein ligase | Non-Toxin | 52.63% (10/19) |
| 53 | DRB1*15:01 | 123 | 137 | 15 | Consensus (smm/nn/sturniolo) | SILRGLVNWYIANNT | 0.27 | E3 ubiquitin-protein ligase | Non-Toxin | 100.00% (19/19) |
| 54 | DRB1*07:01 | 227 | 241 | 15 | Consensus (comb.lib./smm/nn) | TRFRKITMSKFYKLV | 0.27 | E3 ubiquitin-protein ligase | Non-Toxin | 73.68% (14/19) |
| 55 | DRB3*02:02 | 32 | 46 | 15 | NetMHCIIpan | VCIIRNINNITYYIN | 0.27 | E3 ubiquitin-protein ligase | Non-Toxin | 52.63% (10/19) |
| 56 | DRB1*15:01 | 122 | 136 | 15 | Consensus (smm/nn/sturniolo) | QSILRGLVNWYIANN | 0.28 | E3 ubiquitin-protein ligase | Non-Toxin | 100.00% (19/19) |
| **#** | **allele** | **start** | **end** | **length** | **method** | **peptide** | **percentile _rank** | **Protein** | **Toxicity** | **Conservancy** |
| 57 | DRB1*08:02 | 19 | 33 | 15 | Consensus (smm/nn/sturniolo) | TELIRRVRRYQIAQY | 0.22 | Thymidine kinase | Non-Toxin | 99.84% (606/607) |
| 58 | DRB1*08:02 | 20 | 34 | 15 | Consensus (smm/nn/sturniolo) | ELIRRVRRYQIAQYK | 0.27 | Thymidine kinase | Non-Toxin | 99.84% (606/607) |
| 59 | DRB1*04:05 | 115 | 129 | 15 | Consensus (smm/nn/sturniolo) | RRPFNNILNLIPLSE | 0.41 | Thymidine kinase | Non-Toxin | 99.67% (605/607) |
| 60 | DRB1*04:05 | 116 | 130 | 15 | Consensus (smm/nn/sturniolo) | RPFNNILNLIPLSEM | 0.44 | Thymidine kinase | Non-Toxin | 99.67% (605/607) |
| 61 | DRB1*08:02 | 21 | 35 | 15 | Consensus (smm/nn/sturniolo) | LIRRVRRYQIAQYKC | 0.65 | Thymidine kinase | Non-Toxin | 99.84% (606/607) |
| 62 | DRB1*08:02 | 16 | 30 | 15 | Consensus (smm/nn/sturniolo) | GKSTELIRRVRRYQI | 0.75 | Thymidine kinase | Non-Toxin | 99.84% (606/607) |
| 63 | DRB1*08:02 | 17 | 31 | 15 | Consensus (smm/nn/sturniolo) | KSTELIRRVRRYQIA | 0.75 | Thymidine kinase | Non-Toxin | 99.84% (606/607) |
| 64 | DRB1*08:02 | 18 | 32 | 15 | Consensus (smm/nn/sturniolo) | STELIRRVRRYQIAQ | 0.75 | Thymidine kinase | Non-Toxin | 99.84% (606/607) |
| 65 | DRB1*04:05 | 114 | 128 | 15 | Consensus (smm/nn/sturniolo) | QRRPFNNILNLIPLS | 0.78 | Thymidine kinase | Non-Toxin | 99.67% (605/607) |
| 66 | DRB1*04:05 | 113 | 127 | 15 | Consensus (smm/nn/sturniolo) | FQRRPFNNILNLIPL | 0.79 | Thymidine kinase | Non-Toxin | 99.67% (605/607) |
| 67 | DRB1*08:02 | 22 | 36 | 15 | Consensus (smm/nn/sturniolo) | IRRVRRYQIAQYKCV | 0.94 | Thymidine kinase | Non-Toxin | 99.84% (606/607) |
| 68 | DRB1*11:01 | 16 | 30 | 15 | Consensus (smm/nn/sturniolo) | GKSTELIRRVRRYQI | 1.1 | Thymidine kinase | Non-Toxin | 99.84% (606/607) |
| 69 | DRB1*11:01 | 17 | 31 | 15 | Consensus (smm/nn/sturniolo) | KSTELIRRVRRYQIA | 1.1 | Thymidine kinase | Non-Toxin | 99.84% (606/607) |
| 70 | DRB1*11:01 | 18 | 32 | 15 | Consensus (smm/nn/sturniolo) | STELIRRVRRYQIAQ | 1.1 | Thymidine kinase | Non-Toxin | 99.84% (606/607) |
| 71 | DRB1*11:01 | 19 | 33 | 15 | Consensus (smm/nn/sturniolo) | TELIRRVRRYQIAQY | 1.1 | Thymidine kinase | Non-Toxin | 99.84% (606/607) |
| 72 | DRB1*04:05 | 112 | 126 | 15 | Consensus (smm/nn/sturniolo) | TFQRRPFNNILNLIP | 1.1 | Thymidine kinase | Non-Toxin | 99.67% (605/607) |
| 73 | DRB1*12:01 | 118 | 132 | 15 | Consensus (smm/nn) | FNNILNLIPLSEMVV | 1.11 | Thymidine kinase | Non-Toxin | 99.67% (605/607) |
| 74 | DRB1*15:01 | 20 | 34 | 15 | Consensus (smm/nn/sturniolo) | ELIRRVRRYQIAQYK | 1.3 | Thymidine kinase | Non-Toxin | 99.84% (606/607) |
| 75 | DRB1*15:01 | 18 | 32 | 15 | Consensus (smm/nn/sturniolo) | STELIRRVRRYQIAQ | 1.3 | Thymidine kinase | Non-Toxin | 99.84% (606/607) |
| 76 | DRB1*15:01 | 19 | 33 | 15 | Consensus (smm/nn/sturniolo) | TELIRRVRRYQIAQY | 1.3 | Thymidine kinase | Non-Toxin | 99.84% (606/607) |
| **#** | **allele** | **start** | **end** | **length** | **method** | **peptide** | **percentile _rank** | **Protein** | **Toxicity** | **Conservancy** |
| 77 | HLA-DRB1*04:05 | 111 | 125 | 15 | Consensus (smm/nn/sturniolo) | NEKIIHFLTINENGV | 0.43 | Cu-Zn superoxide dismutase-like protein | Non-Toxin | 100.00% (86/86) |
| 78 | HLA-DRB1*04:05 | 109 | 123 | 15 | Consensus (smm/nn/sturniolo) | YINEKIIHFLTINEN | 0.44 | Cu-Zn superoxide dismutase-like protein | Non-Toxin | 100.00% (86/86) |

| | | | | | | | | | | |
|---|---|---|---|---|---|---|---|---|---|---|
| 79 | HLA-DRB1*04:05 | 110 | 124 | 15 | Consensus (smm/nn/sturniolo) | INEKIIHFLTINENG | 0.45 | Cu-Zn superoxide dismutase-like protein | Non-Toxin | 100.00% (86/86) |
| 80 | HLA-DPA1*01/DPB1*04:01 | 59 | 73 | 15 | Consensus (comb.lib./smm) | EIFIGNIFVNRYGVA | 0.6 | Cu-Zn superoxide dismutase-like protein | Non-Toxin | 100.00% (86/86) |
| 81 | HLA-DPA1*01/DPB1*04:01 | 56 | 70 | 15 | Consensus (comb.lib./smm) | GSPEIFIGNIFVNRY | 0.6 | Cu-Zn superoxide dismutase-like protein | Non-Toxin | 100.00% (86/86) |
| 82 | HLA-DPA1*01/DPB1*04:01 | 55 | 69 | 15 | Consensus (comb.lib./smm) | IGSPEIFIGNIFVNR | 0.6 | Cu-Zn superoxide dismutase-like protein | Non-Toxin | 100.00% (86/86) |
| 83 | HLA-DPA1*01/DPB1*04:01 | 58 | 72 | 15 | Consensus (comb.lib./smm) | PEIFIGNIFVNRYGV | 0.6 | Cu-Zn superoxide dismutase-like protein | Non-Toxin | 100.00% (86/86) |
| 84 | HLA-DPA1*01/DPB1*04:01 | 57 | 71 | 15 | Consensus (comb.lib./smm) | SPEIFIGNIFVNRYG | 0.6 | Cu-Zn superoxide dismutase-like protein | Non-Toxin | 100.00% (86/86) |
| 85 | HLA-DRB1*04:05 | 68 | 82 | 15 | Consensus (smm/nn/sturniolo) | NRYGVAYVYLDTDVN | 0.88 | Cu-Zn superoxide dismutase-like protein | Non-Toxin | 100.00% (86/86) |
| 86 | HLA-DRB1*04:05 | 69 | 83 | 15 | Consensus (smm/nn/sturniolo) | RYGVAYVYLDTDVNI | 1.1 | Cu-Zn superoxide dismutase-like protein | Non-Toxin | 100.00% (86/86) |
| 87 | HLA-DRB1*04:05 | 108 | 122 | 15 | Consensus (smm/nn/sturniolo) | SYINEKIIHFLTINE | 1.1 | Cu-Zn superoxide dismutase-like protein | Non-Toxin | 100.00% (86/86) |
| 88 | HLA-DRB3*01:01 | 104 | 118 | 15 | Consensus (comb.lib./smm/nn) | VIGISYINEKIIHFL | 1.1 | Cu-Zn superoxide dismutase-like protein | Non-Toxin | 100.00% (86/86) |
| 89 | HLA-DRB3*01:01 | 106 | 120 | 15 | Consensus (comb.lib./smm/nn) | GISYINEKIIHFLTI | 1.2 | Cu-Zn superoxide dismutase-like protein | Non-Toxin | 100.00% (86/86) |
| 90 | HLA-DRB3*01:01 | 105 | 119 | 15 | Consensus (comb.lib./smm/nn) | IGISYINEKIIHFLT | 1.2 | Cu-Zn superoxide dismutase-like protein | Non-Toxin | 100.00% (86/86) |
| 91 | HLA-DRB1*11:01 | 36 | 50 | 15 | Consensus (smm/nn/sturniolo) | GTYSLIIHRYGDISR | 1.3 | Cu-Zn superoxide dismutase-like protein | Non-Toxin | 100.00% (86/86) |
| 92 | HLA-DRB1*11:01 | 34 | 48 | 15 | Consensus (smm/nn/sturniolo) | KSGTYSLIIHRYGDI | 1.3 | Cu-Zn superoxide dismutase-like protein | Non-Toxin | 100.00% (86/86) |
| 93 | HLA-DRB1*11:01 | 35 | 49 | 15 | Consensus (smm/nn/sturniolo) | SGTYSLIIHRYGDIS | 1.3 | Cu-Zn superoxide dismutase-like protein | Non-Toxin | 100.00% (86/86) |

| # | allele | start | end | length | method | peptide | percentile _rank | Protein | Toxicity | Conservancy |
|---|---|---|---|---|---|---|---|---|---|---|
| 94 | DRB1*15:01 | 15 | 29 | 15 | Consensus (smm/nn/sturniolo) | AVCLLFIQSYSIYEN | 0.13 | Envelope protein A28 | Non-Toxin | 100.00% (574/574) |
| 95 | DRB1*15:01 | 16 | 30 | 15 | Consensus (smm/nn/sturniolo) | VCLLFIQSYSIYENY | 0.13 | Envelope protein A28 | Non-Toxin | 100.00% (574/574) |
| 96 | DRB1*15:01 | 14 | 28 | 15 | Consensus (smm/nn/sturniolo) | AAVCLLFIQSYSIYE | 0.16 | Envelope protein A28 | Non-Toxin | 100.00% (574/574) |
| 97 | DRB1*01:01 | 4 | 18 | 15 | Consensus (comb.lib./smm/nn) | LSIFFIVVATAAVCL | 0.16 | Envelope protein A28 | Non-Toxin | 100.00% (574/574) |
| 98 | DRB1*01:01 | 5 | 19 | 15 | Consensus (comb.lib./smm/nn) | SIFFIVVATAAVCLL | 0.16 | Envelope protein A28 | Non-Toxin | 100.00% (574/574) |
| 99 | DRB1*01:01 | 6 | 20 | 15 | Consensus (comb.lib./smm/nn) | IFFIVVATAAVCLLF | 0.24 | Envelope protein A28 | Non-Toxin | 100.00% (574/574) |
| 100 | DRB1*01:01 | 2 | 16 | 15 | Consensus (comb.lib./smm/nn) | NSLSIFFIVVATAAV | 0.24 | Envelope protein A28 | Non-Toxin | 100.00% (574/574) |
| 101 | DRB1*01:01 | 3 | 17 | 15 | Consensus (comb.lib./smm/nn) | SLSIFFIVVATAAVC | 0.24 | Envelope protein A28 | Non-Toxin | 100.00% (574/574) |

| # | allele | start | end | length | method | peptide | percentile _rank | Protein | Toxicity | Conservancy |
|---|---|---|---|---|---|---|---|---|---|---|
| 102 | DRB4*01:01 | 755 | 769 | 15 | Consensus (comb.lib./smm/nn) | EDGIIIKKQFIQRGG | 0.02 | DNA-directed RNA polymerase | Non-Toxin | 100.00% (10/10) |
| 103 | DRB4*01:01 | 754 | 768 | 15 | Consensus (comb.lib./smm/nn) | QEDGIIIKKQFIQRG | 0.02 | DNA-directed RNA polymerase | Non-Toxin | 100.00% (10/10) |
| 104 | DRB4*01:01 | 756 | 770 | 15 | Consensus (comb.lib./smm/nn) | DGIIIKKQFIQRGGL | 0.03 | DNA-directed RNA polymerase | Non-Toxin | 100.00% (10/10) |
| 105 | DRB4*01:01 | 757 | 771 | 15 | Consensus (comb.lib./smm/nn) | GIIIKKQFIQRGGLD | 0.03 | DNA-directed RNA polymerase | Non-Toxin | 100.00% (10/10) |
| 106 | DRB4*01:01 | 758 | 772 | 15 | Consensus (comb.lib./smm/nn) | IIIKKQFIQRGGLDI | 0.03 | DNA-directed RNA polymerase | Non-Toxin | 100.00% (10/10) |
| 107 | DPA1*01:03/DPB1*02:01 | 32 | 46 | 15 | Consensus (comb.lib./smm/nn) | HFQYVSYSNFILHRL | 0.05 | DNA-directed RNA polymerase | Non-Toxin | 100.00% (10/10) |
| 108 | DPA1*01:03/DPB1*02:01 | 31 | 45 | 15 | Consensus (comb.lib./smm/nn) | LHFQYVSYSNFILHR | 0.05 | DNA-directed RNA polymerase | Non-Toxin | 100.00% (10/10) |
| 109 | DPA1*01:03/DPB1*02:01 | 30 | 44 | 15 | Consensus (comb.lib./smm/nn) | PLHFQYVSYSNFILH | 0.05 | DNA-directed RNA polymerase | Non-Toxin | 100.00% (10/10) |
| 110 | DPA1*01:03/DPB1*02:01 | 29 | 43 | 15 | Consensus (comb.lib./smm/nn) | RPLHFQYVSYSNFIL | 0.05 | DNA-directed RNA polymerase | Non-Toxin | 100.00% (10/10) |
| 111 | DRB3*01:01 | 510 | 524 | 15 | Consensus (comb.lib./smm/nn) | CEYIRSYYKDDISYF | 0.08 | DNA-directed RNA polymerase | Non-Toxin | 100.00% (10/10) |
| 112 | DRB3*01:01 | 511 | 525 | 15 | Consensus (comb.lib./smm/nn) | EYIRSYYKDDISYFE | 0.08 | DNA-directed RNA polymerase | Non-Toxin | 100.00% (10/10) |
| 113 | DRB3*01:01 | 513 | 527 | 15 | Consensus (comb.lib./smm/nn) | IRSYYKDDISYFETG | 0.08 | DNA-directed RNA polymerase | Non-Toxin | 100.00% (10/10) |
| 114 | DPA1*02:01/DPB1*01:01 | 30 | 44 | 15 | Consensus (comb.lib./smm/nn) | PLHFQYVSYSNFILH | 0.08 | DNA-directed RNA polymerase | Non-Toxin | 100.00% (10/10) |
| 115 | DRB3*01:01 | 514 | 528 | 15 | Consensus (comb.lib./smm/nn) | RSYYKDDISYFETGF | 0.08 | DNA-directed RNA polymerase | Non-Toxin | 100.00% (10/10) |
| 116 | DRB3*01:01 | 512 | 526 | 15 | Consensus (comb.lib./smm/nn) | YIRSYYKDDISYFET | 0.08 | DNA-directed RNA polymerase | Non-Toxin | 100.00% (10/10) |
| 117 | DPA1*02:01/DPB1*01:01 | 31 | 45 | 15 | Consensus (comb.lib./smm/nn) | LHFQYVSYSNFILHR | 0.09 | DNA-directed RNA polymerase | Non-Toxin | 100.00% (10/10) |
| 118 | DPA1*01:03/DPB1*02:01 | 33 | 47 | 15 | Consensus (comb.lib./smm/nn) | FQYVSYSNFILHRLH | 0.11 | DNA-directed RNA polymerase | Non-Toxin | 100.00% (10/10) |
| 119 | DPA1*02:01/DPB1*01:01 | 29 | 43 | 15 | Consensus (comb.lib./smm/nn) | RPLHFQYVSYSNFIL | 0.11 | DNA-directed RNA polymerase | Non-Toxin | 100.00% (10/10) |
| 120 | DQA1*05:01/DQB1*02:01 | 890 | 904 | 15 | Consensus (comb.lib./smm/nn) | KGTVAYIADETELPY | 0.12 | DNA-directed RNA polymerase | Non-Toxin | 100.00% (10/10) |
| 121 | DPA1*02:01/DPB1*01:01 | 32 | 46 | 15 | Consensus (comb.lib./smm/nn) | HFQYVSYSNFILHRL | 0.13 | DNA-directed RNA polymerase | Non-Toxin | 100.00% (10/10) |

| # | allele | start | end | length | method | peptide | percentile _rank | Protein | Toxicity | Conservancy |
|---|---|---|---|---|---|---|---|---|---|---|
| 122 | DRB1*09:01 | 6 | 20 | 15 | Consensus (comb.lib./smm/nn) | DQLVFNSISARALKA | 0.03 | Telomere-binding protein | Non-Toxin | 99.48% (579/582) |
| 123 | DRB1*09:01 | 7 | 21 | 15 | Consensus (comb.lib./smm/nn) | QLVFNSISARALKAY | 0.03 | Telomere-binding protein | Non-Toxin | 99.48% (579/582) |
| 124 | DRB1*09:01 | 5 | 19 | 15 | Consensus (comb.lib./smm/nn) | EDQLVFNSISARALK | 0.06 | Telomere-binding protein | Non-Toxin | 99.48% (579/582) |

| | | | | | | | | | | |
|---|---|---|---|---|---|---|---|---|---|---|
| 125 | DPA1*02:01/DPB1*14:01 | 7 | 21 | 15 | NetMHCIIpan | QLVFNSISARALKAY | 0.08 | Telomere-binding protein | Non-Toxin | 99.48% (579/582) |
| 126 | DPA1*02:01/DPB1*14:01 | 8 | 22 | 15 | NetMHCIIpan | LVFNSISARALKAYF | 0.11 | Telomere-binding protein | Non-Toxin | 99.48% (579/582) |
| 127 | DRB1*09:01 | 4 | 18 | 15 | Consensus (comb.lib./smm/nn) | FEDQLVFNSISARAL | 0.12 | Telomere-binding protein | Non-Toxin | 99.48% (579/582) |
| 128 | DRB1*09:01 | 8 | 22 | 15 | Consensus (comb.lib./smm/nn) | LVFNSISARALKAYF | 0.12 | Telomere-binding protein | Non-Toxin | 99.48% (579/582) |
| 129 | DRB1*09:01 | 9 | 23 | 15 | Consensus (comb.lib./smm/nn) | VFNSISARALKAYFT | 0.14 | Telomere-binding protein | Non-Toxin | 99.48% (579/582) |
| 130 | DPA1*02:01/DPB1*14:01 | 6 | 20 | 15 | NetMHCIIpan | DQLVFNSISARALKA | 0.17 | Telomere-binding protein | Non-Toxin | 99.48% (579/582) |
| 131 | DRB5*01:01 | 7 | 21 | 15 | Consensus (smm/nn/sturniolo) | QLVFNSISARALKAY | 0.21 | Telomere-binding protein | Non-Toxin | 99.48% (579/582) |
| **#** | **allele** | **start** | **end** | **length** | **method** | **peptide** | **percentile _rank** | **Protein** | **Toxicity** | **Conservancy** |
| 132 | DRB1*04:01 | 21 | 35 | 15 | Consensus (smm/nn/sturniolo) | AIVDYKTTKNVLAAI | 0.52 | Profilin | Non-Toxin | 99.31% (577/581) |
| 133 | DRB1*04:01 | 20 | 34 | 15 | Consensus (smm/nn/sturniolo) | AAIVDYKTTKNVLAA | 0.55 | Profilin | Non-Toxin | 99.14% (576/581) |
| 134 | DRB1*04:01 | 19 | 33 | 15 | Consensus (smm/nn/sturniolo) | DAAIVDYKTTKNVLA | 0.55 | Profilin | Non-Toxin | 99.14% (576/581) |
| 135 | DRB1*04:01 | 22 | 36 | 15 | Consensus (smm/nn/sturniolo) | IVDYKTTKNVLAAIP | 0.55 | Profilin | Non-Toxin | 99.66% (579/581) |
| 136 | DRB1*04:01 | 23 | 37 | 15 | Consensus (smm/nn/sturniolo) | VDYKTTKNVLAAIPN | 0.58 | Profilin | Non-Toxin | 99.66% (579/581) |
| 137 | DRB1*04:01 | 92 | 106 | 15 | Consensus (smm/nn/sturniolo) | SPIVIARTHTALIFL | 0.63 | Profilin | Non-Toxin | 99.48% (578/581) |
| 138 | DRB1*04:01 | 91 | 105 | 15 | Consensus (smm/nn/sturniolo) | VSPIVIARTHTALIF | 0.63 | Profilin | Non-Toxin | 99.48% (578/581) |
| **#** | **allele** | **start** | **end** | **length** | **method** | **peptide** | **percentile _rank** | **Protein** | **Toxicity** | **Conservancy** |
| 139 | DPA1*01:03/DPB1*02:01 | 236 | 250 | 15 | Consensus (comb.lib./smm/nn) | DKLFYVYYNLFLLFE | 0.02 | Intermediate transcription factor 3 | Non-Toxin | 100.00% (9/9) |
| 140 | DPA1*01:03/DPB1*02:01 | 235 | 249 | 15 | Consensus (comb.lib./smm/nn) | IDKLFYVYYNLFLLF | 0.02 | Intermediate transcription factor 3 | Non-Toxin | 100.00% (9/9) |
| 141 | DPA1*01:03/DPB1*02:01 | 237 | 251 | 15 | Consensus (comb.lib./smm/nn) | KLFYVYYNLFLLFED | 0.03 | Intermediate transcription factor 3 | Non-Toxin | 100.00% (9/9) |
| 142 | DPA1*01:03/DPB1*02:01 | 238 | 252 | 15 | Consensus (comb.lib./smm/nn) | LFYVYYNLFLLFEDI | 0.03 | Intermediate transcription factor 3 | Non-Toxin | 100.00% (9/9) |
| 143 | DPA1*01:03/DPB1*02:01 | 234 | 248 | 15 | Consensus (comb.lib./smm/nn) | MIDKLFYVYYNLFLL | 0.03 | Intermediate transcription factor 3 | Non-Toxin | 100.00% (9/9) |
| 144 | DPA1*01:03/DPB1*02:01 | 239 | 253 | 15 | Consensus (comb.lib./smm/nn) | FYVYYNLFLLFEDII | 0.05 | Intermediate transcription factor 3 | Non-Toxin | 100.00% (9/9) |
| 145 | DRB1*15:01 | 320 | 334 | 15 | Consensus (smm/nn/sturniolo) | DSKIKFFKGKKLNIV | 0.06 | Intermediate transcription factor 3 | Non-Toxin | 100.00% (9/9) |
| 146 | DRB1*15:01 | 319 | 333 | 15 | Consensus (smm/nn/sturniolo) | EDSKIKFFKGKKLNI | 0.07 | Intermediate transcription factor 3 | Non-Toxin | 100.00% (9/9) |
| 147 | DRB1*15:01 | 321 | 335 | 15 | Consensus (smm/nn/sturniolo) | SKIKFFKGKKLNIVA | 0.07 | Intermediate transcription factor 3 | Non-Toxin | 100.00% (9/9) |
| 148 | DQA1*01:01/DQB1*05:01 | 241 | 255 | 15 | Consensus (comb.lib./smm/nn) | VYYNLFLLFEDIIQN | 0.07 | Intermediate transcription factor 3 | Non-Toxin | 100.00% (9/9) |
| 149 | DQA1*01:01/DQB1*05:01 | 240 | 254 | 15 | Consensus (comb.lib./smm/nn) | YVYYNLFLLFEDIIQ | 0.07 | Intermediate transcription factor 3 | Non-Toxin | 100.00% (9/9) |
| 150 | DQA1*01:01/DQB1*05:01 | 242 | 256 | 15 | Consensus (comb.lib./smm/nn) | YYNLFLLFEDIIQNE | 0.07 | Intermediate transcription factor 3 | Non-Toxin | 11.11% (1/9) |
| 151 | DQA1*01:01/DQB1*05:01 | 243 | 257 | 15 | Consensus (comb.lib./smm/nn) | YNLFLLFEDIIQNEY | 0.09 | Intermediate transcription factor 3 | Non-Toxin | 11.11% (1/9) |
| 152 | DQA1*01:01/DQB1*05:01 | 244 | 258 | 15 | Consensus (comb.lib./smm/nn) | NLFLLFEDIIQNEYF | 0.11 | Intermediate transcription factor 3 | Non-Toxin | 11.11% (1/9) |
| 153 | DPA1*01/DPB1*04:01 | 236 | 250 | 15 | Consensus (comb.lib./smm) | DKLFYVYYNLFLLFE | 0.12 | Intermediate transcription factor 3 | Non-Toxin | 100.00% (9/9) |
| 154 | DPA1*01/DPB1*04:01 | 235 | 249 | 15 | Consensus (comb.lib./smm) | IDKLFYVYYNLFLLF | 0.12 | Intermediate transcription factor 3 | Non-Toxin | 100.00% (9/9) |
| 155 | DPA1*01/DPB1*04:01 | 237 | 251 | 15 | Consensus (comb.lib./smm) | KLFYVYYNLFLLFED | 0.12 | Intermediate transcription factor 3 | Non-Toxin | 100.00% (9/9) |
| 156 | DPA1*01/DPB1*04:01 | 238 | 252 | 15 | Consensus (comb.lib./smm) | LFYVYYNLFLLFEDI | 0.12 | Intermediate transcription factor 3 | Non-Toxin | 100.00% (9/9) |
| 157 | DRB1*15:01 | 318 | 332 | 15 | Consensus (smm/nn/sturniolo) | DEDSKIKFFKGKKLN | 0.13 | Intermediate transcription factor 3 | Non-Toxin | 100.00% (9/9) |
| 158 | DRB3*01:01 | 208 | 222 | 15 | Consensus (comb.lib./smm/nn) | TCVKIFKDSTMHIIL | 0.13 | Intermediate transcription factor 3 | Non-Toxin | 100.00% (9/9) |
| 159 | DRB3*01:01 | 209 | 223 | 15 | Consensus (comb.lib./smm/nn) | CVKIFKDSTMHIILS | 0.15 | Intermediate transcription factor 3 | Non-Toxin | 100.00% (9/9) |
| 160 | DPA1*01/DPB1*04:01 | 239 | 253 | 15 | Consensus (comb.lib./smm) | FYVYYNLFLLFEDII | 0.15 | Intermediate transcription factor 3 | Non-Toxin | 100.00% (9/9) |
| 161 | DRB3*01:01 | 210 | 224 | 15 | Consensus (comb.lib./smm/nn) | VKIFKDSTMHIILSK | 0.15 | Intermediate transcription factor 3 | Non-Toxin | 100.00% (9/9) |
| 162 | DPA1*01:03/DPB1*02:01 | 240 | 254 | 15 | Consensus (comb.lib./smm/nn) | YVYYNLFLLFEDIIQ | 0.18 | Intermediate transcription factor 3 | Non-Toxin | 100.00% (9/9) |
| **#** | **allele** | **start** | **end** | **length** | **method** | **peptide** | **percentile _rank** | **Protein** | **Toxicity** | **Conservancy** |
| 163 | DRB3*02:02 | 60 | 74 | 15 | NetMHCIIpan | GLTVFANNYAVKVNK | 0.01 | Probable host range protein 2 | Non-Toxin | 100.00% (1/1) |
| 164 | DRB3*02:02 | 61 | 75 | 15 | NetMHCIIpan | LTVFANNYAVKVNKV | 0.01 | Probable host range protein 2 | Non-Toxin | 100.00% (1/1) |
| 165 | DRB3*02:02 | 62 | 76 | 15 | NetMHCIIpan | TVFANNYAVKVNKVD | 0.05 | Probable host range protein 2 | Non-Toxin | 100.00% (1/1) |
| 166 | DRB3*02:02 | 59 | 73 | 15 | NetMHCIIpan | KGLTVFANNYAVKVN | 0.06 | Probable host range protein 2 | Non-Toxin | 100.00% (1/1) |
| 167 | DRB3*02:02 | 58 | 72 | 15 | NetMHCIIpan | VKGLTVFANNYAVKV | 0.17 | Probable host range protein 2 | Non-Toxin | 100.00% (1/1) |
| 168 | DRB3*02:02 | 63 | 77 | 15 | NetMHCIIpan | VFANNYAVKVNKVDY | 0.2 | Probable host range protein 2 | Non-Toxin | 100.00% (1/1) |

**Supplementary Table S4:** Construct of CTL-MEV and HTL-MEV.

| S.No. | Vaccine | Vaccine constructs |
|---|---|---|
| 1 | CTL-MEV construct (Comprising of identified CTL Epitopes from Monkeypox virus proteins): | GIGDPVTCLKSGAICHPVFCPRRYKQIGTCGLPGTKCCKKPEAAAKRYIGALLPMFGGGGSRYIGALLPMFGGGGSMSAPFDSVFGGGGSMSAPFDSVFYGGGGSMSAPFDSVFGGGGSMSAPFDSVFYGGGGSKYIEGNKTFGGGGSMSAPFDSVFYGGGGSIIRNINNITYGGGGSDIIKILDKYGGGGSAQYKCVTIKYGGGGSEIIGGNDMYGGGGSRVRRYQIAQYGGGGSFVNRYGVAYGGGGSSYINEKIIHFGGGGSFSDVINIDIYGGGGSNATHAAFEYGGGGSKQKWRCVVYGGGGSLLFIQSYSIYGGGGSTMRQCIDFTFGGGGSKSYDALATFGGGGSYTPDQLKGFYGGGGSFPAEFRDGYGGGGSKSYDALATFGGGGSFARDNQISFGGGGSKYHPSQYLYFGGGGSSEMDQRLGYGGGGSTQFNFNGHTYGGGGSAEFEDQLVFGGGGSAEFEDQLVFGGGGSYAMELLTGYGGGGSYAMELLTGYGGGGSYAMELLTGYGGGGSVYYNLFLLFGGGGSRYARTIFNFGGGGSRYARTIFNFGGGGSKVNKVDYTLYGGGGSKVNKVDYTLYYGGGGSKLKLRFVIREAAAKGIINTLQKYYCRVRGGRCAVLSCLPKEEQIGKCSTRGRKCCRRKKHHHHHH |
| 2 | HTL-MEV construct (Comprising of identified HTL Epitopes from Monkeypox virus proteins): | GIGDPVTCLKSGAICHPVFCPRRYKQIGTCGLPGTKCCKKPEAAAKMRFKRGAVLPIPNLVGGGGSRFKRGAVLPIPNLVKGGGGSYRHKNYYALSGIGYEGGGGSLSKFRTLLSSSNHEGGGGGSQLSKFRTLLSSSNHEGGGGSDQLSKFRTLLSSSNHGGGGSFVFILTAILFLMSQRGGGGSIVFVFILTAILFLMSGGGGSTVCIIRNINNITYYIGGGGSVCIIRNINNITYYINGGGGSLTVCIIRNINNITYYGGGGSTELIRRVRRYQIAQYGGGGSELIRRVRRYQIAQYKGGGGSRRPFNNILNLIPLSEGGGGSNEKIIHFLTINENGVGGGGSYINEKIIHFLTINENGGGGSINEKIIHFLTINENGGGGGSAVCLLFIQSYSIYENGGGGSVCLLFIQSYSIYENYGGGGSAAVCLLFIQSYSIYEGGGGSEDGIIIKKQFIQRGGGGGGSQEDGIIIKKQFIQRGGGGGSDGIIIKKQFIQRGGLGGGGSGIIIKKQFIQRGGLDGGGGSIIIKKQFIQRGGLDIGGGGSHFQYVSYSNFILHRLGGGGSLHFQYVSYSNFILHRGGGGSDQLVFNSISARALKAGGGGSQLVFNSISARALKAYGGGGSEDQLVFNSISARALKGGGGSAIVDYKTTKNVLAAIGGGGSAAIVDYKTTKNVLAAGGGGSDKLFYVYYNLFLLFEGGGGSIDKLFYVYYNLFLLFGGGGSKLFYVYYNLFLLFEDGGGGSLFYVYYNLFLLFEDIGGGGSMIDKLFYVYYNLFLLGGGGSGLTVFANNYAVKVNKGGGGSLTVFANNYAVKVNKVEAAAKGIINTLQKYYCRVRGGRCAVLSCLPKEEQIGKCSTRGRKCCRRKKHHHHHH |

**Supplementary Table S5:** HLA alleles covered by the CTL and HTL epitopes.

| S.No. | HLA alleles | Class of HLA alleles | | S.No. | HLA alleles | Class of HLA alleles |
|---|---|---|---|---|---|---|
| 1 | A*01:01 | HLA Class I alleles | | 13 | DPA1*01:03/DPB1*02:01 | HLA Class II alleles |
| 2 | A*23:01 | | | 14 | DPA1*02:01/DPB1*14:01 | |
| 3 | A*24:02 | | | 15 | DPA1*03:01/DPB1*04:02 | |
| 4 | A*26:01 | | | 16 | DRB1*04:01 | |
| 5 | A*30:02 | | | 17 | DRB1*04:05 | |
| 6 | A*31:01 | | | 18 | DRB1*08:02 | |
| 7 | A*32:01 | | | 19 | DRB1*09:01 | |
| 8 | B*15:01 | | | 20 | DRB1*13:02 | |
| 9 | B*35:01 | | | 21 | DRB1*15:01 | |
| 10 | B*40:01 | | | 22 | DRB3*02:02 | |
| 11 | B*44:03 | | | 23 | DRB4*01:01 | |
| 12 | B*58:01 | | | 24 | DRB1*04:05 | |

**Supplementary Table S6:** Empirical physicochemical properties of Multi-Patch Vaccine candidates.

| MPVs | Length | Molecular Weight | Theoretical pI | Expected half-life | Aliphatic index | Grand average of Hydropathicity (GRAVY) | Instability index score |
|---|---|---|---|---|---|---|---|
| CTL-MEV | 661 aa | 67.58 kDa | 9.16 | E.Coli: >10 Hr<br>Yeast: >20 Hr<br>Mammalian cell: 30 Hr | 54.02 | -0.227 | 52.65 |
| HTL-MEV | 877 aa | 92.18 kDa | 9.70 | E.Coli: >10 Hr<br>Yeast: >20 Hr<br>Mammalian cell: 30 Hr | 88.93 | 0.008 | 46.95 |

**Supplementary Table S7:** INF-γ inducing POSITIVE epitopes, screened from the CTL MEV.

| # | Sequence | Method | Result | Score | | # | Sequence | Method | Result | Score |
|---|---|---|---|---|---|---|---|---|---|---|
| 1 | RRYKQIGTCGLPGTK | MERCI | POSITIVE | 1 | | 38 | GGGSSEMDQRLGYGG | MERCI | POSITIVE | 1 |
| 2 | RYKQIGTCGLPGTKC | MERCI | POSITIVE | 1 | | 39 | TYGGGGSAEFEDQLV | MERCI | POSITIVE | 2 |
| 3 | YKQIGTCGLPGTKCC | MERCI | POSITIVE | 1 | | 40 | YGGGGSAEFEDQLVF | MERCI | POSITIVE | 3 |
| 4 | KQIGTCGLPGTKCCK | MERCI | POSITIVE | 1 | | 41 | GGGGSAEFEDQLVFG | MERCI | POSITIVE | 3 |
| 5 | QIGTCGLPGTKCCKK | MERCI | POSITIVE | 1 | | 42 | GGGSAEFEDQLVFGG | MERCI | POSITIVE | 3 |
| 6 | GGGGSKYIEGNKTFG | MERCI | POSITIVE | 1 | | 43 | GGSAEFEDQLVFGGG | MERCI | POSITIVE | 3 |
| 7 | GGGSKYIEGNKTFGG | MERCI | POSITIVE | 1 | | 44 | GSAEFEDQLVFGGGG | MERCI | POSITIVE | 1 |
| 8 | GGSKYIEGNKTFGGG | MERCI | POSITIVE | 1 | | 45 | SAEFEDQLVFGGGGS | MERCI | POSITIVE | 1 |
| 9 | GSKYIEGNKTFGGGG | MERCI | POSITIVE | 1 | | 46 | AEFEDQLVFGGGGSA | MERCI | POSITIVE | 1 |
| 10 | SKYIEGNKTFGGGGS | MERCI | POSITIVE | 1 | | 47 | EFEDQLVFGGGGSAE | MERCI | POSITIVE | 1 |
| 11 | KYIEGNKTFGGGGSM | MERCI | POSITIVE | 1 | | 48 | FEDQLVFGGGGSAEF | MERCI | POSITIVE | 1 |
| 12 | YIEGNKTFGGGGSMS | MERCI | POSITIVE | 1 | | 49 | EDQLVFGGGGSAEFE | MERCI | POSITIVE | 3 |
| 13 | IEGNKTFGGGGSMSA | MERCI | POSITIVE | 1 | | 50 | DQLVFGGGGSAEFED | MERCI | POSITIVE | 2 |
| 14 | SIIRNINNITYGGGG | MERCI | POSITIVE | 6 | | 51 | QLVFGGGGSAEFEDQ | MERCI | POSITIVE | 2 |
| 15 | IIRNINNITYGGGGS | MERCI | POSITIVE | 6 | | 52 | LVFGGGGSAEFEDQL | MERCI | POSITIVE | 2 |
| 16 | IRNINNITYGGGGSD | MERCI | POSITIVE | 6 | | 53 | VFGGGGSAEFEDQLV | MERCI | POSITIVE | 2 |
| 17 | RNINNITYGGGGSDI | MERCI | POSITIVE | 6 | | 54 | FGGGGSAEFEDQLVF | MERCI | POSITIVE | 3 |
| 18 | LKGFYGGGGSFPAEF | MERCI | POSITIVE | 1 | | 55 | GGGGSAEFEDQLVFG | MERCI | POSITIVE | 3 |
| 19 | KGFYGGGGSFPAEFR | MERCI | POSITIVE | 2 | | 56 | GGGSAEFEDQLVFGG | MERCI | POSITIVE | 3 |
| 20 | GFYGGGGSFPAEFRD | MERCI | POSITIVE | 2 | | 57 | GGSAEFEDQLVFGGG | MERCI | POSITIVE | 3 |
| 21 | FYGGGGSFPAEFRDG | MERCI | POSITIVE | 2 | | 58 | GSAEFEDQLVFGGGG | MERCI | POSITIVE | 1 |
| 22 | YGGGGSFPAEFRDGY | MERCI | POSITIVE | 2 | | 59 | SAEFEDQLVFGGGGS | MERCI | POSITIVE | 1 |
| 23 | GGGGSFPAEFRDGYG | MERCI | POSITIVE | 2 | | 60 | AEFEDQLVFGGGGSY | MERCI | POSITIVE | 1 |
| 24 | GGGSFPAEFRDGYGG | MERCI | POSITIVE | 2 | | 61 | EFEDQLVFGGGGSYA | MERCI | POSITIVE | 1 |
| 25 | GGSFPAEFRDGYGGG | MERCI | POSITIVE | 2 | | 62 | FEDQLVFGGGGSYAM | MERCI | POSITIVE | 1 |
| 26 | GSFPAEFRDGYGGGG | MERCI | POSITIVE | 2 | | 63 | EDQLVFGGGGSYAME | MERCI | POSITIVE | 1 |
| 27 | SFPAEFRDGYGGGGS | MERCI | POSITIVE | 2 | | 64 | TGYGGGGSVYYNLFL | MERCI | POSITIVE | 1 |
| 28 | QISFGGGGSKYHPSQ | MERCI | POSITIVE | 2 | | 65 | GYGGGGSVYYNLFLL | MERCI | POSITIVE | 1 |
| 29 | HPSQYLYFGGGGSSE | MERCI | POSITIVE | 1 | | 66 | YGGGGSVYYNLFLLF | MERCI | POSITIVE | 1 |

| | | | | | | | | | | |
|---|---|---|---|---|---|---|---|---|---|---|
| 30 | PSQYLYFGGGGSSEM | MERCI | POSITIVE | 1 | | 67 | GGGGSVYYNLFLLFG | MERCI | POSITIVE | 1 |
| 31 | SQYLYFGGGGSSEMD | MERCI | POSITIVE | 1 | | 68 | GGGSVYYNLFLLFGG | MERCI | POSITIVE | 1 |
| 32 | QYLYFGGGGSSEMDQ | MERCI | POSITIVE | 1 | | 69 | GGSVYYNLFLLFGGG | MERCI | POSITIVE | 1 |
| 33 | YLYFGGGGSSEMDQR | MERCI | POSITIVE | 1 | | 70 | GSVYYNLFLLFGGGG | MERCI | POSITIVE | 1 |
| 34 | LYFGGGGSSEMDQRL | MERCI | POSITIVE | 1 | | 71 | SVYYNLFLLFGGGGS | MERCI | POSITIVE | 1 |
| 35 | YFGGGGSSEMDQRLG | MERCI | POSITIVE | 1 | | 72 | VYYNLFLLFGGGGSR | MERCI | POSITIVE | 1 |
| 36 | FGGGGSSEMDQRLGY | MERCI | POSITIVE | 1 | | 73 | STRGRKCCRRKKHHH | MERCI | POSITIVE | 1 |
| 37 | GGGGSSEMDQRLGYG | MERCI | POSITIVE | 1 | | 74 | TRGRKCCRRKKHHHH | MERCI | POSITIVE | 1 |

**Supplementary Table S8:** INF-γ inducing POSITIVE epitopes, screened from the HTL MPVs.

| Serial No. | Sequence | Method | Result | Score | Serial No. | Sequence | Method | Result | Score |
|---|---|---|---|---|---|---|---|---|---|
| 1 | RRYKQIGTCGLPGTK | MERCI | POSITIVE | 1 | 48 | GSDKLFYVYYNLFLL | MERCI | POSITIVE | 1 |
| 2 | RYKQIGTCGLPGTKC | MERCI | POSITIVE | 1 | 49 | SDKLFYVYYNLFLLF | MERCI | POSITIVE | 1 |
| 3 | YKQIGTCGLPGTKCC | MERCI | POSITIVE | 1 | 50 | DKLFYVYYNLFLLFE | MERCI | POSITIVE | 1 |
| 4 | KQIGTCGLPGTKCCK | MERCI | POSITIVE | 1 | 51 | KLFYVYYNLFLLFEG | MERCI | POSITIVE | 1 |
| 5 | QIGTCGLPGTKCCKK | MERCI | POSITIVE | 1 | 52 | LFYVYYNLFLLFEGG | MERCI | POSITIVE | 1 |
| 6 | CKKPEAAAKMRFKRG | MERCI | POSITIVE | 1 | 53 | FYVYYNLFLLFEGGG | MERCI | POSITIVE | 1 |
| 7 | YYGGGGSTELIRRVR | MERCI | POSITIVE | 1 | 54 | YVYYNLFLLFEGGGG | MERCI | POSITIVE | 1 |
| 8 | YGGGGSTELIRRVRR | MERCI | POSITIVE | 1 | 55 | VYYNLFLLFEGGGGS | MERCI | POSITIVE | 1 |
| 9 | GGGGSTELIRRVRRY | MERCI | POSITIVE | 1 | 56 | GSIDKLFYVYYNLFL | MERCI | POSITIVE | 1 |
| 10 | GGGSTELIRRVRRYQ | MERCI | POSITIVE | 1 | 57 | SIDKLFYVYYNLFLL | MERCI | POSITIVE | 1 |
| 11 | LNLIPLSEGGGGSNE | MERCI | POSITIVE | 1 | 58 | IDKLFYVYYNLFLLF | MERCI | POSITIVE | 1 |
| 12 | NLIPLSEGGGGSNEK | MERCI | POSITIVE | 1 | 59 | DKLFYVYYNLFLLFG | MERCI | POSITIVE | 1 |
| 13 | LIPLSEGGGGSNEKI | MERCI | POSITIVE | 1 | 60 | KLFYVYYNLFLLFGG | MERCI | POSITIVE | 1 |
| 14 | IPLSEGGGGSNEKII | MERCI | POSITIVE | 1 | 61 | LFYVYYNLFLLFGGG | MERCI | POSITIVE | 1 |
| 15 | PLSEGGGGSNEKIIH | MERCI | POSITIVE | 1 | 62 | FYVYYNLFLLFGGGG | MERCI | POSITIVE | 1 |
| 16 | LSEGGGGSNEKIIHF | MERCI | POSITIVE | 1 | 63 | YVYYNLFLLFGGGGS | MERCI | POSITIVE | 1 |
| 17 | KQFIQRGGGGGGSQE | MERCI | POSITIVE | 1 | 64 | VYYNLFLLFGGGGSK | MERCI | POSITIVE | 1 |
| 18 | QFIQRGGGGGGSQED | MERCI | POSITIVE | 1 | 65 | GGGSKLFYVYYNLFL | MERCI | POSITIVE | 1 |
| 19 | FIQRGGGGGGSQEDG | MERCI | POSITIVE | 1 | 66 | GGSKLFYVYYNLFLL | MERCI | POSITIVE | 1 |
| 20 | IQRGGGGGGSQEDGI | MERCI | POSITIVE | 1 | 67 | GSKLFYVYYNLFLLF | MERCI | POSITIVE | 1 |
| 21 | QRGGGGGGSQEDGII | MERCI | POSITIVE | 1 | 68 | SKLFYVYYNLFLLFE | MERCI | POSITIVE | 1 |
| 22 | RGGGGGGSQEDGIII | MERCI | POSITIVE | 1 | 69 | KLFYVYYNLFLLFED | MERCI | POSITIVE | 1 |
| 23 | GGGGGGSQEDGIIIK | MERCI | POSITIVE | 1 | 70 | LFYVYYNLFLLFEDG | MERCI | POSITIVE | 1 |
| 24 | GGGGGSQEDGIIIKK | MERCI | POSITIVE | 1 | 71 | FYVYYNLFLLFEDGG | MERCI | POSITIVE | 1 |
| 25 | GGGGSQEDGIIIKKQ | MERCI | POSITIVE | 1 | 72 | YVYYNLFLLFEDGGG | MERCI | POSITIVE | 1 |
| 26 | GGGSQEDGIIIKKQF | MERCI | POSITIVE | 1 | 73 | VYYNLFLLFEDGGGG | MERCI | POSITIVE | 1 |
| 27 | FILHRGGGGSDQLVF | MERCI | POSITIVE | 1 | 74 | GGGGSLFYVYYNLFL | MERCI | POSITIVE | 1 |
| 28 | ILHRGGGGSDQLVFN | MERCI | POSITIVE | 1 | 75 | GGGSLFYVYYNLFLL | MERCI | POSITIVE | 1 |
| 29 | LHRGGGGSDQLVFNS | MERCI | POSITIVE | 1 | 76 | GGSLFYVYYNLFLLF | MERCI | POSITIVE | 1 |
| 30 | HRGGGGSDQLVFNSI | MERCI | POSITIVE | 1 | 77 | GSLFYVYYNLFLLFE | MERCI | POSITIVE | 1 |
| 31 | RGGGGSDQLVFNSIS | MERCI | POSITIVE | 1 | 78 | SLFYVYYNLFLLFED | MERCI | POSITIVE | 1 |
| 32 | GGGGSDQLVFNSISA | MERCI | POSITIVE | 1 | 79 | LFYVYYNLFLLFEDI | MERCI | POSITIVE | 1 |
| 33 | GGGSDQLVFNSISAR | MERCI | POSITIVE | 1 | 80 | FYVYYNLFLLFEDIG | MERCI | POSITIVE | 1 |
| 34 | GGSDQLVFNSISARA | MERCI | POSITIVE | 1 | 81 | YVYYNLFLLFEDIGG | MERCI | POSITIVE | 1 |
| 35 | GSDQLVFNSISARAL | MERCI | POSITIVE | 1 | 82 | VYYNLFLLFEDIGGG | MERCI | POSITIVE | 1 |
| 36 | SDQLVFNSISARALK | MERCI | POSITIVE | 1 | 83 | SMIDKLFYVYYNLFL | MERCI | POSITIVE | 1 |
| 37 | LKAYGGGGSEDQLVF | MERCI | POSITIVE | 1 | 84 | MIDKLFYVYYNLFLL | MERCI | POSITIVE | 1 |
| 38 | KAYGGGGSEDQLVFN | MERCI | POSITIVE | 1 | 85 | IDKLFYVYYNLFLLG | MERCI | POSITIVE | 1 |
| 39 | AYGGGGSEDQLVFNS | MERCI | POSITIVE | 1 | 86 | DKLFYVYYNLFLLGG | MERCI | POSITIVE | 1 |
| 40 | YGGGGSEDQLVFNSI | MERCI | POSITIVE | 1 | 87 | KLFYVYYNLFLLGGG | MERCI | POSITIVE | 1 |
| 41 | GGGGSEDQLVFNSIS | MERCI | POSITIVE | 1 | 88 | LFYVYYNLFLLGGGG | MERCI | POSITIVE | 1 |

| | | | | |
|---|---|---|---|---|
| 42 | GGGSEDQLVFNSISA | MERCI | POSITIVE | 1 |
| 43 | GGSEDQLVFNSISAR | MERCI | POSITIVE | 1 |
| 44 | GSEDQLVFNSISARA | MERCI | POSITIVE | 1 |
| 45 | SEDQLVFNSISARAL | MERCI | POSITIVE | 1 |
| 46 | EDQLVFNSISARALK | MERCI | POSITIVE | 1 |
| 47 | GGSDKLFYVYYNLFL | MERCI | POSITIVE | 1 |

| | | | | |
|---|---|---|---|---|
| 89 | FYVYYNLFLLGGGGS | MERCI | POSITIVE | 1 |
| 90 | YVYYNLFLLGGGGSG | MERCI | POSITIVE | 1 |
| 91 | VYYNLFLLGGGGSGL | MERCI | POSITIVE | 1 |
| 92 | STRGRKCCRRKKHHH | MERCI | POSITIVE | 1 |
| 93 | TRGRKCCRRKKHHHH | MERCI | POSITIVE | 1 |
| 94 | RGRKCCRRKKHHHHH | MERCI | POSITIVE | 1 |
| 95 | GRKCCRRKKHHHHHH | MERCI | POSITIVE | 2 |

**Supplementary Table S9:** AllergenFP and Vaxijen analysis of MPVs. For the Vaxijen the default threshold is 0.4, and here all the MPVs have scored above 0.4, indicating potential antigenic nature.

| S.No. | MEVs | AllergenFP v.1.0 | Vaxijen | |
|---|---|---|---|---|
| 1 | CTL-MEV | NON-ALLERGEN | ANTIGENS | 0.5740 |
| 2 | HTL-MEV | NON-ALLERGEN | ANTIGENS | 0.4377 |

**Supplementary Table S10:** Analysis of codon-optimized cDNA of all the MEVs.

| S.No. | MPVs | GC content | CAI (Codon Adaptation Index) score | Tandem rare codons |
|---|---|---|---|---|
| 1 | CTL-MEV | 70.00% | 1 | 0% |
| 2 | HTL-MEV | 66.32% | 1 | 0% |
| | Ideal values | 30-70% | 0.8-1.0 | <30% |
